# A molecular switch between mammalian MLL complexes dictates response to Menin-MLL inhibition

**DOI:** 10.1101/2021.10.22.465184

**Authors:** Yadira M. Soto-Feliciano, Francisco J. Sánchez-Rivera, Florian Perner, Douglas W. Barrows, Edward R. Kastenhuber, Yu-Jui Ho, Thomas Carroll, Yijun Xiong, Disha Anand, Alexey Soshnev, Leah Gates, Mary Clare Beytagh, David Cheon, Shengqing Gu, X. Shirley Liu, Andrei V. Krivtsov, Maximiliano Meneses, Elisa de Stanchina, Richard M. Stone, Scott A. Armstrong, Scott W. Lowe, C. David Allis

## Abstract

The chromatin adaptor Menin interacts with oncogenic fusion proteins encoded by *MLL1*-rearrangements (*MLL1*-r), and small molecules that disrupt these associations are currently in clinical trials for the treatment of leukemia. By integrating chromatin-focused and genome-wide CRISPR screens with genetic, pharmacological, and biochemical approaches in mouse and human systems, we discovered a molecular switch between the MLL1-Menin and MLL3/4-UTX chromatin modifying complexes that dictates response to Menin-MLL inhibitors. We show that MLL1-Menin safeguards leukemia survival by impeding binding of the MLL3/4-UTX complex at a subset of target gene promoters. Disrupting the interaction between Menin and MLL1 leads to UTX-dependent transcriptional activation of a tumor suppressor gene-program that is crucial for a therapeutic response in murine and human leukemia. We establish the therapeutic relevance of this mechanism by showing that CDK4/6 inhibitors allow re-activation of this tumor-suppressor program in Menin-inhibitor insensitive leukemia cells, mitigating treatment resistance. The discovery of a molecular switch between MLL1-Menin and MLL3/4-UTX complexes on chromatin sheds light on novel functions of these evolutionary conserved epigenetic mediators and is particularly relevant to understand and target molecular pathways determining response and resistance in ongoing phase 1/2 clinical trials.

## INTRODUCTION

Menin is an evolutionarily conserved nuclear factor that associates with chromatin to recruit (adapt) interacting proteins(1). These include the Trithorax (*Trx*)-related MLL1 (KMT2A) and MLL2 (KMT2B) histone methyltransferase complexes(2, 3), MLL1 oncogenic fusion proteins(4), transcription factors (e.g., c-MYC(5), JUND(6, 7), SMADs(8, 9)), and other chromatin-bound proteins (e.g. LEDGF(10) (reviewed in ref. (11)).

Menin is a core subunit of the MLL1 (ref. (12)) and MLL2 complexes(2), and is responsible for targeting these to chromatin(3). Menin is required for MLL1/MLL2-dependent H3K4 trimethylation of *HOX* genes and their stable long-term expression during development(2, 13). Menin has context-specific functions in human diseases, acting as a tumor suppressor in neuroendocrine malignancies(14, 15) and in certain skin(16), lung(17), and CNS tumors(18), and as an oncogenic co-factor in other cancers, including hepatocellular carcinoma(19) and *MLL1*-rearranged (*MLL1*-r) leukemias(4, 20). Furthermore, over 1000 germline and somatic *MEN1* variants have been identified, some of which are linked to cancer predisposition(21).

Given the pro-oncogenic role of Menin in acute leukemia and other malignancies, small molecule inhibitors targeting the Menin-MLL1 and Menin-MLL2 protein-protein interactions have shown great promise for intercepting and treating different types of cancers(19,22–27). Notably, three structurally different Menin-MLL inhibitors have recently entered clinical trials (NCT04065399, NCT04067336, NCT04811560) and at least one has been granted fast track designation by the FDA for treatment of relapsed/refractory acute leukemias(25,27,28). Thus, an understanding of the molecular mechanisms of action of these drugs would facilitate the development of biomarkers to predict therapeutic response and resistance, and lead to rational design of more effective combination treatments.

## RESULTS

### Functional interplay between MLL1-Menin and MLL3/4-UTX chromatin modifying complexes

To understand the dependency of *MLL1*-r leukemias to Menin and identify factors that dictate response and resistance to Menin-MLL inhibitors, we performed a series of CRISPR-based genetic screens. First, we screened a chromatin-focused CRISPR library in Cas9-expressing mouse leukemia cells driven by a human *MLL1-AF9* transgene (hereafter referred to as MLL-AF9 cells)(29) (**Supplementary Fig. 1A-G, Supplementary Table 1**). Library-transduced cells were cultured in media with DMSO (vehicle) or a Menin-MLL inhibitor (MI-503) (ref. (22)) for 12 cell population doublings, followed by screen deconvolution using next-generation sequencing (**Fig. 1A**). We calculated a score for each gene included in the library by assessing the changes in abundance of sgRNAs during the culture period (**Fig. 1B**). Consistent with previous work, sgRNAs targeting known *MLL1*-r leukemia dependencies, including *Dot1/* (refs. (29–32))*, Brd4* (ref. (33)), and *Myb* (ref. (34)) were strongly depleted in both treatments while control sgRNAs remained largely neutral (**Supplementary Table 2**).

**Figure 1.**
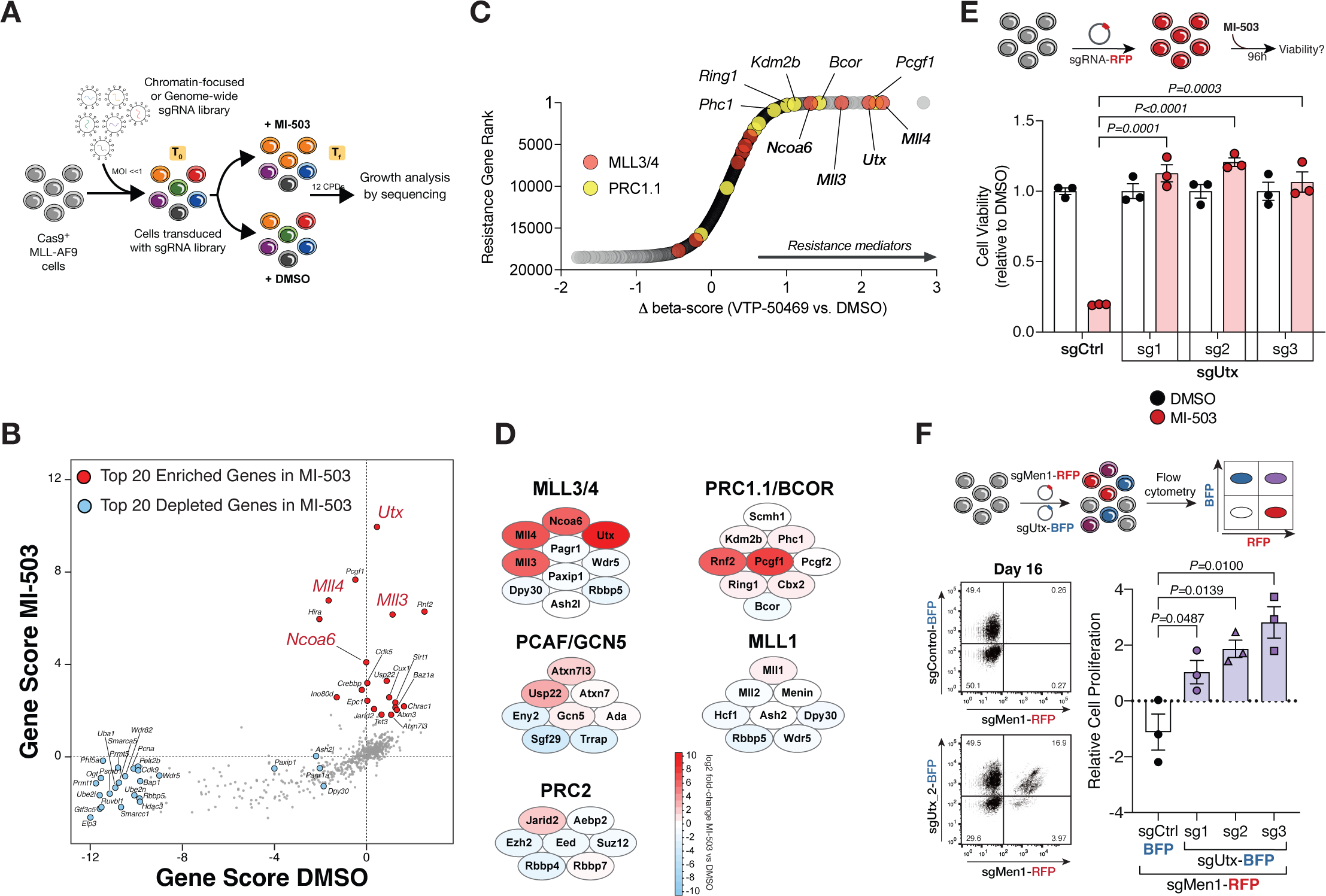
CRISPR screens uncover functional interplay between the mammalian MLL1/2 and MLL3/4 chromatin modifying complexes. **(A)** CRISPR-Cas9-based screening strategy to identify regulators of response to Menin-MLL inhibition. CPD, cell population doublings; sgRNA, single guide RNA; T_f_, final timepoint; T_0_, initial timepoint. **(B)** Chromatin-focused CRISPR screening data showing the top 20 most significantly enriching (red) and depleting (blue) genes in the Menin-MLL inhibitor (MI-503) treatment relative to vehicle (DMSO). Gene scores are shown as the mean log2 fold-change in abundance of the 6 sgRNAs targeting each gene in each condition. **(C)** Genome-wide CRISPR screening data showing gene-level ranking based on differential enrichment of sgRNAs in the Menin-MLL inhibitor treatment (VTP-50469) relative to vehicle (DMSO). Differential (Δ) beta-score between VTP-50469 and DMSO conditions was calculated using MaGeCK. Red circles denote MLL3/4-UTX complex subunits. Yellow circles denote PRC1.1 complex subunits. **(D)** Schematic representation of the top scoring chromatin regulators in the chromatin-focused MI-503 screen and their corresponding protein complexes. Red denotes enriching subunits. Blue denotes depleting subunits. Color scale represents the log_2_ fold-change in abundance of the 6 sgRNAs targeting each subunit in the Menin-MLL inhibitor (MI-503) treatment relative to vehicle (DMSO). **(E)** Viability assay from cells treated with vehicle (DMSO, black) or Menin-MLL inhibitor (MI-503, red) for 96 hours (mean±SEM, n=3 infection replicates, *P*-value calculated by Student’s t-test). sgCtrl, control sgRNA targeting a non-genic region on chromosome 8. **(F)** Differential fitness is shown as the relative fitness of double positive cells (sgMen1-RFP + sgUtx-BFP or sgMen1-RFP + sgCtrl-BFP) to single positive cells (sgMen1-RFP) 16 days post-infection measured by flow cytometry (mean±SEM, n=3 infection replicates, *P*-value calculated by Student’s t-test).

We observed that sgRNAs targeting the histone H3 lysine 27 (H3K27) demethylase *Utx* (*Kdm6a*) and the histone H3 lysine 4 (H3K4) mono-methyltransferases *M//3* (*Kmt2c*) and *M//4* (*Kmt2d*) were the most significantly enriched in the Menin-MLL inhibitor context (**Fig. 1B**, red circles). This result was unexpected given previously described canonical functions for the mammalian MLL complexes(35). For example, the MLL1/2-Menin complex (disrupted by MI-503) is known to catalyze chromatin modifications associated with transcription at *promoters*, including di- or tri-methylation of H3K4 (H3K4me2/3) (13,36,37). On the other hand, the MLL3/4-UTX complex has been shown to regulate *enhancer* states by serving as the major H3K4 mono-methyltransferase (H3K4me1) (refs. (38–42)).

To assess whether these results were idiosyncratic of the cell line, library, or inhibitor used, we performed a genome-wide CRISPR screen in an independently derived MLL-AF9 mouse cell line using VTP-50469, a more potent, selective, and orally bioavailable Menin-MLL inhibitor (refs. (25,27,28)). sgRNAs targeting *Utx*, *M//3*, and *M//4* were also among the most significantly enriched candidate genes identified in this genome-wide screen (**Fig. 1C, Supplementary Fig. 2A-C, Supplementary Table 2**), while shared subunits between the two types of MLL complexes scored similarly in both vehicle and Menin-MLL inhibitor conditions (**Fig. 1B-D, Supplementary Fig. 2D**). These results suggest that core subunits of the MLL3/4-UTX complex(43) modulate the therapeutic response of leukemia cells to Menin-MLL inhibition, pointing to a previously unknown functional cross-talk between these chromatin modifying complexes.

To determine the effects of MLL3/4-UTX loss-of-function in leukemia cell proliferation, we performed *in vitro* growth competition assays (**Supplementary Fig. 3A**). We found that *Utx* or *M//3* inactivation by CRISPR did not have a significant impact on leukemia cell proliferation, but *M//4* inactivation decreased leukemia cell growth (44) (**Supplementary Fig. 3B-D**). Consistent with our genetic screening results, *Utx* disruption using three distinct sgRNAs significantly increased the viability of MI-503-treated leukemia cells (**Fig. 1E, Supplementary Fig. 3E**). In addition, *M//3-* or *M//4*-deficient leukemia cells treated with MI-503 exhibited a proliferative advantage over wild-type cells under drug treatment (**Supplementary Fig. 3F-G**). These orthogonal results establish the MLL3/4-UTX complex as a central modulator of therapy response to Menin-MLL inhibition in acute leukemia.

Since UTX was the most significantly enriched chromatin factor in our screen and this protein is shared by both MLL3 and MLL4 complexes(42,45,46), we focused on UTX disruption to probe the molecular mechanisms linked to resistance to Menin-MLL inhibition. We first tested whether genetic *Utx* inactivation could rescue the effects of Menin-specific ablation in MLL-AF9 leukemia cells. CRISPR-mediated deletion of *Men1* led to robust inhibition of proliferation (**Supplementary Fig. 4A**), but co-deletion of *Utx* suppressed this phenotype, such that *Men1*-deficient MLL-AF9 cells were able to proliferate (**Fig. 1F, Supplementary Fig. 4B-E**). These results establish a previously unknown epistatic relationship between *Men1* and *Utx* in acute leukemia.

To determine if the genetic interaction between Menin and UTX depends on the MLL-fusion protein (MLL-FP) present in these mouse leukemia cells, we ablated the human *MLL1-AF9* oncogenic transgene used to generate this model(47). We targeted the 5 -end of the human *MLL1* gene(48, 49) with CRISPR (48, 49) survival of leukemia cells depend on the sustained presence of the MLL-AF9 fusion protein(20) (**Supplementary Fig. 5A**). We also found that leukemia cells lacking both *MLL-AF9* and *Utx* exhibited proliferation defects similar to cells only lacking *MLL-AF9* (**Supplementary Fig. 5B-C**). These results are consistent with the established dependency of leukemia cells to the pleiotropic gene regulatory activities of MLL1 oncogenic fusions(20) and suggest that the epistatic relationship between Menin and UTX is independent from these activities. Accordingly, co-deletion of *UTX* and *MEN1* in a Menin-dependent non *MLL1*-r human leukemia cell line(50, 51) bypassed the proliferation defects associated with loss of *MEN1* alone (**Supplementary Fig. 5D-E**). Therefore, Menin-MLL inhibitors act, in part, through an evolutionary conserved pathway that involves a functional crosstalk between Menin and UTX, and is downstream or independent of MLL-FPs.

Menin is required for expression of canonical MLL-FP target genes like *Meis1*, which is required for leukemia maintenance(25,36,37,52,53). Moreover, *Meis1* over-expression was recently shown to partially rescue the leukemic stem cell transcriptional program suppressed by Menin-MLL inhibition(27). To determine if the genetic interaction between Menin and UTX regulates expression of these MLL-FP targets, we first treated mouse MLL-AF9 leukemia cells with MI-503 and confirmed that it leads to robust downregulation of *Meis1* (**Supplementary Fig. 5F**). However, to our surprise, cells genetically deficient for both *Men1* and *Utx* showed a similar reduction in *Meis1* levels (**Supplementary Fig. 5G**), yet were able to proliferate (**Fig. 1F, Supplementary Fig. 4E**). These data support a genetic epistasis model between Menin and UTX in Menin-dependent mammalian cells, and suggest that factors beyond *Meis1* can sustain the proliferative capacity of *MLL1*-r and non *MLL1*-r leukemia cells.

### MLL1-Menin complex restricts chromatin occupancy of MLL3/4-UTX at target gene promoters

Given previous work suggesting a primary role for Menin at promoters(6, 54) and UTX at enhancers(42), we performed chromatin immunoprecipitation followed by sequencing (ChIP-Seq) to examine the genome-wide binding patterns of their respective protein complexes. Menin showed strong enrichment at promoter regions (here defined as transcription start sites (TSSs) ± 2kb), which was decreased when its canonical interactions with MLL1/2 and MLL-FPs were disrupted by MI-503 (**Fig. 2A**). Genome-wide enrichment of Menin was also decreased, as evidenced by the reduction in the number of ChIP peaks post MI-503 treatment (**Supplementary Fig. 6A**). We observed a small fraction of UTX at TSSs under basal conditions however, MI-503 treatment led to a ∼5-fold enrichment of UTX at promoters (**Fig. 2B**, **Supplementary Fig. 6A**). We also confirmed that the MI-503-dependent enrichment of UTX on chromatin was not simply the result of increased *Utx* expression or UTX protein stability (**Fig. 2C-D**). These data show that disruption of the Menin-MLL1 interaction leads to dynamic recruitment of UTX to promoter regions (**Fig. 2B**), implying a previously unrecognized functional role for the MLL3/4-UTX complex in promoter regulation and gene transcription in leukemia.

**Figure 2.**
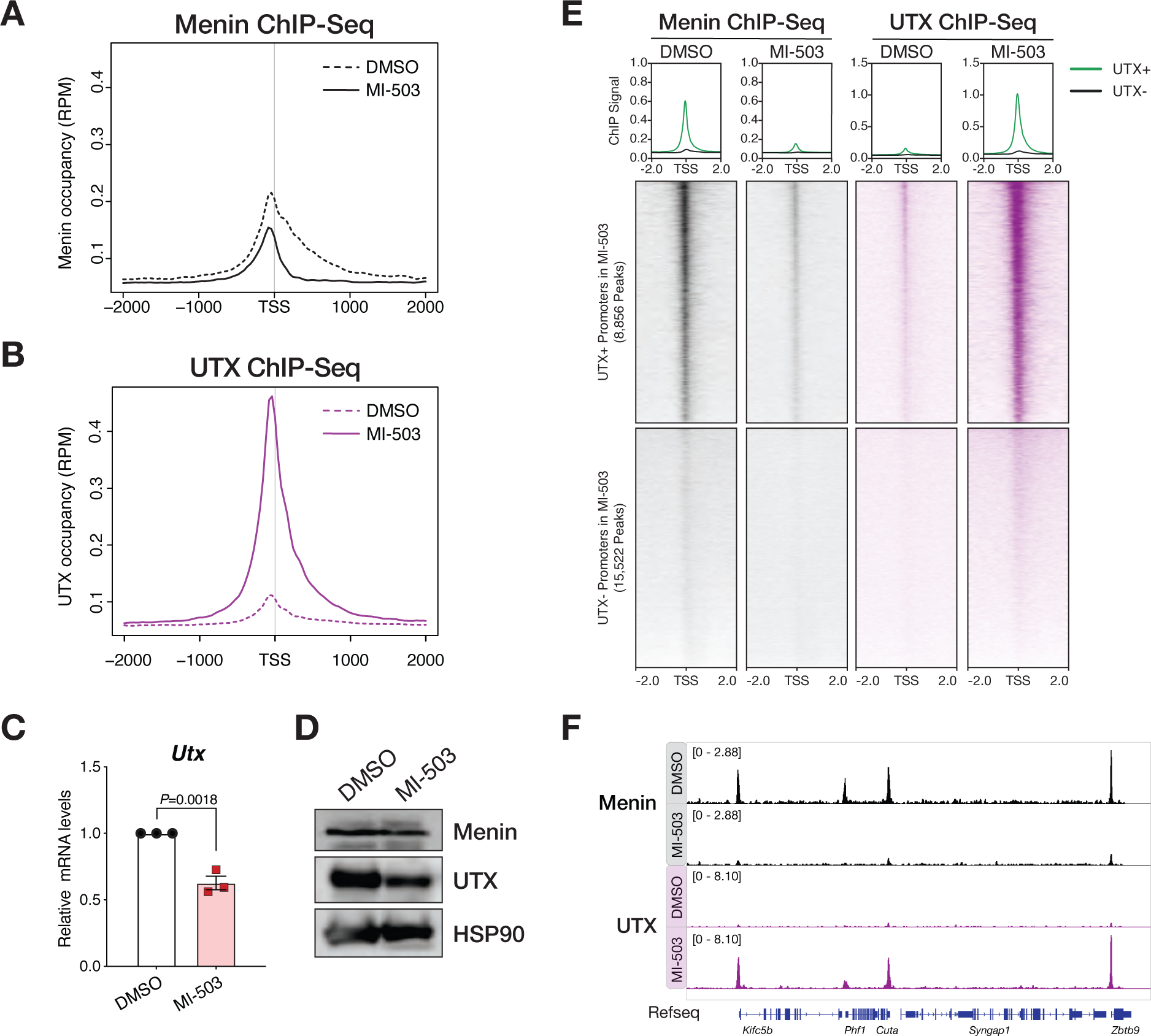
MLL1-Menin complex restricts chromatin occupancy of MLL3/4-UTX at promoters of target genes. **(A)** Metagene analysis showing the average chromatin occupancy of Menin at transcription start sites (TSS), and a region 2000bp downstream and upstream of the TSS. Signals corresponding to cells treated with Menin-MLL inhibitor (MI-503, solid) compared to cells treated with vehicle (DMSO, dotted) for 96 hours are shown. RPM, reads per million. **(B)** Metagene analysis showing the average chromatin occupancy of UTX at transcription start sites (TSS), and a region 2000bp downstream and upstream of the TSS. Signals corresponding to cells treated with Menin-MLL inhibitor (MI-503, solid) compared to cells treated with vehicle (DMSO, dotted) for 96 hours are shown. RPM, reads per million. **(C)** Relative *Utx* mRNA levels determined by qPCR in mouse MLL-AF9 leukemia cells treated with Menin-MLL inhibitor (MI-503, red) compared to vehicle (DMSO, black) for 96 hours (mean ± SEM, n=3 replicates, *P*-value calculated by Student’s t-test). **(D)** Immunoblot analysis of Menin, UTX, and HSP90 proteins (loading control) upon Menin-MLL inhibitor (MI-503) treatment of mouse MLL-AF9 leukemia cells for 96 hours. **(E)** Heatmaps displaying Menin (black) and UTX (purple) ChIP-Seq signals mapping to a 4kb window around TSS. Data is shown for DMSO and MI-503-treated cells for 96 hours. Metagene plot shows the average ChIP-Seq signal for Menin or UTX at promoters that are UTX+ (green) or UTX- (black) post MI-503 treatment. **(F)** Genome browser representation of ChIP-Seq normalized reads for Menin (black) and UTX (purple) in mouse MLL-AF9 leukemia cells treated with either vehicle (DMSO) or Menin-MLL inhibitor (MI-503) for 96 hours.

Intriguingly, promoter regions that became uniquely occupied by UTX significantly overlapped with those where Menin was lost, suggesting a dynamic molecular mechanism between these complexes at a subset of gene promoters (**Fig. 2E-F**, **Supplementary Fig. 6B**). Correlation analysis also suggested that reduction of Menin binding by MI-503 coincided with increased UTX chromatin occupancy at the same genomic loci (**Supplementary Fig. 6C**). Treatment of mouse fibroblasts MEFs (that are not dependent on Menin) with MI-503 did not force UTX mobilization or binding to promoters, indicating that this molecular switch was specific to Menin-dependent cells (**Supplementary Fig. 7A-E**).

The above results indicate that Menin inhibition displaces Menin-MLL1 transcriptional regulatory complexes from promoters, enabling UTX to bind and potentially regulate target gene expression (**Fig. 2**). In agreement, analysis of the chromatin-binding profiles of their cognate H3K4 methyltransferases and their enzymatic histone modifications (**Supplementary Fig. 8A-B**) revealed that Menin-MLL1 inhibition led to a significant decrease in MLL1 chromatin enrichment and a concomitant decrease of its enzymatic product, H3K4me3 (**Supplementary Fig. 8C-E**). In contrast, Menin-MLL1 inhibition led to increased enrichment of the MLL3/4 enzymes at promoter regions co-occupied by UTX and a concomitant increase in H3K4me1 signal at these same loci (**Supplementary Fig. 8C and 8F-G**). Of note, these loci are distinct from those known to be bound and regulated by the MLL-FPs(30) (**Supplementary Fig. 9A-C**). These data indicate that disruption of the MLL1-Menin interaction induces targeting of the core enzymatic subunits of the MLL3/4-UTX complex to non-canonical sites that are normally bound by the MLL1-Menin complex.

Promoter-associated H3K4me1 has been shown to facilitate transcriptional repression in other cellular settings(55), suggesting that deposition of H3K4me1 at gene promoters could depend on context(56). To determine the functional implications of the Menin-MLL inhibitor-induced colocalization of UTX, MLL3/4, and H3K4me1 at target gene promoters, we leveraged the H3 lysine-4-to-methionine (K4M) ’oncohistone mutant as an orthogonal molecular tool to destabilize and alter the function of the MLL3/4-UTX complex(57) (**Supplementary Fig. 10A**). Expression of H3.1K4M in MLL-AF9 leukemia cells led to a proliferative advantage only in the context of Menin-MLL inhibition (**Supplementary Fig. 10B-D**), demonstrating that destabilization of the MLL3/4-UTX complex (**Supplementary Fig. 10E**) can phenocopy the intrinsic resistance of *Utx*-, *M//3*-, or *M//4*-deficient cells to Menin-MLL inhibition (**Fig. 1E, Supplementary Fig. 3E-G**). These results further support a model whereby the MLL3/4-UTX complex serves as a context-specific and central modulator of therapy response to Menin-MLL inhibition in leukemia cells.

### NF-YA confers genomic specificity to a Menin-UTX molecular switch on chromatin

Since Menin lacks a defined DNA binding domain(58), we tested whether a sequence-specific DNA binding factor may regulate the switch between MLL1-Menin and MLL3/4-UTX occupancy at specific promoters. Motif analysis on the genomic regions bound by Menin and UTX in leukemia cells and fibroblasts revealed that NF-Y sequence motifs were the most significantly enriched (59) in leukemia cells (*P* = 1e^-130^) (**Fig. 3A**) and selective to leukemia cells (**Supplementary Fig. 11A-B**). In agreement, ChIP-Seq analysis for NF-YA (the DNA binding and transactivation subunit of the NF-Y complex(59)) showed that it co-occupies sites reciprocally bound by Menin and UTX (**Fig. 3B, Supplementary Fig. 11C**). The sites bound by UTX in the context of Menin-MLL inhibition also had lower NF-YA chromatin enrichment, an effect that was not due to decreased protein levels (**Fig. 3B-C, Supplementary Fig. 11C**).

**Figure 3.**
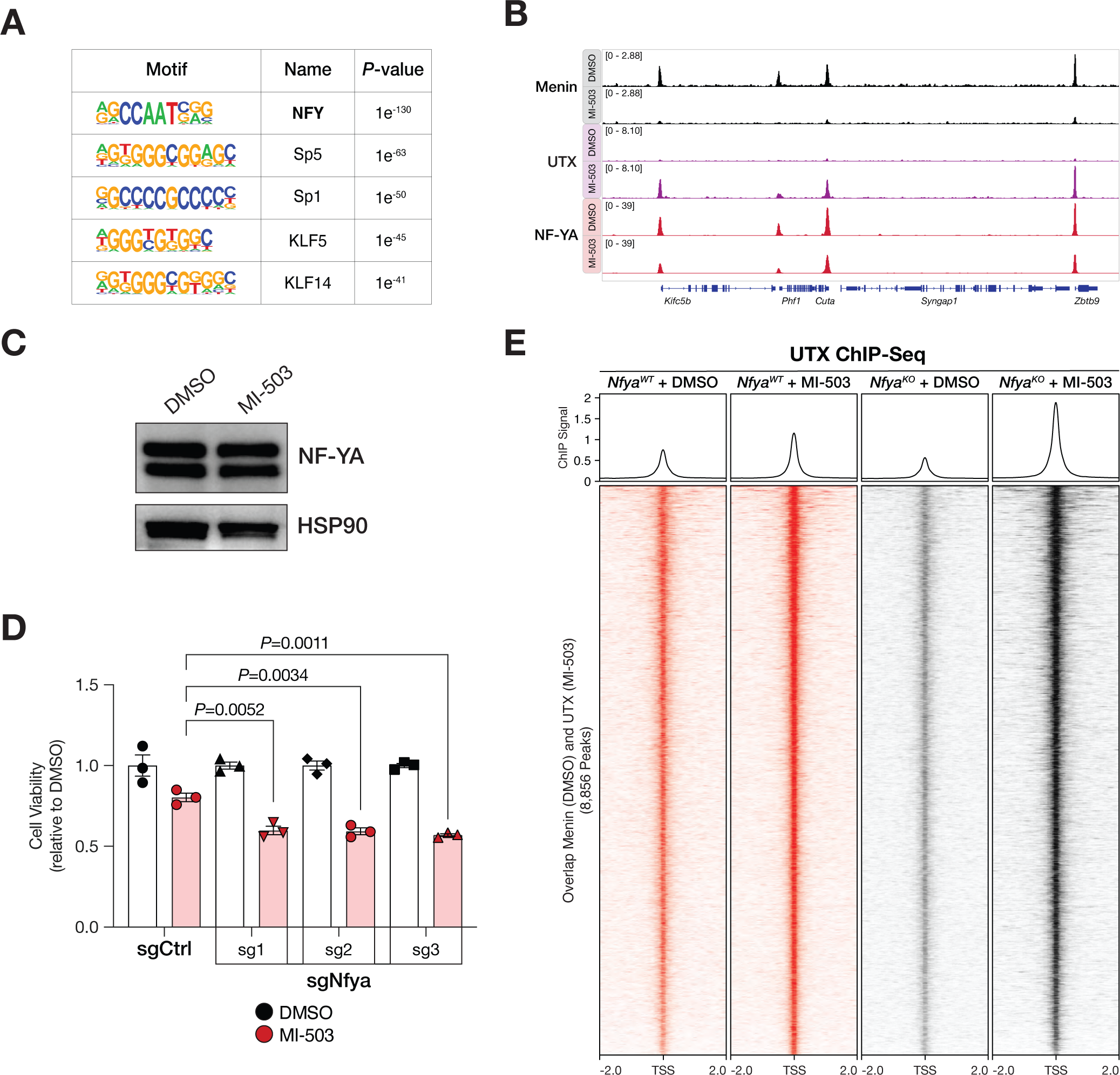
NF-YA contributes to genomic specificity of the Menin-UTX molecular switch on chromatin. **(A)** HOMER *de novo* motif analysis of overlapping ChIP-Seq peaks between Menin (in DMSO) and UTX (in MI-503) in mouse MLL-AF9 leukemia cells. **(B)** Genome browser representation of ChIP-Seq normalized reads (average RPKM) for representative loci bound by Menin (black), UTX (purple), and NF-YA (red) in cells treated with vehicle (DMSO) or Menin-MLL inhibitor (MI-503) for 96 hours. **(C)** Immunoblot analysis of NF-YA and HSP90 proteins (loading control) upon Menin-MLL inhibitor (MI-503) treatment of mouse MLL-AF9 leukemia cells for 96 hours. **(D)** Viability assay from cells treated with vehicle (DMSO, black) or Menin-MLL inhibitor (MI-503, red) for 96 hours (mean±SEM, n=3 infection replicates, *P*-value calculated by Student’s t-test). sgCtrl, control sgRNA targeting a non-genic region on chromosome 8. **(E)** Heatmaps displaying UTX ChIP-Seq signal mapping to a 4-kb window around TSS in *Nfya*^WT^ (red) or *Nfya*^KO^ (black) mouse MLL-AF9 leukemia cells treated with vehicle (DMSO) or Menin-MLL inhibitor (MI-503) for 96 hours. Metagene plot represents the average UTX ChIP-Seq signal at promoters.

To functionally dissect the relationship between Menin, UTX, and NF-YA, we generated *Nfya*^KO^ MLL-AF9 leukemia cells and treated them with MI-503 (**Supplementary Fig. 12A**). Disruption of *Nfya* decreased the viability of MI-503-treated leukemia cells relative to control cells at two different time points, suggesting a synthetic lethal relationship between *Nfya* and *Men1* (**Fig. 3D, Supplementary Fig. 12B**). Moreover, treatment of *Nfya*^KO^ cells with MI-503 led to a significant increase in UTX occupancy at genomic regions normally bound by Menin and NF-YA at steady-state (**Fig. 3E**, **Supplementary Fig. 12C**). These observations were also consistent with co-essentiality analyses in human cells(60) showing that a subset of *MEN1*-dependent cells are also dependent on members of the NF-Y complex for survival (**Supplementary Fig. 12D-G)**. Collectively, these results implicate NF-Y in restricting UTX occupancy at promoter regions and suggest that Menin binding to these sites is in part mediated by the sequence-specific DNA binding activity of NF-Y.

### Transcriptional co-regulation of tumor suppressive pathways by a Menin-UTX molecular switch

To probe the molecular basis of this switch further, we performed transcriptional profiling of MLL-AF9 leukemia cells treated with MI-503 and identified pathways that are reciprocally regulated by Menin and UTX. While the activity of the MLL1-Menin complex is associated with actively transcribed developmental genes(2, 13), MI-503 treatment resulted in both up- and down-regulation of gene expression, with the majority of significantly upregulated genes reciprocally bound by Menin and UTX (**Fig. 4A-B**).

**Figure 4.**
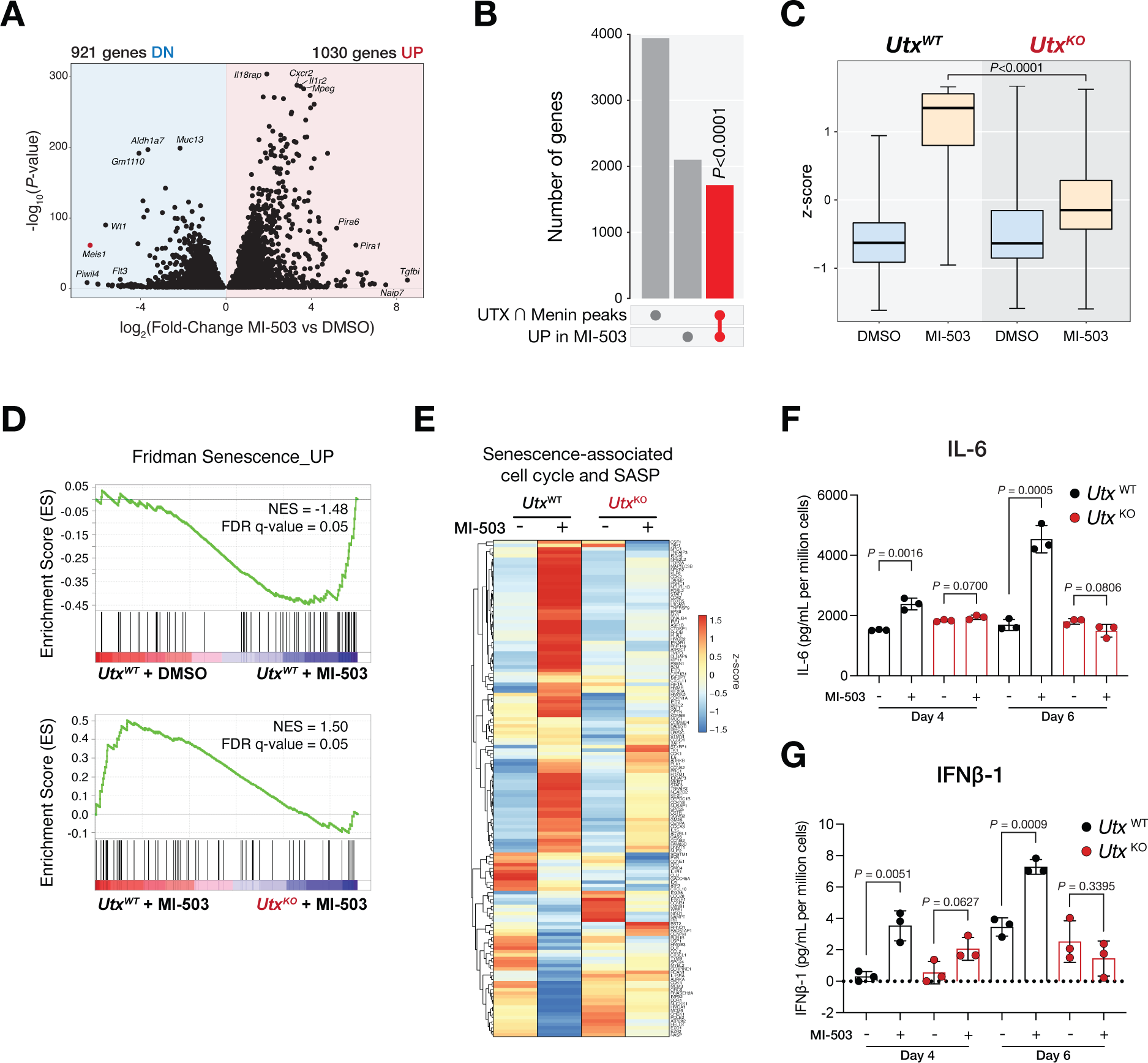
Transcriptional co-regulation of tumor suppressive pathways by the Menin-UTX switch. **(A)** Volcano plot of differentially expressed genes in mouse MLL-AF9 leukemia cells treated with Menin inhibitor (MI-503) or vehicle (DMSO) for 96 hours. Significantly (*P*<0.05) downregulated (DN) genes are shown on the left (n=921 genes). Significantly (*P*<0.05) upregulated (UP) genes are shown on the right (n=1030). **(B)** Upset plot showing significant overlap (red) between genes that undergo replacement of Menin by UTX at their promoters and MI-503-induced genes. *P*-value for overlap is shown. **(C)** Boxplot showing expression levels of genes that are induced by Menin-MLL inhibitor (MI-503) treatment and are bound by UTX at their promoters in this condition and by Menin at steady-state. Expression levels are shown for *Utx*^WT^ and *Utx*^KO^ leukemia cells. Midline in boxplots represent median. *P*-value for MI-503 comparison is shown. **(D)** Gene set enrichment analysis (GSEA) showing that Menin-UTX targets induced by Menin-MLL inhibitor (MI-503) are significantly enriched for genes regulating cellular senescence. FDR, false discovery rate; NES, normalized enrichment score. **(E)** Heatmap showing relative gene expression levels of senescence-associated cell cycle and senescence-associated secretory phenotype (SASP) genes in mouse *Utx*^WT^ and *Utx*^KO^ MLL-AF9 leukemia cells treated with Menin-MLL inhibitor (MI-503) or vehicle (DMSO) for 96 hours. **(F)** and **(G)** Secreted levels of IL-6 and IFN β-1 in conditioned-media derived from mouse *Utx*^WT^ (black) and *Utx*^KO^ (red) MLL-AF9 leukemia cells treated with Menin-MLL inhibitor (MI-503) or vehicle (DMSO) for four or six days. Data are quantified as pg/mL of secreted cytokine per million cells Data are quantified as pg/mL of secreted cytokine per million cells (mean±SEM, n=3 replicates, *P*-value calculated by Student’s t-test).

To gain further insights into mechanism, we performed ChIP-Seq against acetylated H3K27 and H4K16 (histone modifications associated with gene activation(61)) and found that the levels of these modifications increased at sites where the MLL3/4-UTX complex is enriched at upon Menin-MLL inhibition (**Supplementary Fig. 13A-B**). These loci also showed increased binding of MOF, a histone H4K16 acetyltransferase(62–64)(**Supplementary Fig. 13C**). Thus, Menin-MLL inhibition produces dynamic changes in gene expression due to loss of MLL1-Menin-dependent repressive activity and a concomitant increase in histone modifications associated with gene activation (including H4K16ac and H3K27ac)(65, 66).

To determine if UTX was necessary for gene activation, we generated *Utx^KO^* MLL-AF9 leukemia cells, treated these with MI-503, and profiled by RNA-Seq (**Supplementary Fig. 14A-B**). We found that genes bound by Menin and UTX that are induced by MI-503 failed to get activated in *Utx^KO^* cells, suggesting UTX function is necessary for their transcriptional activation upon Menin displacement from chromatin (we refer to these genes as ’Menin-UTX targets) (**Fig. 4C**, **Supplementary Table 4**). Consistent with the idea that this mechanism is independent of the MLL-FP, we found that MI-503-treated *Utx^KO^* cells still exhibited downregulation of canonical MLL-AF9 targets and induction of myeloid differentiation programs (**Supplementary Fig. 15A-C**), as has been observed in *Utx^WT^* cells(22,25,27). Moreover, these cells were able to proliferate without re-expression of *Meis1* and *Hoxa9* - two critical Menin-MLL-FP targets(25,27,52,53) (**Fig. 1E**, **Supplementary Fig. 5G, Supplementary Fig. 15D**). These data suggest a new paradigm whereby the effects of Menin-MLL inhibition on MLL-FP target genes are independent of its effects on Menin-UTX transcriptional targets, and that concomitant induction of tumor suppressive gene expression programs and repression of canonical MLL-FP targets is required for the anti-leukemic activity of Menin-MLL1 inhibitors.

To gain insight into cellular and molecular pathways regulated by the Menin-UTX switch, we performed gene ontology analysis(67, 68) of Menin-UTX targets and found significant evidence for their association with transcriptional programs related to proliferation, differentiation, and survival (**Supplementary Fig. 16A, Supplementary Table 4**,). To further evaluate the relevance of these gene ontology terms, we performed Gene Set Enrichment Analysis (GSEA)(69) on transcriptional data from MI-503-treated *Utx^WT^* and *Utx^KO^* cells and observed a significant correlation between the presence of UTX and the expression of senescence-associated genes following treatment(70) (**Fig. 4D**, **Supplementary Fig. 16B-G**, **Supplementary Table 4**). This correlation was also observed when we analyzed a curated list of genes involved in cell cycle arrest and therapy-induced senescence(71)(**Fig. 4E**). These transcriptional changes are consistent with the fact that Menin-MLL inhibition induces a combination of cell cycle arrest, apoptosis, and differentiation (**Supplementary Fig. 1E-G**)(22), as well as our observation that the senescence-associated H4K16ac modification increases with MI-503 treatment(72) (**Supplementary Fig. 13B-C**). To gain additional insights into senescence-associated gene activation mechanisms, we performed ChIP-Seq for RNA Polymerase II and observed increased occupancy at the promoters of senescence-associated genes that are up-regulated by MI-503 treatment(**Supplementary Fig. 16H**). Thus, the interplay between Menin and UTX in the context of Menin-MLL inhibition regulates the expression of genes involved in cell cycle arrest and senescence.

The cellular senescence program is highly complex and characterized in part by induction of permanent cell cycle arrest and a senescence-associated secretory phenotype (SASP)(73). To determine if Menin-MLL inhibition induces the SASP, we measured cytokine and chemokine levels in conditioned media from MLL-AF9 leukemia cells treated with MI-503. Secretion of prototypical SASP cytokines like IL-6, IFNβ-1, IL-3, and IL-15 was induced upon Menin-MLL inhibition in a *Utx*-dependent manner, supporting a direct role for the Menin-UTX switch in regulating cellular senescence(**Fig. 4F-G, Supplementary Fig. 17A-C)**. Thus, Menin-MLL inhibition engages a tumor suppressive network that includes therapy-induced cellular senescence and is regulated by the MLL3/4-UTX complex.

### The enzymatic activity of UTX is dispensable for tumor suppressive functions in response to Menin-MLL inhibition

The UTX protein contains several functional domains, including a JmjC demethylase domain that catalyzes the removal of the H3K27me3 histone mark(74–77). To determine the regions of UTX that are necessary for treatment-associated UTX-dependent phenotypes, we performed structure-function-rescue experiments in *Utx*^KO^-null leukemia cells using lentiviral constructs encoding dual N-terminal HA- and C-terminal Flag-tagged truncations of UTX (**Fig. 5A, Supplementary Fig. 18A-C)**. Full-length UTX, or a UTX truncation harboring the first 500 amino acids (UTX^1-500^) and lacking the JmjC demethylase domain(74–77), were sufficient to re-sensitize cells to MI-503 treatment while truncations proximal to the C-terminus were unable to do so (**Fig. 5B**). Consistent with these cellular phenotypes, full-length UTX and UTX^1-500^ were sufficient to rescue UTX-dependent transcriptional phenotypes associated with Menin-MLL inhibition (**Fig. 5C**), including induction of Menin-UTX target genes (**Fig. 5D, Supplementary Fig. 19A**). These results demonstrate that an N-terminal truncation of UTX, lacking the JmjC demethylase domain and a recently reported intrinsically disordered region(78), is both necessary and sufficient to drive treatment-associated tumor suppressive responses.

**Figure 5.**
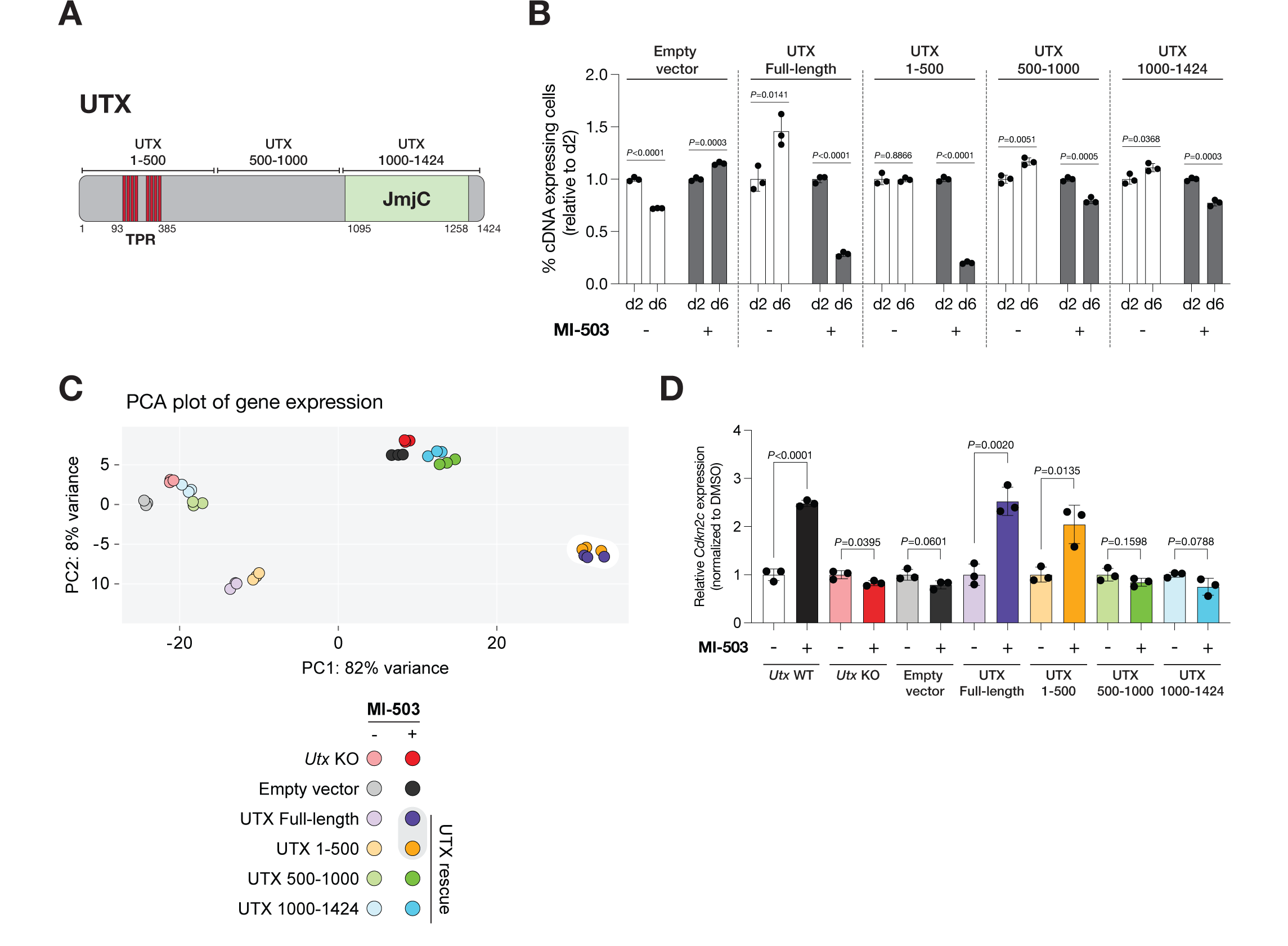
Enzymatic activity of UTX is dispensable for its tumor suppressive functions in response to Menin-MLL inhibition. **(A)** Schematic of UTX protein. Highlighted are the 8 tetratricopeptide repeats (TPR) (93-385aa) and the histone demethylase (JmjC) domain (1095-1258aa). Three ∼500 amino acid long truncations are also represented. **(B)** Growth competition assay in mouse *Utx*^KO^ MLL-AF9 leukemia cells expressing different RFP-tagged *Utx* cDNAs and treated with Menin-MLL inhibitor (MI-503) for 2 or 6 days. Graph shows the relative growth of leukemia cells infected with RFP-tagged *Utx* cDNAs measured by flow cytometry (mean±SEM, n=3 infection replicates, *P*-value calculated by Student’s t-test). **(C)** Principal component analysis (PCA) of gene expression data from *Utx*^KO^ MLL-AF9 leukemia cells expressing different RFP-tagged *Utx* cDNAs and treated with vehicle (DMSO) or Menin-MLL inhibitor (MI-503) for 96 hours. **(D)** *Cdkn2c* expression (mean normalized read counts) from different *Utx* truncations in mouse MLL-AF9 leukemia and treated with vehicle (DMSO) or Menin-MLL (MI-503) for 96 hours (mean±SEM, n=3 replicates, *P*-value calculated by Student’s t-test).

To further confirm that UTX employs non-catalytic mechanisms to regulate gene expression in the context of Menin-MLL inhibition, we performed ChIP-Seq for H3K27me3, a modification catalyzed by PRC2 (ref. (79)) and removed by UTX (refs. (74–77)). We found that the genomic redistribution of UTX induced upon treatment with MI-503 (**Fig. 2B and 2E**) did not affect global levels or distribution of H3K27me3 (**Supplementary Fig. 20A-B**), suggesting that this histone modification might not play a critical role in these phenotypes. To functionally test this possibility, we generated isogenic *Ezh2^KO^ Utx^WT^* and *Ezh2^KO^ Utx^KO^* MLL-AF9 leukemia cells and confirmed the absence of H3K27me3 by immunoblotting (**Supplementary Fig. 20C**). Consistent with our model, *Ezh2^KO^ Utx^WT^*leukemia cells remained exquisitely sensitive to MI-503 while *Ezh2^KO^ Utx^KO^* cells were resistant to Menin-MLL inhibition (**Supplementary Fig. 20D**). These results demonstrate that the catalytic function of UTX is dispensable for UTX-dependent phenotypes and therapeutic responses to Menin-MLL inhibitors.

### Combinatorial targeting of Menin and CDK4/6 overcomes resistance associated with MLL3/4-UTX dysfunction

Senescence can be regulated at both transcriptional and post-transcriptional levels, and chromatin regulation has been functionally implicated in modulating these programs(72,80,81). Thus,, we examined whether the Menin-UTX molecular switch directly regulates the expression of cell cycle arrest- and senescence-associated genes by direct binding to their promoters(82, 83). Consistent with this model, we found that Menin is bound to the promoters of the cyclin-dependent kinase (CDK) inhibitors *Cdkn2cllnk4c* and *Cdkn2dllnk4d* at basal conditions(84), and that enrichment is decreased upon Menin-MLL inhibition, coinciding with their increased expression (**Fig. 6A, Supplementary Fig. 21A**). Conversely, we found that UTX binds to these promoters only in the context of Menin-MLL inhibition, leading to UTX-dependent upregulation of both CDK inhibitors (**Fig. 6A-B**). Thus, the Menin-UTX molecular switch regulates the expression of these CDK inhibitors by direct chromatin regulation.

**Figure 6.**
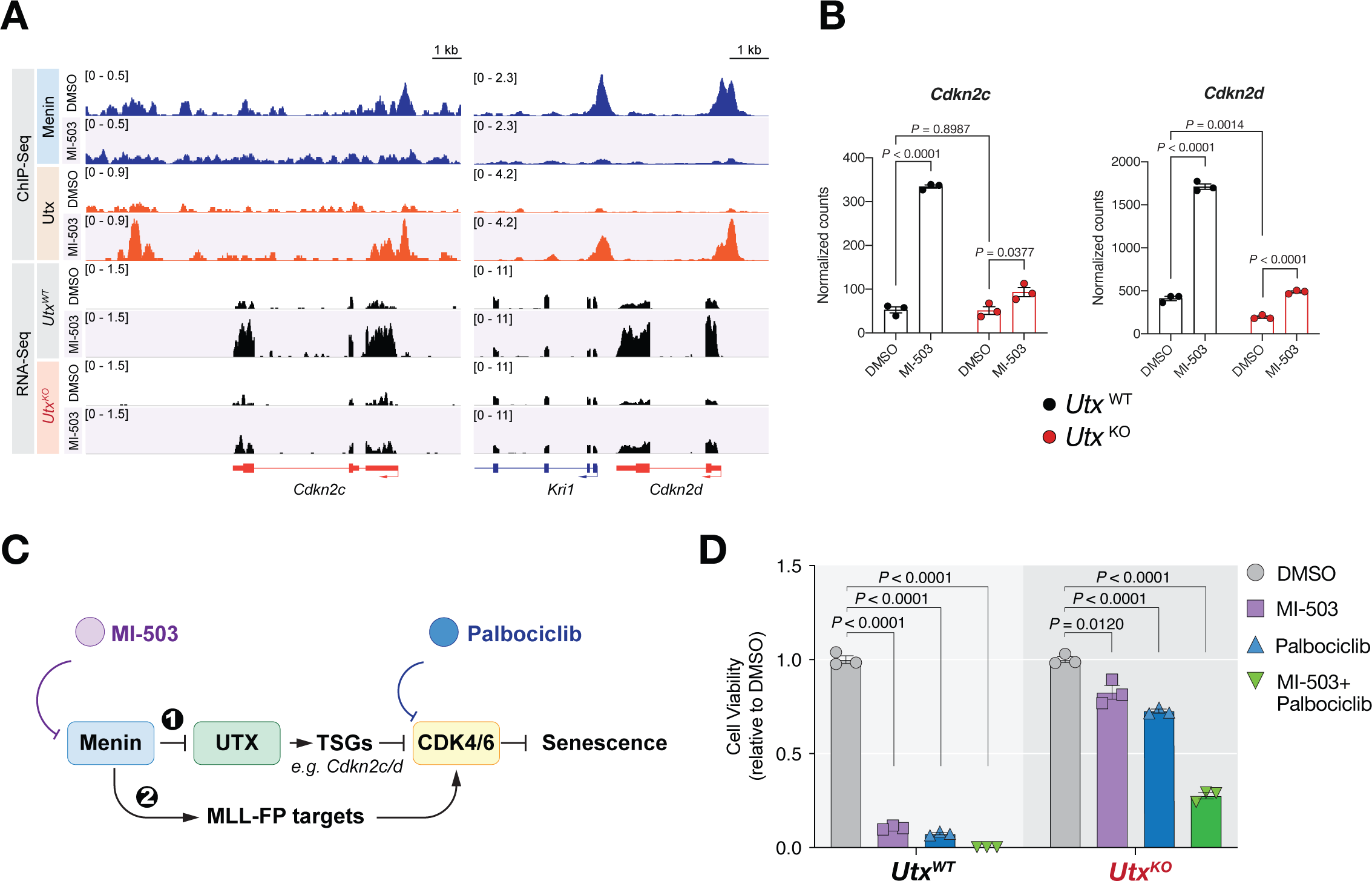
Combinatorial targeting of Menin and CDK4/6 overcomes resistance associated with MLL3/4 dysfunction. **(A)** Genome browser representation of ChIP-Seq (top) and RNA-Seq (bottom) normalized reads (average RPKM) for *Cdkn2c* and *Cdkn2d* loci from mouse *Utx*^WT^ or *Utx*^KO^ MLL-AF9 leukemia cells treated with vehicle (DMSO) or Menin-MLL inhibitor (MI-503) for 96 hours. **(B)** *Cdkn2c* and *Cdkn2d* expression (mean normalized read counts) from mouse *Utx*^WT^ (black) and *Utx*^KO^ (red) MLL-AF9 leukemia cells treated with vehicle (DMSO) or Menin-MLL inhibitor (MI-503) for 96 hours (mean±SEM, n=3 replicates, *P*-value calculated by Student’s t-test). **(C)** Proposed model and rationale for combination therapies based on Menin-MLL and CDK4/6 inhibitors. Our data support a model whereby (1) Menin restricts UTX-mediated transcriptional activation of tumor suppressor genes, including *Cdkn2c* and *Cdkn2d*, which are natural inhibitors of the CDK4 and CDK6 kinases, which in turn inhibit cell cycle arrest and senescence. Model predicts that CDK4/6 inhibition using Palbociclib should boost the anti-cancer activity of Menin-MLL inhibitors, which we show induces a MLL3/4-UTX tumor suppressive axis, by more potently inhibiting these downstream kinases. (2) Menin is known to be required for activation of MLL-FP targets like *Meis1* and *Cdk6* itself to sustain leukemia. Our model predicts that combination therapies based on Menin-MLL and CDK4/6 inhibitors should act synergistically to suppress leukemia proliferation by potently engaging two parallel pathways that converge on regulation of cell cycle progression. **(D)** Relative viability of *Utx*^WT^ and *Utx*^KO^ MLL-AF9 leukemia cells treated with either vehicle (DMSO), Menin-MLL inhibitor (MI-503), CDK4/6 inhibitor (Palbociclib), or a combination of both inhibitors for 6 days (mean±SEM, n=3 replicates, *P*-value calculated by Student’s t-test).

Since the proteins encoded by these two genes are natural inhibitors of the CDK4 and CDK6 kinases(84), we tested whether pharmacological inhibition of CDK4/6 could bypass the intrinsic resistance of *Utx^KO^* cells to MI-503 while retaining the therapeutic effects of Menin-MLL inhibition on MLL-FP targets (**Fig. 6C**). Treatment of MLL-AF9 leukemia cells with MI-503 and the FDA-approved CDK4/6 inhibitor Palbociclib(85) Sherr et al. 2016(85) showed that *Utx^KO^* cells were more resistant to either MI-503 or Palbociclib alone relative to *Utx^WT^*cells, likely due to higher basal levels of *Cdk6* transcripts (**Fig. 6D, Supplementary Fig. 21D**) (82, 86). However, combined inhibition of Menin-MLL and CDK4/6 resulted in a synergistic effect (CD=0.4) (ref. (87)) on inhibiting cell proliferation to levels similar to those achieved by MI-503 treatment of *Utx^WT^*cells (**Fig. 5D**). These results demonstrate that targeting pathways regulated by Menin and UTX can produce combinatorial therapeutic effects and suggest that the anti-leukemic effects of MI-503(22) are primarily through reactivation of tumor suppressor pathways and not solely through dampening transcription of MLL-FP targets like *Meis1* (refs. (22,25,27,52)) (**Supplementary Fig. 5G and 15D**). Thus, the combination of Palbociclib with Menin-MLL inhibitors may represent a novel and more effective therapeutic strategy for Menin-dependent cancers(23,24,26,88).

### *In vivo* response to Menin-MLL inhibitors is accompanied by induction of MLL3/4-UTX-dependent tumor suppressive programs

Small molecule inhibitors of the Menin-MLL interaction have shown significant promise in preclinical models of acute lymphoid and myeloid leukemia(25, 89) and are currently in Phase II clinical trials for treatment of patients with acute leukemia (SNDX-5613 (NCT04067336), KO-539 (NCT04065399), and JNJ-75276617 (NCT04811560)).

Notably, SNDX-5613 was recently granted fast track designation by the FDA for treatment of relapsed/refractory acute leukemias. To examine whether the Menin-UTX molecular switch described above is operative in leukemia patients treated with Menin-MLL inhibitors, we performed longitudinal RNA-Seq analysis of primary AML cells derived from two patients with *NPM1c*-mutated (Patient 1) and *MLL*-rearranged (Patient 2) leukemia treated with SNDX-5613 (AUGMENT-101 clinical trial NCT04065399) (**Fig. 7A, Supplementary Fig. 22A**). Consistent with our hypothesis, SNDX-5613 led to concomitant suppression of canonical MLL-FP target genes (e.g., *MElS1*) and induction of CDK inhibitors (e.g., *CDKN2C*) (**Fig. 7B-E, Supplementary Fig. 22B-C**).

**Figure 7.**
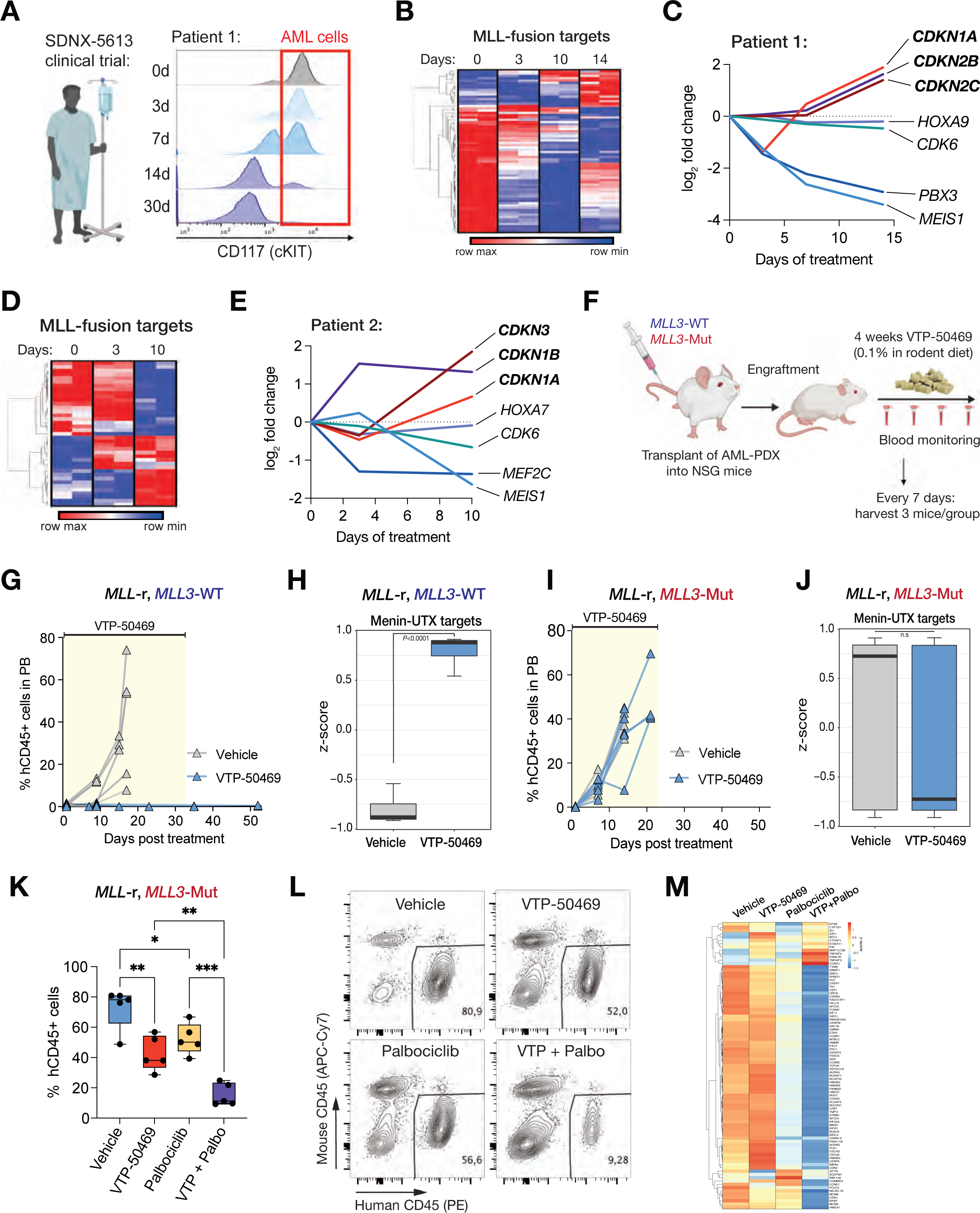
*In vivo* response to Menin-MLL inhibition is accompanied by induction of MLL3/4-UTX-dependent tumor suppressive programs. **(A)** Longitudinal flow cytometry analysis showing the fraction of CD45lo, cKIT+ leukemia cells in the peripheral blood of an *NPM1c* mutant patient (Patient 1) during cycle 1 of Menin-MLL inhibitor (SDNX-5613) treatment as part of the AUGMENT-101 clinical trial (NCT04065399). **(B)** Temporal gene expression changes for MLL-FP targets in FACS-sorted leukemia blasts cells isolated from Patient 1 as part of the AUGMENT-101 clinical trial (NCT04065399). Heatmap shows all MLL-FP targets that are differentially expressed at day 14 vs. day 0 of treatment cycle 1. **(C)** Temporal expression levels of genes involved in cell cycle arrest and senescence (*CDKN1A*, *CDKN2B*, and *CDKN2C*) and MLL-FP targets (*HOXA9*, *CDK6*, *PBX3*, and *MEIS1*) in leukemia blasts cells isolated from Patient 1 treated with SDNX-5613. **(D)** Temporal gene expression changes for MLL-FP targets in FACS-sorted leukemia blasts cells isolated from Patient 2 treated with SDNX-5613 as part of the AUGMENT-101 clinical trial (NCT04065399). Heatmap shows all MLL-FP targets that are differentially expressed at day 10 vs. day 0 of treatment cycle 1. **(E)** Temporal expression levels of genes involved in cell cycle arrest and senescence (*CDKN1A*, *CDKN1B*, and *CDKN3*) and MLL-FP targets (*HOXA7*, *CDK6*, MEF2C, and *MEIS1*) in leukemia blasts cells isolated from Patient 2 treated with SDNX-5613. **(F)** Schematic of *in vivo* treatment experiments using genetically-defined acute myeloid leukemia (AML) patient-derived xenografts (PDXs). Mice were transplanted with either *MLL3*-WT or *MLL3*-Mutant AML PDXs and, upon disease engraftment, were randomized into Menin-MLL inhibitor (VTP-50469, 0.1% in rodent diet) or normal chow for a duration of 4 weeks. Disease progression was monitored weekly by bleeding and AML cells were sorted at 7 days post initiation of treatment using magnetic mouse cell depletion from the bone marrow of animals to perform RNA-Seq. **(G)** Disease progression as measured by the percentage of human CD45+ cells in the peripheral blood (PB) of mice harboring *MLL3*-WT leukemia treated with vehicle (grey) or Menin-MLL inhibitor (VTP-50469, blue). **(H)** Boxplot denoting gene expression changes of Menin-UTX targets in AML cells isolated from mice harboring *MLL3*-WT leukemia treated with vehicle (grey) or Menin-MLL inhibitor (VTP-50469, blue). **(I)** Disease progression as measured by the percentage of human CD45+ cells in the peripheral blood (PB) of mice harboring *MLL3*-mutant leukemia treated with vehicle (grey) or Menin-MLL inhibitor (VTP-50469, blue). **(J)** Boxplot denoting gene expression changes of Menin-UTX targets in AML cells isolated from mice harboring MLL3-mutant leukemia treated with vehicle (grey) or Menin-MLL inhibitor (VTP-50469, blue). **(K)** Leukemia burden in the bone marrow of recipient mice transplanted with the MLL3-mutant AML PDX and treated with Menin-MLL inhibitor (VTP-50469), CDK4/6 inhibitor (Palbociclib) or the combination of these two inhibitors (measured by % of human CD45+ cells). VTP-50469 was administered via drug-supplemented rodent chow (0.1%) for 10 days, Palbociclib was given once daily via intraperitoneal injections (35mg/kg) for 7 days. **(L)** Representative FACS plots showing the abundance of human leukemia cells in recipient mice from each treatment group. **(M)** Heatmap denoting changes in cell cycle-associated gene expression signatures in FACS-sorted human leukemia cells isolated from recipient mice transplanted with the MLL3-mutant PDX and treated with VTP-50469, Palbociclib, or the combination of these two inhibitors.

As an orthogonal approach, we analyzed the *in vivo* response and resistance of *MLL*-rearranged acute leukemia patient-derived xenografts (PDXs) to VTP-50469 (a close analog of SNDX-5613)(25) (**Fig. 7F, Supplementary Fig. 23A**). Consistent with our model, mice transplanted with an *MLL3* wild type PDX showed a potent therapeutic response to VTP-50469 and transcriptional induction of Menin-UTX targets (**Fig. 7G-H, Supplementary Fig. 24A**). Conversely, an *MLL3* mutant PDX exhibited primary resistance to VTP-50469 and failed to induce this gene expression program (**Fig. 7I-J, Supplementary Fig. 23B and 24A**), linking the induction of Menin-UTX-targets to preclinical drug response. Thus, gene expression programs regulated by MLL1-Menin and MLL3/4-UTX complexes are operative in AML patients and PDX models treated with orally bioavailable Menin-MLL inhibitors that are currently under clinical investigation.

To test whether CDK4/6 inhibition can overcome the blunted induction of endogenous CDK inhibitors in the context of Menin inhibitor resistance, we treated mice harboring the *MLL3*-mutant AML PDX with VTP-50469 and Palbociclib (**Fig. 7K**). While single treatment with Menin or CDK4/6 inhibitors led to a minor decrease in leukemia burden in the bone marrow of recipient mice after 10 days of treatment, combination treatment with VTP-50469 and Palbociclib induced significant leukemia regression (**Fig. 7K-L**). Moreover, RNA-Seq from human leukemia cells isolated from these animals showed induction of cell cycle arrest and senescence-associated gene expression signatures in mice treated with both VTP-50469 and Palbociclib (**Fig. 7M, Supplementary Fig. 24B-D**). Altogether, this data provides pre-clinical evidence for the feasibility and efficacy of combined Menin- and CDK4/6-inhibition to overcome the blunted induction of senescence-associated programs in patient-derived leukemias resistant to Menin-inhibitor monotherapy.

## DISCUSSION

The chromatin adaptor protein Menin exhibits context-specific functions in different tissues, acting as a tumor suppressor gene in neuroendocrine(14, 15), lung(17, 90), skin(16), and CNS malignancies(18), and as an oncogenic cofactor in hepatocellular(19)and hematologic cancers(4, 20). Given that Menin can interact with similar cofactors in disparate settings and the biological and molecular basis for these ostensibly paradoxical findings has remained unclear. For example, Menin functionally cooperates with MLL proteins to activate transcription of the *Cdkn1blp27^Kip1^*and *Cdkn2dllnk4d* CDK inhibitors as a tumor suppressive mechanism in neuroendocrine tumors and lung cancer(3,17,91,92), yet the same protein-protein interaction is critical for leukemia maintenance(4, 10).

Our study sheds light on these paradoxical observations by revealing a functional interaction between the mammalian histone methyltransferase complexes MLL1-Menin and MLL3/4-UTX and, in doing so, challenges the paradigm that these complexes are restricted to certain genomic compartments (**Fig. 8**). In leukemia cells harboring MLL fusions, Menin-MLL and NF-Y complexes coordinately co-repress the expression of a tumor suppressive network that involves the *Cdkn2cllnk4c* and *Cdkn2dllnk4d* tumor suppressor genes in leukemia by direct binding to gene promoters. Disruption of Menin-MLL or NF-Y complexes using genetic or pharmacological approaches triggers a molecular switch that leads to recruitment of the MLL3/4-UTX complex to the promoters of tumor suppressive genes, among others, leading to a UTX-dependent increase in the levels of gene activation-associated histone modifications and a concomitant increase in gene expression(72,78,80). Importantly, these phenotypes are independent of UTX catalytic activity as the first 500 amino acids of UTX (lacking the histone demethylase domain) are sufficient to induce full length UTX-dependent cellular and transcriptional phenotypes. This finding is particularly interesting given a recent study suggesting that UTX requires a much bigger protein fragment to drive tumor suppressive activity and transcriptional regulation via phase separation mechanisms(78). Instead, our study demonstrates that UTX does not require a core intrinsically disordered region to drive tumor suppressive responses in the context of Menin-MLL inhibition. These results establish a new paradigm by which UTX employs non-catalytic mechanisms to regulate gene expression and cellular phenotypes that impact response to epigenetic therapies.

**Figure 8.**
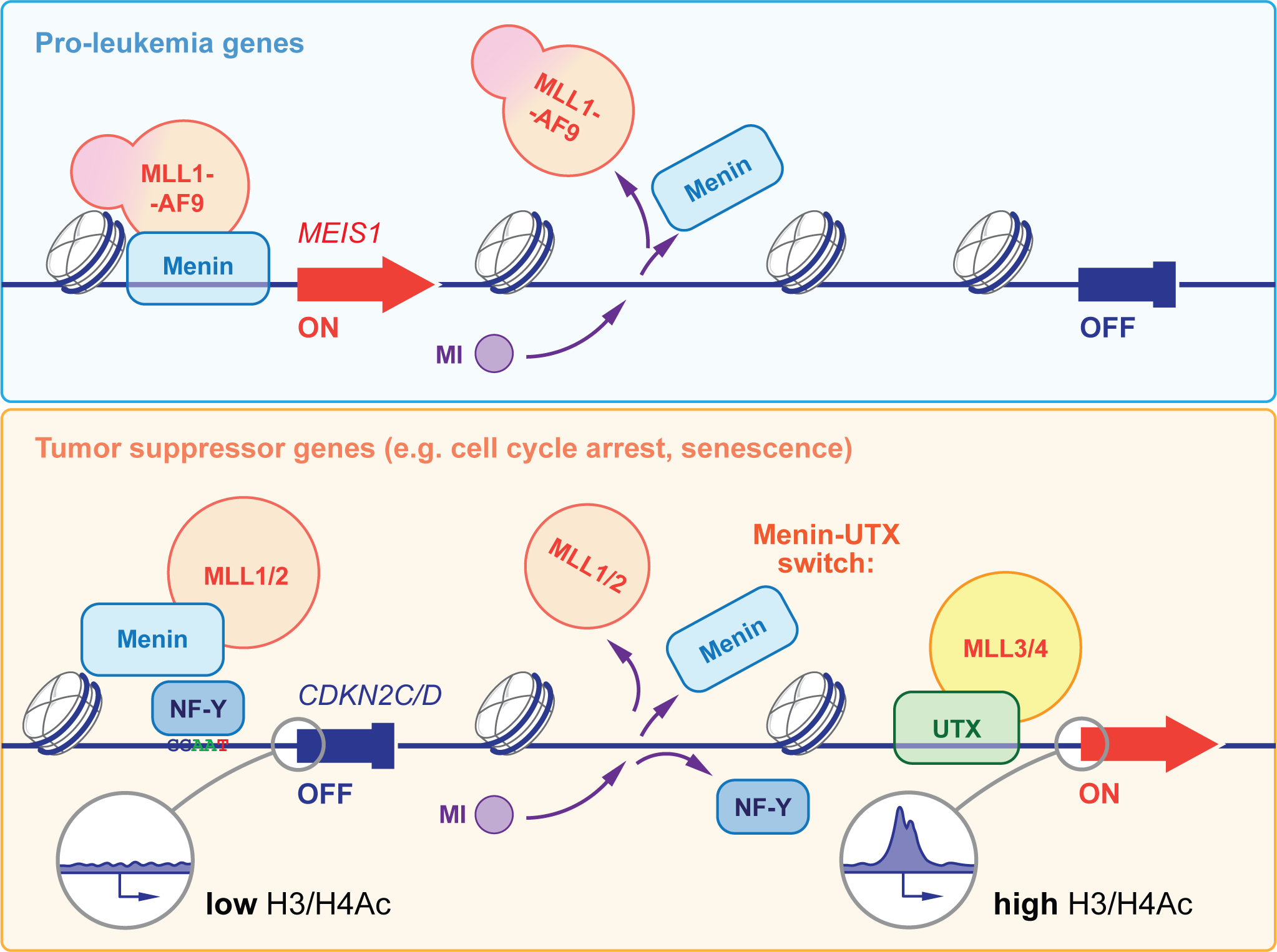
Molecular switch between the mammalian MLL complexes dictates cellular response to Menin-MLL inhibition

Our findings have important therapeutic implications by establishing a molecular mechanism that regulates response and resistance of leukemia cells to Menin-MLL inhibitors, which could guide clinical treatment. We show that combinatorial targeting of this molecular axis using Menin-MLL and CDK4/6 inhibitors has a strong synergistic effect on inhibiting leukemia cell proliferation *in vitro* and *in vivo*, including in a non-responsive leukemia PDX model. Importantly, we show this mechanism is at play in human AML subjects in the context of the AUGMENT-101 clinical trial (NCT04065399).

As Palbociclib and other CDK4/6 inhibitors are already FDA-approved(93)(85), our findings suggest that this combination treatment could be a viable therapeutic option to both boost the effectiveness of Menin inhibitors as monotherapies while potentially overcoming the intrinsic resistance to Menin-MLL inhibition conferred by loss-of-function of the MLL3/4-UTX complex.

## MATERIALS AND METHODS

### Plasmids and sgRNA cloning

To generate stable Cas9-expressing cell lines, we used lentiCas9-Blast (Addgene, 52962). Human wild-type or mutant (K4M) histone H3.1 were cloned into pCDH-EF1-MCS-IRES-RFP (System Biosciences, CD531A-2). To express sgRNAs, we generated the pUSEPR (U6-sgRNA-EFS-Puro-P2A-TurboRFP) and pUSEPB (U6-sgRNA-EFS-Puro-P2A-TagBFP) lentiviral vectors by Gibson assembly of the following DNA fragments: (i) PCR-amplified U6-sgRNA (improved scaffold)(94) cassette, (ii) PCR-amplified EF1s promoter, (iii) PCR-amplified Puro-P2A-TurboRFP (or -TagBFP) gene fragment (IDT), and (iv) BsrGI/PmeI-digested pLL3-based lentiviral backbone(95). For sgRNA cloning, pUSEPR and pUSEPB vectors were linearized with BsmBI (NEB) and ligated with BsmBI-compatible annealed and phosphorylated oligos encoding sgRNAs using high concentration T4 DNA ligase (NEB). All sgRNA sequences used are listed in Supplementary Table 1.

### Cell culture

Mouse MLL-AF9 leukemia cells were kindly shared by David Chen (Chun-Wei Chen) and were originally generated by transformation of female mouse bone marrow Lin-Sca1+cKit+ (LSK) cells with an MSCV-IRES-GFP (pMIG) retrovirus expressing the human MLL-AF9 fusion protein and transplanted into sub-lethally irradiated recipient mice as described previously(29, 47). Leukemic blasts were harvested from moribund mice and cultured in vitro in IMDM (Gibco) supplemented with 15% FBS (Gibco), mouse IL-6 (10ng/µL, PeproTech), mouse IL-3 (10ng/µL, PeproTech), mouse SCF (20ng/µL, PeproTech), penicillin (100U/mL, Gibco), streptomycin (100µg/uL, Gibco), L-glutamine (2mM, Gibco), and plasmocin (5µg/mL, InvivoGen). Human leukemia cell lines MV4 11 and OCI-AML3 were kindly shared by Zhaohui Feng and were cultured in RPMI 1640 (Corning) supplemented with 10% FBS (Gibco), penicillin (100U/mL, Gibco), streptomycin (100µg/uL, Gibco), L-glutamine (2mM, Gibco), and plasmocin (5µg/mL, InvivoGen). Mouse NIH-3T3 cells were maintained in DMEM (Corning) supplemented with 10% FCS (ATCC), penicillin (100U/mL, Gibco), streptomycin (100µg/uL, Gibco), and plasmocin (5µg/mL, InvivoGen). Human HEK293 cells were maintained in DMEM (Corning) supplemented with 10% FBS (Gibco), penicillin (100U/mL, Gibco), streptomycin (100µg/uL, Gibco), and plasmocin (5µg/mL, InvivoGen). Cas9-expressing cells were generated by lentiviral transduction of lentiCas9-Blast followed by Blasticidin (InvivoGen) selection and validation of Cas9 expression and activity. All cells were confirmed to be free of Mycoplasma contamination and cultured at 37°C and 5% CO_2_.

### Virus production

Lentiviruses were produced by co-transfection of HEK293T (ATCC) cells with pUSEPR-EpiV2.0 sgRNA library, individual sgRNA plasmids, or lentiCas9-Blast, and packaging vectors psPax2 (Addgene, 12260) and pMD2.G (Addgene, 12259) using Lipofectamine 2000 (Invitrogen). Viral supernatants were collected at 48 and 72 hours post transfection and stored at -80°C.

### Transduction of cell lines

Leukemia cells were seeded at a density of 2.5 x 10^5^ cells/well of a non TC-treated 12-well plate in complete medium containing polybrene (10µg/mL, EMD Millipore), and then transduced with lentivirus by centrifugation at 2,500rpm for 90 minutes at 37°C. After a 24-hour incubation, cells were transferred to a new plate containing fresh culture medium. Antibiotic selection or cell sorting was done 48 hours post transduction.

### Drug treatments

For MI-503 (Active Biochem) treatments, leukemia cells were seeded at a density of 4 x 10^5^ cells/mL, treated with limiting dilutions of the inhibitor as indicated or 0.25% DMSO (vehicle). Cells were re-plated every 4 days to the initial density and re-treated. Viability was assessed at various time points by using the CellTiter-Glo Luminescent Cell Viability assay (Promega) following the manufacturer s guidelines. Ratio of luminescence signal from metabolically active cells in MI-503 versus DMSO were plotted to calculate IC50 values (Prism 8, GraphPad). For MI-503 (Active Biochem) and Palbociclib HCl (Selleckchem) combination treatments, 25,000 leukemia cells in 250µL of drug-containing medium were seeded in a 48-well plate and viability was assessed every 4 days by using the CellTiter-Glo Luminescent Cell Viability assay (Promega). For RNA-Seq and ChIP-Seq experiments, cells were cultured at 4 x 10^5^ cells/mL, treated with MI-503 (concentrations as indicated in Fig.legends) or 0.25% DMSO for 4 days. Cells were collected, washed with PBS, pelleted, and flash-frozen before RNA or chromatin isolation. For *in vivo* VTP-50649 treatment, mice were randomly assigned to either normal or 0.1% VTP-50469 rodent special diet^26^. Mice were bled weekly to monitor leukemia burden and euthanized when showing clinical signs of disease (experimental endpoint).

### Flow cytometric analyses

Immunophenotyping of leukemia cells treated with MI-503 (or vehicle), was done by collecting cells post treatment and staining using the indicated conjugated primary antibodies. Stained samples were analyzed on an LSRFortessa (BD Biosciences) flow cytometer. Data analysis was performed using FlowJo (BD Biosciences) software. Intracellular antigens detection was done by using the Foxp3/Transcription Factor Staining Buffer Set (eBioscience) following the manufacturer s guidelines. Conjugated primary antibodies used were: Pacific Blue anti-CD11b (Biolegend, 101224), Alexa Fluor 647 anti-Cas9 (CST, 48796).

### Xenograft models of AML

All animal experiments were performed with the approval of Dana-Farber Cancer Institute s Institutional Animal Care and Use Committee (IACUC). NOD.Cg-*Prkdc^scid^ l/2rg^tm1Sug^*/JicTac (NOG) mice were obtained from Taconic Biosciences (Rensselaer, NY, USA). Non-irradiated 8-12 weeks old adult mice were transplanted with previously established patient-derived xenografts (PDXs)(25)via tail vein injection (250,000 cells/mouse). Engraftment of human cells (hCD45^+^) was analyzed and monitored longitudinally by weekly bleeding to quantify hCD45^+^ cells in the peripheral blood by flow cytometry with human CD45-PE and anti-mouse CD45-APC-Cy7 antibodies (Biolegend, San Diego, CA, USA). Mice were monitored closely to detect disease onset and treatment started when hCD45^+^ cells were detectable in the peripheral blood. Mice were randomly assigned to either normal or 0.1% VTP-50469 rodent special diet(25). Mice were bled weekly to monitor leukemia burden as described above and euthanized when showing clinical signs of disease (experimental endpoint). Leukemia cells from a subset of these animals were harvested after seven days of treatment to perform RNA-Seq.

### Longitudinal analysis of AML patient treated with SNDX-5613

Peripheral blood of patients was taken under informed consent according to the Declaration of Helsinki during routine blood draws at screening and different timepoints during the first cycle of treatment with SDNX-5613 within the AUGMENT-101 clinical trial (NCT04065399). Peripheral blood mononuclear cells (PBMCs) were subsequently isolated using Ficoll (BD Bioscience, Franklin Lakes, NJ, USA) gradient centrifugation, viably frozen, and banked at the Dana-Farber Cancer Institute, Boston, MA (approved institutional protocol, IRB: #01-206). For longitudinal analysis, samples were thawed, washed twice in PBS, and stained with anti-human CD45 (PE) and anti-human CD117 (APC) (Biolegend, San Diego, CA, USA). CD45-low/CD117+ leukemia cells were FACS sorted (MA900 sorter, Sony Biotechnology, San Jose, CA, USA) and subsequently processed for RNA-Seq (see methods section on RNA-Seq).

### Immunoblotting

Whole cell lysates were separated by SDS-PAGE, transferred to a PVDF membrane (EMD Millipore), blocked in 5% non-fat milk in TBS plus 0.5% Tween-20 (Sigma-Aldrich), probed with primary antibodies, and detected with HRP-conjugated anti-rabbit or anti-mouse secondary antibodies (GE Healthcare). Primary antibodies used included: anti-Cas9 (CST, 14697), anti-UTX (CST, 33510), anti-Menin (Bethyl, A300-105A), anti-NF-YA (Santa Cruz Biotechnology, sc-17753), anti-Actin (abcam, ab8224), anti-HSP90 (BD Biosciences, 610418), anti-HA (Biolegend, 901501), anti-H3K4me1 (abcam, ab8895), anti-H3K4me3 (CST, 9751), anti-H3 (abcam, ab1791).

### Locus-specific DNA sequencing

To determine the mutational status of the Men1 and Utx loci in cells targeted by CRISPR-Cas9, we performed next generation sequencing of PCR-amplified target regions. Genomic DNA (gDNA) was isolated using the DNeasy Blood & Tissue Kit (Qiagen) following the manufacturer’s guidelines. Amplification of target regions was performed from 500ng of gDNA using Q5 High-Fidelity 2X Master Mix (NEB) and primers listed in Supplementary Table 1. PCR products were purified using the QIAquick PCR Purification Kit (Qiagen) and sequenced on Illumina instruments at GENEWIZ (Amplicon-EZ service).

### RNA isolation, qRT-PCR analyses, and RNA sequencing

Total RNA was isolated from cells using the RNeasy kit (Qiagen, Hilden, Germany). RNA was reverse transcribed with High-Capacity cDNA Reverse Transcription kits (Applied Biosystems) following the manufacturer s instructions. Quantitative PCRs (qRT-PCRs) were performed using the TaqMan Gene Expression Master Mix (Applied Biosystems) with the StepOne Real-Time System (Applied Biosystems). TaqMan gene expression assays were used. ActB was used as the endogenous control for normalization and relative gene expression was calculated by using the comparative CT method. The mouse gene probes used were: *ActB* (Mm02619580_g1), *Hoxa9* (Mm00439364_m1), *Meis1* (Mm00487664_m1), *Utx* (*Kdm6a*) (Mm00801998_m1), *Men1* (Mm00484957_m1). Quality of extracted RNA for sequencing was assessed by RIN using a Bioanalyzer (Agilent) and quantified by TapeStation (Agilent). Poly(A) mRNA enrichment and library preparation was performed using the NEBNext Poly(A) mRNA Magnetic Isolation Module and NEBNext Ultra II RNA Library Prep kit (NEB). Sequencing was done using the Illumina NextSeq500 to obtain >20 million 75bp single-end or 37bp paired-end reads per sample or at Genewiz (HiSeq, 150bp, paired end) (Illumina, South Plainfield, NJ, USA).

### Cloning of sgRNA library targeting mouse chromatin regulators

sgRNA sequences (six per gene) targeting 616 mouse chromatin regulators (for a total of 3,696 sgRNAs) (**Supplementary Table 1**) were designed using the Broad Institute sgRNA Designer too(96). We also included 36 non-targeting control sgRNAs obtained from the GeCKOv2 Mouse CRISPR library(97)(for a total of 3,732 sgRNAs). This library was divided into 6 pools (each composed of 616 targeting and 6 non-targeting sgRNAs), synthesized by Agilent Technologies, and cloned into the pUSEPR lentiviral vector(98) using a modified version of the protocol published by Doench *et a/.*(*96*) to ensure a library representation of >10,000X. Briefly, each sub-pool was selectively amplified using barcoded forward and reverse primers that append cloning adapters at the 5 - and 3 -ends of the sgRNA insert (**Supplementary Table 1**), purified using the QIAquick PCR Purification Kit (Qiagen), and ligated into BsmBI-digested and dephosphorylated pUSEPR vector using high-concentration T4 DNA ligase (NEB). A minimum of 1.2ug of ligated pUSEPR plasmid DNA per sub-pool was electroporated into Endura electrocompetent cells (Lucigen), recovered for one hour at 37°C, plated across four 15cm LB-Carbenicillin plates (Teknova), and incubated at 37°C for 16 hours. The total number of bacterial colonies per sub-pool was quantified using serial dilution plates to ensure a library representation of >10,000X (>6.2 million colonies per sub-pool). The next morning, bacterial colonies were scraped and briefly expanded for 4 hours at 37°C in 500mL of LB-Carbenicillin. Plasmid DNA was isolated using the Plasmid Plus Maxi Kit (Qiagen).

To assess sgRNA distribution in each of the sub-pools, as well as the master pool (composed of equimolar amounts of plasmid DNA from each individual sub-pool), we amplified the sgRNA target region using primers that append Illumina sequencing adapters on the 5 - and 3 -ends of the amplicon, as well as a random nucleotide stagger and unique demultiplexing barcode on the 5 -end (**Supplementary Table 1**). Library amplicons were size-selected on a 2.5% agarose gel, purified using the QIAquick Gel Extraction Kit (Qiagen), and sequenced on an Illumina NextSeq instrument (75nt single end reads).

### Chromatin-focused CRISPR-Cas9 genetic screening

To ensure that most cells harbor a single sgRNA integration event, we determined the volume of viral supernatant that would achieve an MOI of ∼0.3 upon spinfection of a population of Cas9-expressing leukemia cells. Briefly, cells were plated at a concentration of 2.5x10^5^ per well in 12-well plates along with increasing volumes of master pool viral supernatant (0, 25, 100, 200, 500, 1000, and 2000µL) and polybrene (10µg/mL, EMD Millipore). Cells were then centrifuged at 1,500rpm for 2 hours at 37°C and incubated at 37°C overnight. Viral infection efficiency was determined by the percentage of tRFP+ cells assessed by flow cytometry on an LSRFortessa (BD Biosciences) instrument 72 hours post infection.

Each step of the screen - from infection to sequencing - was optimized to achieve a minimum representation of 1000X. To ensure a representation of 1000X at the transduction step, we spinfected a total of 20 million cells across seven 12-well plates in triplicate (for a total of twenty one 12-well plates) using the volume of viral supernatant that would achieve a 30% infection rate (6 million transduced cells per technical replicate).

24 hours after infection, cells were pooled into 2 x T225 flasks (Corning) per infection replicate and selected with 2.5µg/mL puromycin (Gibco) for 4 days. Subsequently, 6 million puromycin-selected cells were pelleted and stored at -20°C (T_0_/Input population) while the rest were plated into either DMSO- or MI-503-containing media (at an IC50 concentration) and cultured until the population reached 12 cumulative population doublings (T_F_/Final). At least 6 million cells were harvested and pelleted for this final time point. Genomic DNA from MLL-AF9 cells was isolated using the DNeasy Blood & Tissue Kit (Qiagen) following the manufacturer’s guidelines.

As previously published(99), we assumed that each cell contains approximately 6.6pg of genomic DNA (gDNA). Therefore, deconvolution of the screen at 1000X required sampling ∼4 million x 6.6pg of gDNA, or ∼26.4µg. We employed a modified 2-step PCR version of the protocol published by Doench *et a/.*(*96*). Briefly, we perform an initial “enrichment” PCR, whereby the integrated sgRNA cassettes are amplified from gDNA, followed by a second PCR to append Illumina sequencing adapters on the 5 - and 3 -ends of the amplicon, as well as a random nucleotide stagger and unique demultiplexing barcode on the 5 end. Each “PCR1” reaction contains 25µL of Q5 High-Fidelity 2X Master Mix (NEB), 2.5µL of Nuc PCR#1 Fwd Primer (10µM), 2.5µL of Nuc PCR#1 Rev Primer (10µM), and 5µg of gDNA in 20µL of water (for a total volume of 50µL per reaction). The number of PCR1 reactions is scaled accordingly therefore, we performed six PCR1 reactions per technical replicate, per time point (T0 or TF), and per condition (DMSO or MI-503). PCR1 amplicons were purified using the QIAquick PCR Purification Kit (Qiagen) and used as template for “PCR2” reactions. Each PCR2 reaction contains 25µL of Q5 High-Fidelity 2X Master Mix (NEB), 2.5µL of a unique Nuc PCR#2 Fwd Primer (10µM), 2.5µL of Nuc PCR#2 Rev Primer (10µM), and 300ng of PCR1 product in 20µL of water (for a total volume of 50µL per reaction). We performed two PCR2 reactions per PCR1 product. Library amplicons were size-selected on a 2.5% agarose gel, purified using the QIAquick Gel Extraction Kit (Qiagen), and sequenced on an Illumina NextSeq500 instrument (75nt single end reads). All primer sequences are available in Supplementary Table 1. PCR Program for PCR1 and PCR2: 1) 98°C x 30s 2) 98°C x 10s 3) 65°C x 30s 4) 72°C x 30s 5) Go to step 2 x 24 cycles 6) 72°C x 2 min 7) 4°C forever.

### Genome-wide CRISPR-Cas9 genetic screening

Paired mouse genome-scale CRISPR-Cas9 screening libraries (M1/M2) were provided by Shengqing Gu and Xiaole Shirley Liu (Addgene Pooled Library #1000000173). The M1 and M2 libraries cover protein coding genes of the genome with a total of 10 guide RNAs per gene. Lentivirus was produced using each separate library pool and used to transduce each 5x10^8^ MLL-AF9 cells at low MOI. 48 hours after library transduction, cells were selected with blasticidin (5µg/ml). After 5 days of antibiotic selection, a baseline (T0) sample was collected, and cells were cultured in duplicate before harvest of terminal samples after 12 days (TF). Subsequently, genomic DNA was isolated using phenol-chloroform extraction and sgRNA libraries were deconvoluted using next-generation sequencing essentially as described above.

### Analysis of CRISPR-Cas9 genetic screen data

FASTQ files were processed and trimmed to retrieve sgRNA target sequences followed by alignment to the reference sgRNA library file. Sequencing read counts were summarized at gene level per sample and used as input to run differential analysis using DESeq2 package. The log2 fold change values were used as ’Gene Score for the final visualization. Genome-wide screening data was analyzed using MAGeCK MLE essentially as described in the original publication(100). See **Supplementary Table 2** for all raw screening data.

### Growth competition assays

Cas9-expressing cells were virally transduced with the designated constructs (pUSEPR-sgRNA, pUSEPB-sgRNA, pCDH-cDNA) in 12-well plates at 30-40% infection rate (three infection replicates). Cells were monitored by flow cytometry over time using an LSRFortessa (BD Biosciences) flow cytometer and relative growth of sgRNA-containing cells was assesed. Flow cytometry data was analyzed with FlowJo software (BD Biosciences). The percentage of single positive (SP) (tRFP+ or BFP+) or double positive (DP) (tRFP+/BFP+) cells was normalized to their respective “T0” time-point values (assessed on day 2 or 3 post transduction, as indicated in the Fig.legend). Normalized values were log2-transformed, and the differential fitness of cells was calculated as follow: Differential Fitness = log2(Normalized DP) - log2(Normalized SP)

### Chromatin immunoprecipitation (ChIP)

Cross-linking ChIP in mouse leukemia and NIH-3T3 cells was performed with 10-20x10^7^ cells per immunoprecipitation. After drug (or vehicle) treatment, cells were collected, washed once with ice-cold PBS, and flash-frozen. Cells were resuspended in ice-cold PBS and cross-linked using 1% paraformaldehyde (PFA) (Electron Microscopy Sciences) for 5 minutes at room temperature with gentle rotation. Unreacted PFA was quenched with glycine (final concentration 125mM) for 5 minutes at room temperature with gentle rotation. Cells were washed once with ice-cold PBS and pelleted by centrifugation (800g for 5 minutes). To obtain a soluble chromatin extract, cells were resuspended in 1mL of LB1 (50mM HEPES pH 7.5, 140mM NaCl, 1mM EDTA, 10% glycerol, 0.5% NP-40, 0.25% Triton X-100, 1X complete protease inhibitor cocktail) and incubated at 4°C for 10 minutes while rotating. Samples were centrifuged (1400g for 5 minutes), resuspended in 1mL of LB2 (10mM Tris-HCl pH 8.0, 200mM NaCl, 1mM EDTA, 0.5mM EGTA, 1X complete protease inhibitor cocktail), and incubated at 4°C for 10 minutes while rotating. Finally, samples were centrifuged (1400g for 5 minutes) and resuspended in 1mL of LB3 (10mM Tris-HCl pH 8.0, 100mM NaCl, 1mM EDTA, 0.5mM EGTA, 0.1% sodium deoxycholate, 0.5% N-Lauroylsarcosine, 1X complete protease inhibitor cocktail). Samples were homogenized by passing 7-8 times through a 28-gauge needle and Triton X-100 was added to a final concentration of 1%. Chromatin extracts were sonicated for 14 minutes using a Covaris E220 focused ultrasonicator. Lysates were centrifuged at maximum speed for 10 minutes at 4°C and 5% of supernatant was saved as input DNA. Beads were prepared by incubating them in 0.5% BSA in PBS and antibodies overnight (100µL of Dynabeads Protein A or Protein G (Invitrogen) plus 20µL of antibody). Antibodies used were: anti-Menin (Bethyl, A300-105A), anti-UTX (Bethyl, A302-374A), anti-MLL1 (N-term-specific, Bethyl, A300-086A), anti-MLL3/4 (kindly provided by the Wysocka laboratory^41^, anti-NF-YA (Santa Cruz Biotechnology, sc-17753), anti-H3K4me1 (abcam, ab8895), anti-H3K4me3 (Active Motif, 39159), and anti-H4K16ac (Active Motif, 39167). Antibody-Beads mixes were washed with 0.5% BSA in PBS and then added to the lysates overnight while rotating at 4°C. Beads were then washed six times with RIPA buffer (50mM HEPES pH 7.5, 500mM LiCl, 1mM EDTA, 0.7% sodium-deoxycholate, 1% NP-40) and once with TE-NaCl Buffer (10mM Tris-HCl pH 8.0, 50mM NaCl, 1mM EDTA). Chromatin was eluted from beads in Elution buffer (50mM Tris-HCl pH 8.0, 10mM EDTA, 1% SDS) by incubating at 65°C for 30 minutes while shaking, supernatant was removed by centrifugation, and crosslinking was reversed by further incubating chromatin overnight at 65°C. The eluted chromatin was then treated with RNaseA (10mg/mL) for 1 hour at 37°C and with Proteinase K (Roche) for 2 hours at 55°C. DNA was purified by using phenol-chloroform extraction followed with ethanol precipitation. The NEBNext Ultra II DNA Library Prep kit was used to prepare samples for sequencing on an Illumina NextSeq500 (75bp read length, single-end, or 37bp read length, paired-end).

### ChIP-Seq analysis

ChIP-sequencing samples were sequenced using the Illumina NextSeq500. ChIP-seq reads were aligned using Rsubread s align method and predicted fragment lengths calculated by the ChIPQC R Bioconductor package(101, 102). Normalized, fragment extended signal bigWigs were created using the rtracklayer R Bioconductor package. Peak calls were made in MACS2 software(103). Read counts in peaks were calculated using the featureCounts method in the Rsubread library(102). Differential ChIP-seq signal were identified using the binomTest from the edgeR R Bioconductor package(104). Annotation of genomic regions to genes, biological functions, and pathways were performed using the ChIPseeker R Bioconductor package(105). Meta-peak plots were produced using the soGGi package and ChIP-seq signal heatmaps generated using the Deeptools and profileplyr software(106). Plots showing ChIP-Seq read signal over transcription start sites (TSSs) were made with the ngs.plot software package (v2.61) (ref. (107)). Overlaps between peak sets were determined using the ChIPpeakAnno R Bioconductor package with a maximum gap between peaks set to 1kb (ref. (108)). Peaks were annotated with both genes and the various types of genomic regions using the ChIPseeker R Bioconductor package(105). Range-based heatmaps showing signal over genomic regions were generated using the soGGi and profileplyr R Bioconductor package to quantify read signal and group the peak ranges and the deepTools software package (v3.3.1) to generate the heatmaps(106). Any regions included in the ENCODE blacklisted regions of the genome were excluded from all region-specific analyses(109). For some ChIP-seq experiments, raw Illumina NextSeq BCL files were converted to FASTQs using Illumina bcl2fastq v02.14.01.07 and reads trimmed using Trimmomatic v0.36 (phred quality threshold 33) and uploaded to the to the Basepair-server (basepairtech.com). Alignment and ChIP-seq QC was performed on the basepair platform (Bowtie2). Peak calling was performed using MACS (v.1.4) within the basepair platform utilizing the default parameters.

### RNA-Seq analysis

RNA-Seq samples were sequenced using the Illumina NextSeq500. Transcript abundance was computed from FASTQ files using Salmon and the GENCODE reference transcript sequences, transcript counts were imported into R with the tximport R Bioconductor package, and differential gene expression was determined with the DESeq2 R Bioconductor package(110–112). The data was visualized using the ggplot2 R package. Normalized counts were extracted from the DESeq2 results and z-scores for the indicated gene sets were visualized using both heatmaps and boxplots. Heatmaps showing gene expression changes across samples were generated using the pheatmap R package and boxplots were made with the ggplot2 R package. Gene ontology analysis using the KEGG 2019 database was performed using the Enrichr tool(67).

### Statistical analyses

Statistical tests were used as indicated in Figure legends. Generation of plots and statistical analyses were performed using Prism 8 (GraphPad). Error bars represent standard deviation, unless otherwise noted. We used Student s t-test (unpaired, two-tailed) to assess significance between treatment and control groups, and to calculate P values. P<0.05 was considered statistically significant.

### Source data availability

Data supporting the findings of this study are reported in Supplementary Figures 1-24 and Supplementary Tables 1-4. All raw data corresponding to high-throughput approaches (CRISPR screens, RNA-Seq, and ChIP-Seq) is available through NCBI GEO (Accession: GSE186711). All reagents and materials generated in this study will be available to the scientific community through Addgene and/or MTAs. Further information and requests for resources and reagents should be directed to and will be fulfilled by the Lead Contacts: C. David Allis, Scott W. Lowe, and Scott. A. Armstrong.

## ACKNOWLEDGEMENTS

We thank members of the Allis, Lowe, and Armstrong laboratories for their help and support Richard Phillips, Robert G. Roeder, Tom W. Muir, Benjamin A. Garcia, and Charles J. Sherr for scientific discussions David Chen (Chun-Wei Chen) for mouse MLL-AF9 cells, Zhaohui Feng for human leukemia cells, and Laura Whitman (Agilent) for OLS support. C.D.A. was supported by US National Institutes of Health (NIH) grants (P01CA196539 and 5R01CA204639-03), the Leukemia and Lymphoma Society (LLS-SCOR 7006-13), and the Rockefeller University and St. Jude Children s Research Hospital Collaborative on Chromatin Regulation in Pediatric Cancer. S.W.L was supported by a NIH/NCI grant (R01 CA190261) and an Agilent Thought Leader Award, as well as the MSKCC cancer center support grant (P30 CA008748). S.W.L. is the Geoffrey Beene Chair of Cancer Biology and a Howard Hughes Medical Institute Investigator. S.A.A was supported by NIH grants CA176745, CA206963, CA204639 and CA066996. Y.M.S.F was supported by the Damon Runyon-Sohn Pediatric Cancer Fellowship (DRSG-21-17) and NIGMS-MOSAIC K99/R00 Career Development Award (1K99GM140265-01). F.J.S.R. was partially supported by the MSKCC TROT program (5T32CA160001), a MSKCC GMTEC Postdoctoral Researcher Innovation Grant, and is a HHMI Hanna Gray Fellow. F.P. was supported by the German Research Foundation (DFG, PE 3217/1-1) and a Momentum Fellowship award by the Mark Foundation for Cancer Research. D.W.B. was supported by a Ruth L. Kirschstein National Research Service Award (5F32CA217068). E.R.K. was supported by an F31 NRSA predoctoral fellowship from the NIH/NCI (F31CA192835).

## AUTHOR CONTRIBUTIONS

Y.M.S.F., F.J.S.R., F.P, S.A.A, S.W.L., and C.D.A. conceived the study and wrote the manuscript with input from all the authors Y.M.S.F., F.J.S.R., F.P., Y.X., L.G., M.C.B., and D.C. conducted experiments D.W.B., T.C., and Y.J.H. analyzed RNA-Seq and ChIP-Seq data Y.J.H. and E.R.K. analyzed CRISPR-screen data. S.G. and X.S.L. provided genome-wide CRISPR library. A.V.K., M.M., and E.dS. provided patient-derived xenograft samples. R.M.S. provided primary AML samples from patients in the Syndax trial A.S. provided conceptual advice. S.A.A, S.W.L., and C.D.A. supervised the study and secured funding.

---

**Supplementary Figure 1.**
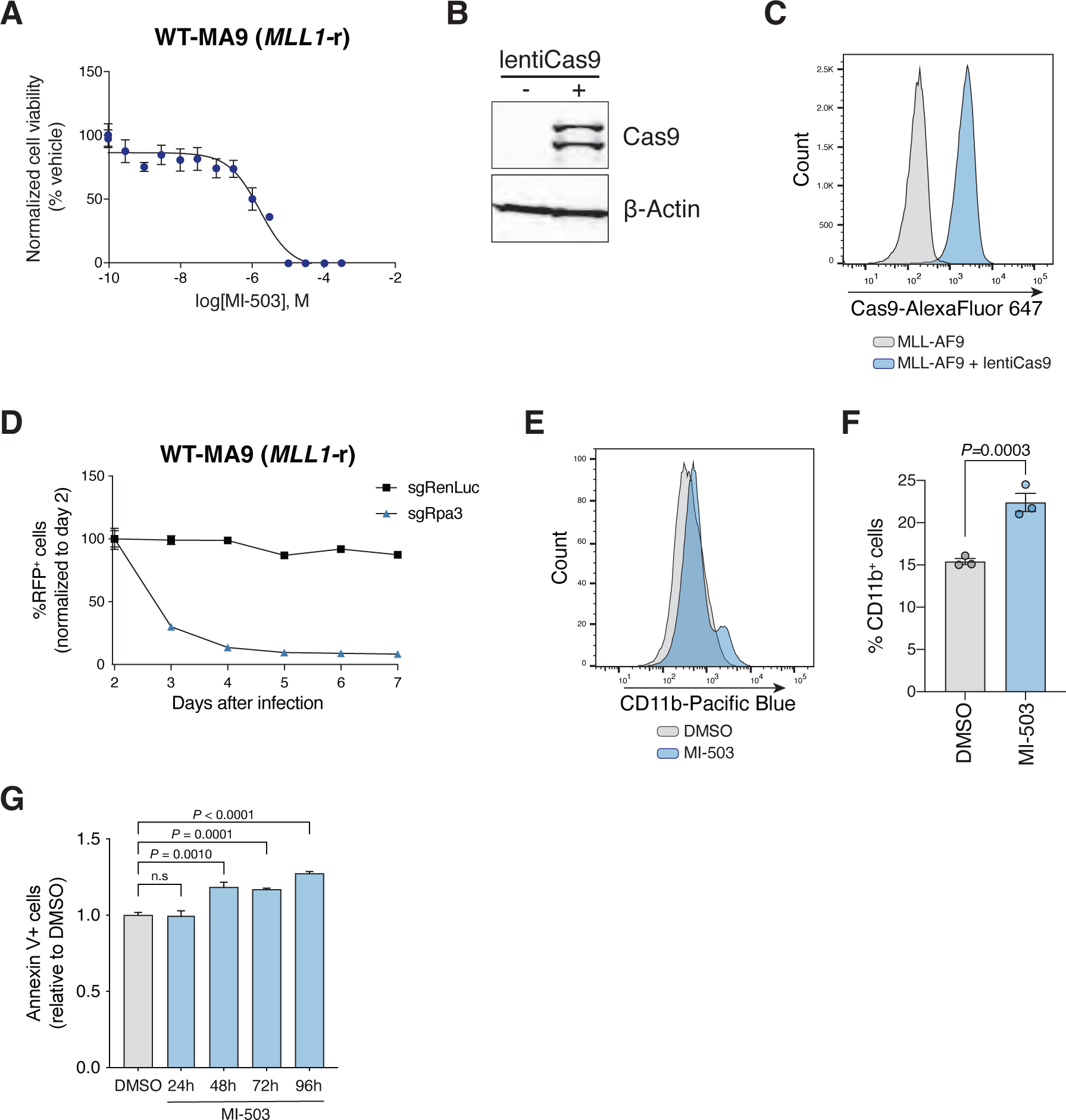
Generation and characterization of Cas9-expressing MLL-AF9 leukemia cells. **(A)** Dose response curve analysis of mouse MLL-AF9 leukemia (WT-MA9) cells treated with vehicle (DMSO) or Menin-MLL inhibitor (MI-503) for 96 hours (mean±SEM, n=3 replicates). **(B)** Immunoblot analysis of mouse Cas9-expressing MLL-AF9 leukemia cells. **(C)** Distribution of mouse Cas9-expressing MLL-AF9 leukemia cells determined by intracellular staining of Cas9 and flow cytometry. **(D)** Growth competition assay to test enzymatic activity of Cas9 in mouse MLL-AF9 leukemia cells (mean±SEM, n=3 infection replicates). **(E)** CD11b cell surface expression measured by flow cytometry for vehicle (DMSO, grey) or Menin-MLL inhibitor (MI-503, blue) treatment of mouse MLL-AF9 leukemia cells for 96 hours. **(F)** Quantification of %CD11b positive cells for vehicle (DMSO, grey) or Menin-MLL inhibitor (MI-503, blue) for 96 hours (mean±SEM, n=3 replicates, *P*-value calculated by Student’s t-test). **(G)** Quantification of %Annexin V positive cells after treatment with vehicle (DMSO, grey) or Menin-MLL inhibitor (MI-503, blue) at different time points (mean±SEM, n=3 replicates, *P*-value calculated by Student’s t-test).

**Supplementary Figure 2.**
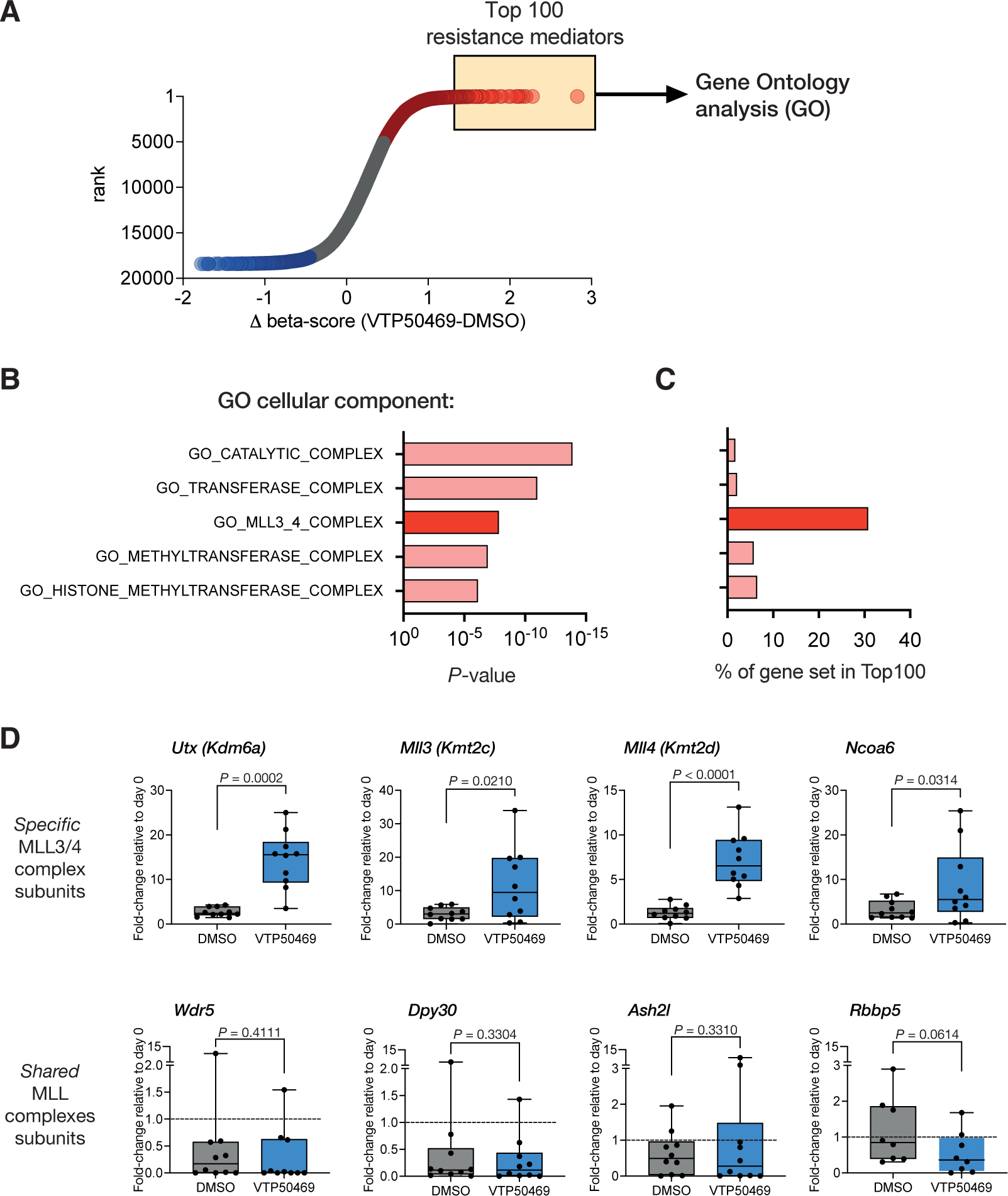
Genome-wide CRISPR screening identifies the MLL3/4-UTX complex as a key determinant of response to Menin-MLL inhibition. **(A)** Genome-wide screening data showing gene-level ranking based on differential enrichment of sgRNAs under Menin-MLL inhibitor treatment (VTP-50469) relative to vehicle (DMSO). Differential (Δ) beta-score between VTP-50469 and DMSO conditions was calculated using MaGeCK. A positive Δ beta-score denotes enrichment of specific gene-targeting sgRNAs. A negative Δ beta-score denotes depletion of specific gene-targeting sgRNAs. Red circles denote genes represented by enriched sgRNAs (genes whose inactivation promotes resistance to Menin-MLL inhibition). Blue circles denote genes represented by depleted sgRNAs (genes whose inactivation promotes sensitivity to Menin-MLL inhibition). The top 100 candidate mediators of resistance were selected for gene ontology (GO) analysis. **(B)** Top-scoring GO categories based on P-values obtained from GO analysis. **(C)** Percentage of genes within a given gene set that scored among the top 100 candidate mediators of resistance. **(D)** Fold-change of individual sgRNAs relative to T_0_ from either VTP-50469-treated or DMSO-treated cells (*P*-value calculated by Student’s t-test).

**Supplementary Figure 3.**
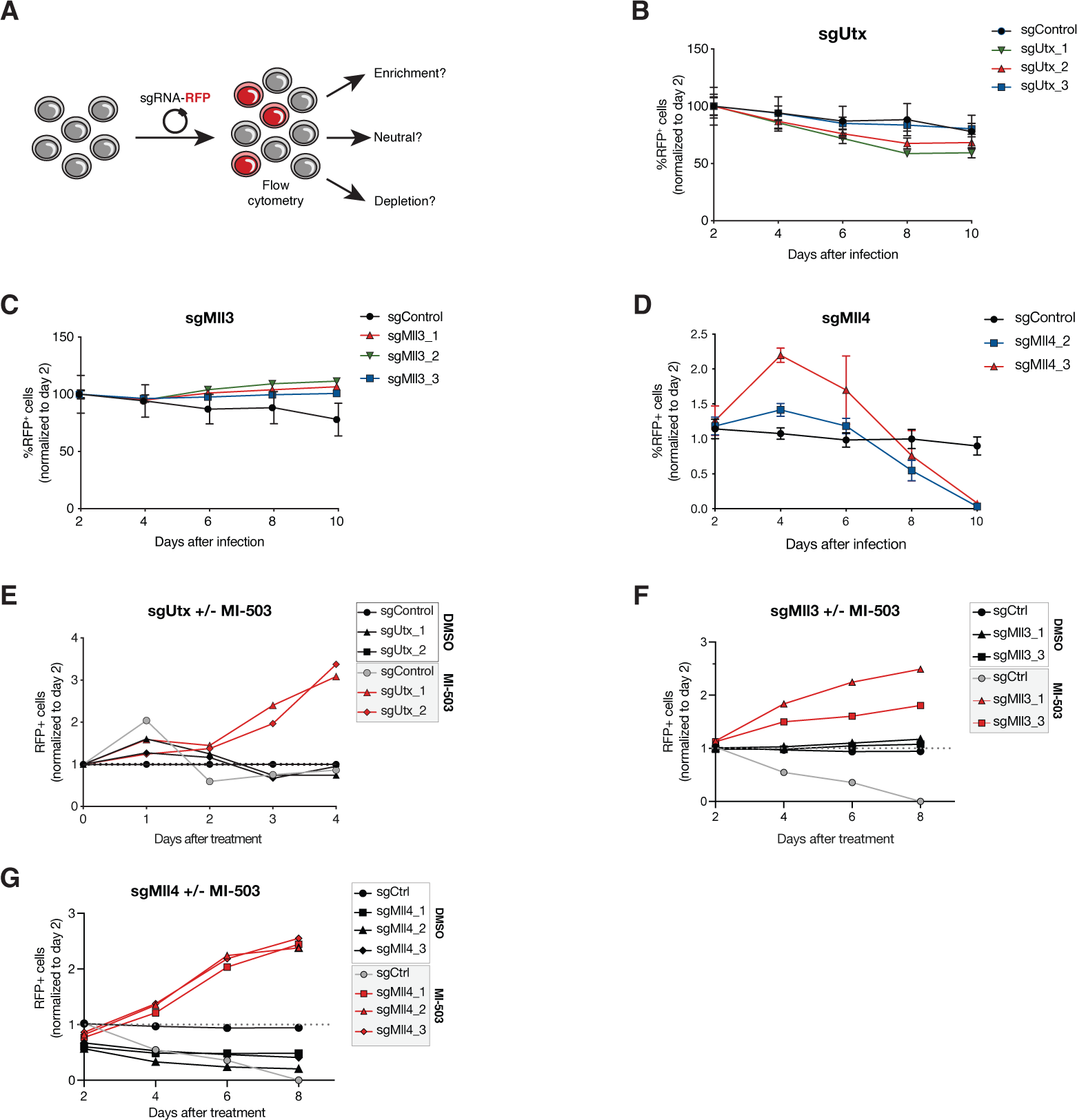
MLL3/4-UTX complex dictates therapeutic response of leukemia cells to Menin-MLL inhibition. **(A)** Layout of growth competition assay to assess the effect of genetic loss of *Utx*, *Mll3*, or *Mll4* in MLL-AF9 leukemia cells. **(B)** Growth competition assay in mouse MLL-AF9 cells. Graph shows the relative growth of cells infected with RFP-tagged *Utx* sgRNAs measured by flow cytometry (mean±SEM, n=3 infection replicates). sgControl targets a non-genic region in chromosome 8 as negative control. **(C)** Growth competition assay in mouse MLL-AF9 cells. Graph shows the relative growth of cells infected with RFP-tagged *Mll3* sgRNAs measured by flow cytometry (mean±SEM, n=3 infection replicates). sgControl targets a non-genic region in chromosome 8 as negative control. **(D)** Growth competition assay in mouse MLL-AF9 cells. Graph shows the relative growth of cells infected with RFP-tagged *Mll4* sgRNAs measured by flow cytometry (mean±SEM, n=3 infection replicates). sgControl targets a non-genic region in chromosome 8 as negative control. **(E)** Relative percentage of sgUtx-expressing (RFP+) cells over time after transduction of mouse MLL-AF9 cells treated with vehicle (DMSO) or Menin-MLL inhibitor (MI-503) (mean±SEM, n=3 infection replicates). **(F)** Relative percentage of sgMll3-expressing (RFP+) cells over time after transduction of MLL-AF9 cells treated with vehicle (DMSO) or Menin-MLL inhibitor (MI-503) (mean±SEM, n=3 infection replicates). **(G)** Relative percentage of sgMll4-expressing (RFP+) cells over time after transduction of MLL-AF9 cells treated with vehicle

**Supplementary Figure 4.**
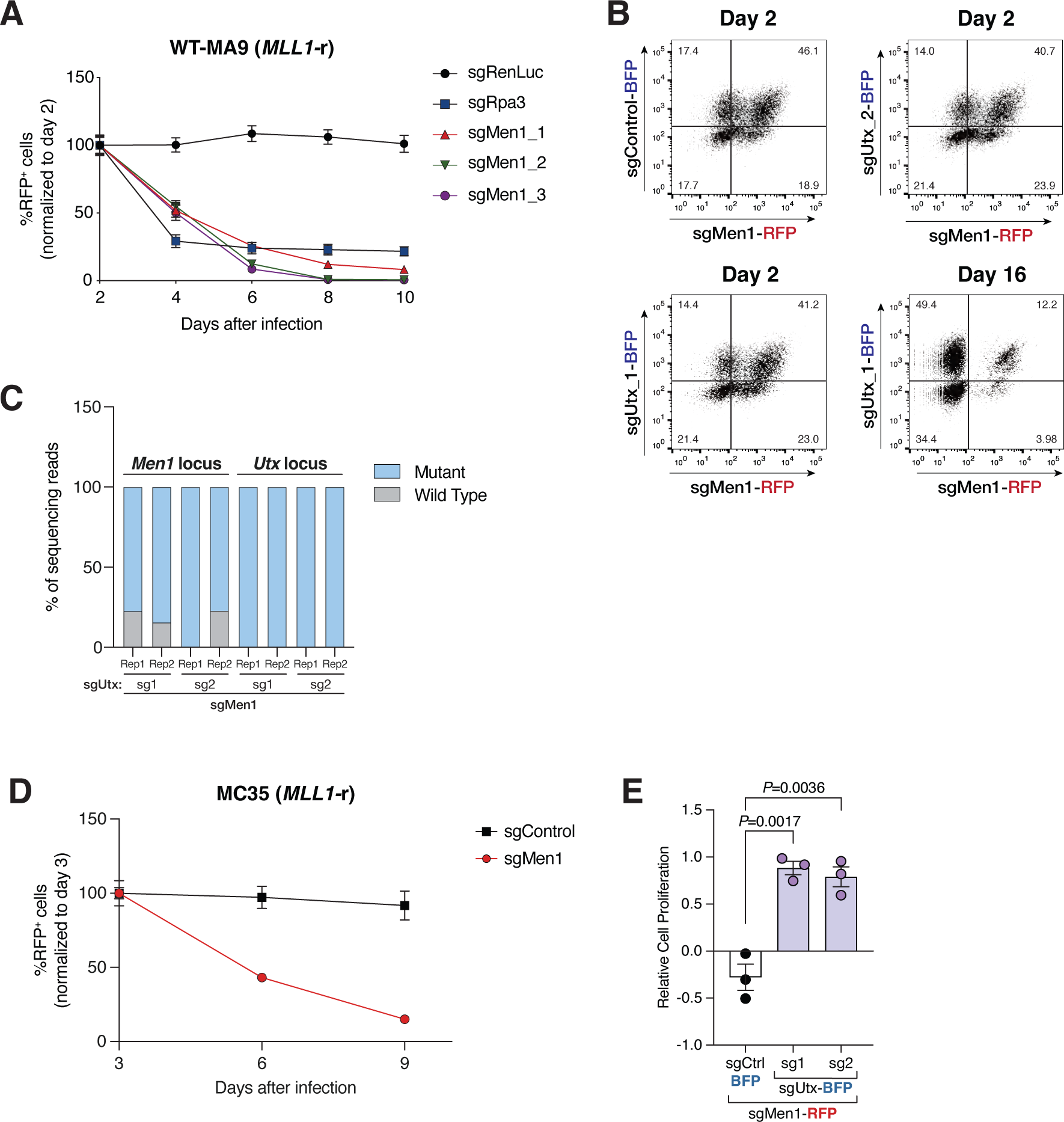
*Utx* deletion suppresses the lethality phenotype associated with *Men1* loss in leukemia. **(A)** Growth competition assay in mouse MLL-AF9 cells. Graph shows the relative growth of cells infected with RFP-tagged *Men1* sgRNAs measured by flow cytometry (mean±SEM, n=3 infection replicates). sgRenilla Luciferase (RenLuc) and sgRpa3 were negative and positive controls, respectively. **(B)** Plots from flow cytometry analysis of leukemia cells co-expressing sgMen1-RFP and sgUtx-BFP (sg1 or sg2) at 2 and 16 days post-infection. **(C)** Amplicon sequencing results of the mouse *Men1* and *Utx* loci from leukemia cells co-expressing sgMen1 and sgUtx (sg1 or sg2). **(D)** Growth competition assay in an independently-derived mouse MLL-AF9 cell line. Graph shows the relative growth of cells infected with RFP-tagged sgRNAs (sgControl and sgMen1) measured by flow cytometry (mean±SEM, n=3 infection replicates). **(E)** Differential fitness of mouse MLL-AF9 cells is shown as the relative fitness of double positive cells (sgMen1-RFP + sgUtx-BFP or sgMen1-RFP + sgControl-BFP) to single positive cells (sgMen1-RFP) 9 days post-infection measured by flow cytometry (mean±SEM, n=3 infection replicates, *P*-value calculated by Studentₑ t-test)

**Supplementary Figure 5.**
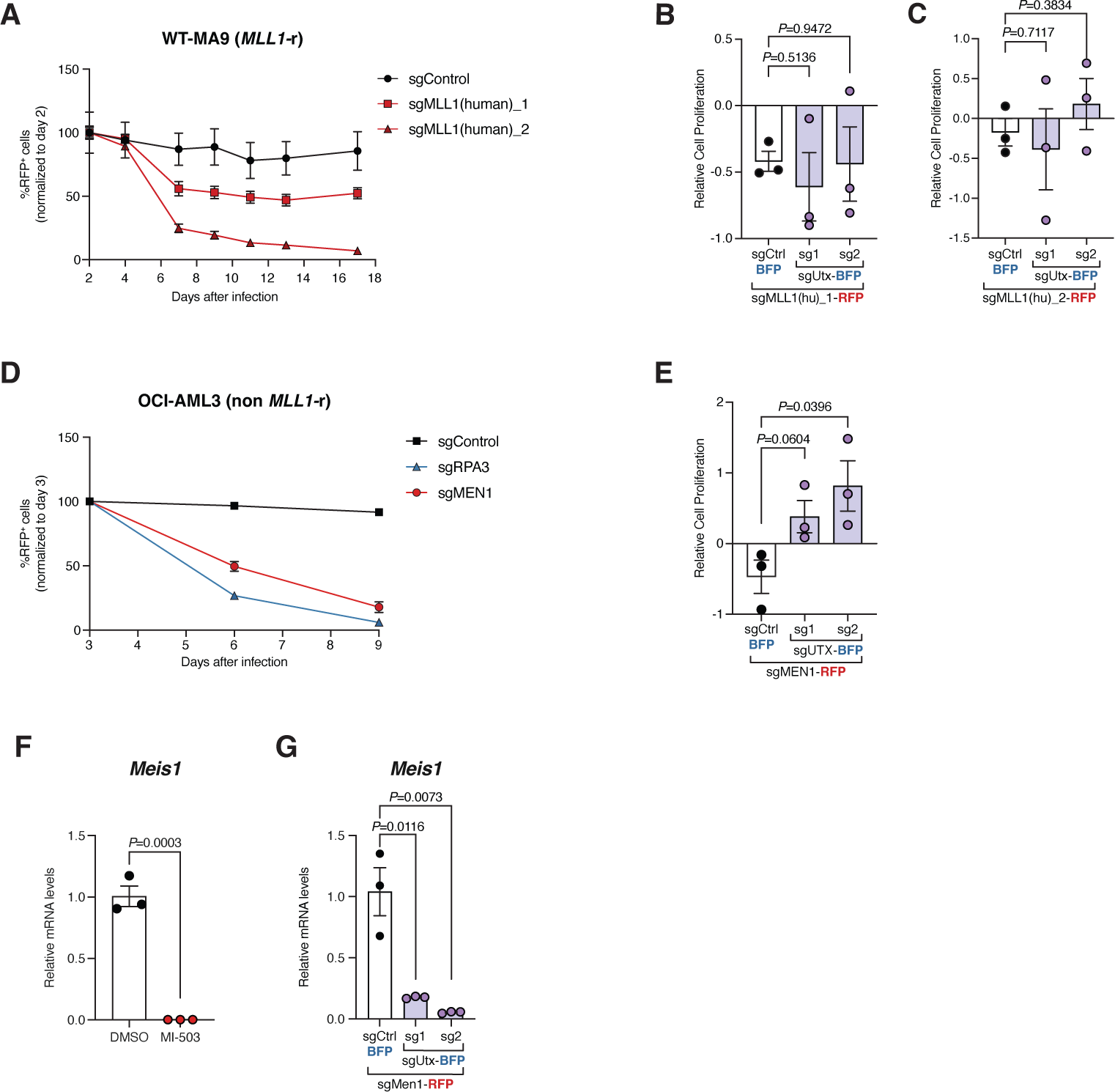
*Men1* and *Utx* epistasis is independent of the MLL-fusion. **(A)** Growth competition assay in mouse MLL-AF9 leukemia cells. Graph shows the relative growth of cells, infected with RFP-tagged sgRNAs that target the 5’-end of the human *MLL1* gene present in the MLL-AF9 fusion, measured by flow cytometry (mean±SEM, n=3 infection replicates). **(B)** Differential fitness of mouse MLL-AF9 leukemia cells is shown as the relative fitness of double positive cells (sgMLL1(hu)_1-RFP + sgUtx-BFP or sgMLL1(hu)_1-RFP + sgControl-BFP) to single positive cells (sgMLL1(hu)_1-RFP) 16 days post-infection measured by flow cytometry (mean±SEM, n=3 infection replicates, *P*-value calculated by Student’s t-test). **(C)** Differential fitness of mouse MLL-AF9 cells is shown as the relative fitness of double positive cells (sgMLL1(hu)_2-RFP + sgUtx-BFP or sgMLL1(hu)_2-RFP + sgControl-BFP) to single positive cells (sgMLL1(hu)_2-RFP) 16 days post-infection measured by flow cytometry (mean±SEM, n=3 infection replicates, *P*-value calculated by Student’s t-test). **(D)** Growth competition assay in human non *MLL1*-r (OCI-AML3) cells. Graph shows the relative growth of cells infected with RFP-tagged sgRNAs measured by flow cytometry (mean±SEM, n=3 infection replicates). **(E)** Differential fitness of human OCI-AML3 cells is shown as the relative fitness of double positive cells to single positive cells 20 days post-infection measured by flow cytometry (mean±SEM, n=3 infection replicates, *P*-value calculated by Student’s t-test). **(F)** qPCR analysis for *Meis1* expression in mouse MLL-AF9 leukemia cells treated with vehicle (DMSO, black) or Menin-MLL inhibitor (MI-503, red) for 96 hours (mean±SEM, n=3 replicates, *P*-value calculated by Student’s t-test). **(G)** qPCR analysis for *Meis1* expression in mouse MLL-AF9 leukemia cells co-expressing sgMen1-RFP and sgUtx-BFP, or a control sgRNA (mean±SEM, n=3 replicates, *P*-value calculated by Student’s t-test).

**Supplementary Figure 6.**
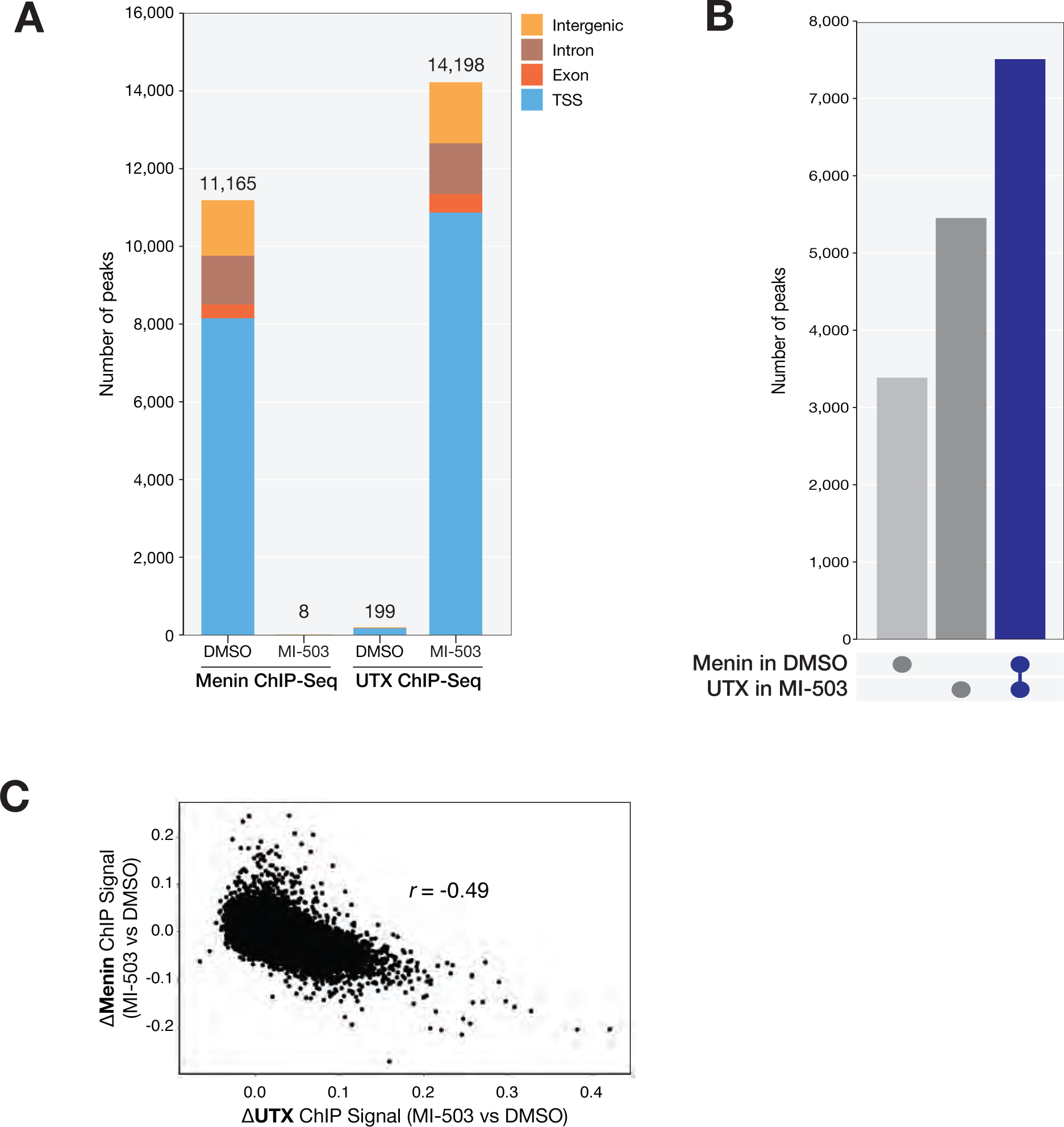
Distinct genomic distribution of Menin and UTX in the context of Menin-MLL inhibition. **(A)** Genomic distribution of Menin and UTX ChIP-Seq peaks from mouse MLL-AF9 leukemia cells treated with vehicle (DMSO) or Menin-MLL inhibitor (MI-503) for 96 hours. **(B)** Upset plot showing a comparison between the genomic distribution of Menin ChIP-Seq peaks in vehicle (DMSO, light grey), genomic distribution of UTX ChIP-Seq peaks in Menin-MLL inhibitor (MI-503, grey), and the overlap between UTX ChIP-Seq peaks in MI-503 and Menin ChIP-Seq peaks in DMSO (blue). **(C)** ChIP-Seq normalized reads per 10-kb bin for Menin (MI-503 vs DMSO, y-axis) and UTX (MI-503 vs DMSO, x-axis). Pearson’s correlation coefficient is indicated.

**Supplementary Figure 7.**
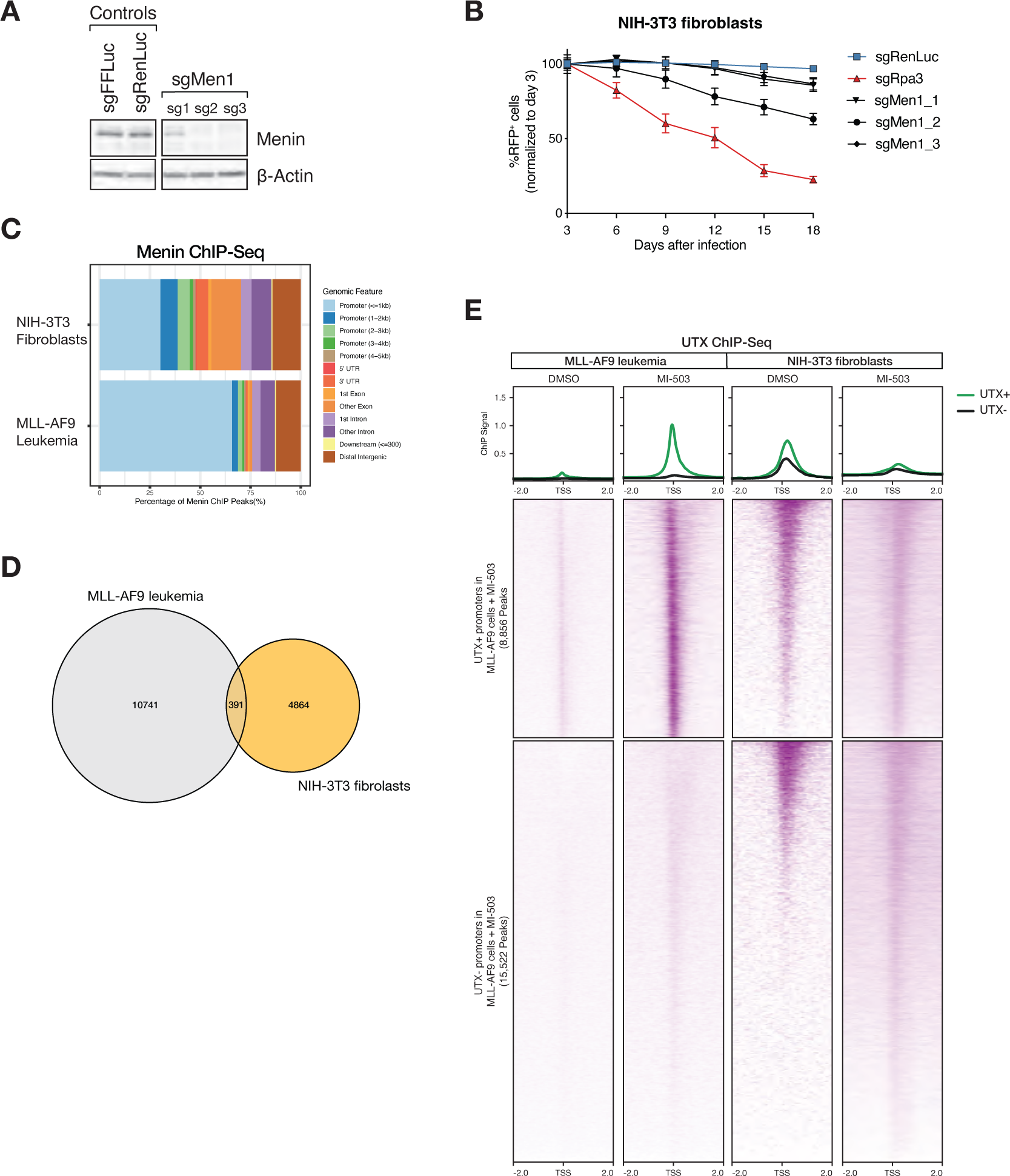
Menin-independent mammalian cells do not exhibit chromatin changes associated with the Menin-UTX molecular switch. **(A)** Immunoblot analysis of Cas9-expressing mouse fibroblast (NIH-3T3) cells expressing different controls and *Men1* sgRNAs. **(B)** Growth competition assay in mouse fibroblast cells. Graph shows the relative growth of cells infected with RFP-tagged sgRNAs measured by flow cytometry (mean±SEM, n=3 infection replicates). **(C)** Genomic distribution of Menin ChIP-Seq peaks from mouse fibroblasts (top) or MLL-AF9 leukemia cells (bottom). **(D)** Venn diagram comparing the number of Menin peaks detected in mouse MLL-AF9 leukemia cells (grey) vs. NIH-3T3 fibroblasts (yellow). **(E)** Heatmaps displaying UTX ChIP-Seq signals mapping to a 4-kb window around TSSs. Data is shown for mouse MLL-AF9 cells (left) and mouse fibroblasts (right) treated with vehicle (DMSO) or Menin-MLL inhibitor (MI-503) for 96 hours. Metagene plot represents the average ChIP-Seq signal for UTX at promoters that are enriched for UTX (green) or not (black) in mouse MLL-AF9 cells treated with MI-503.

**Supplementary Figure 8.**
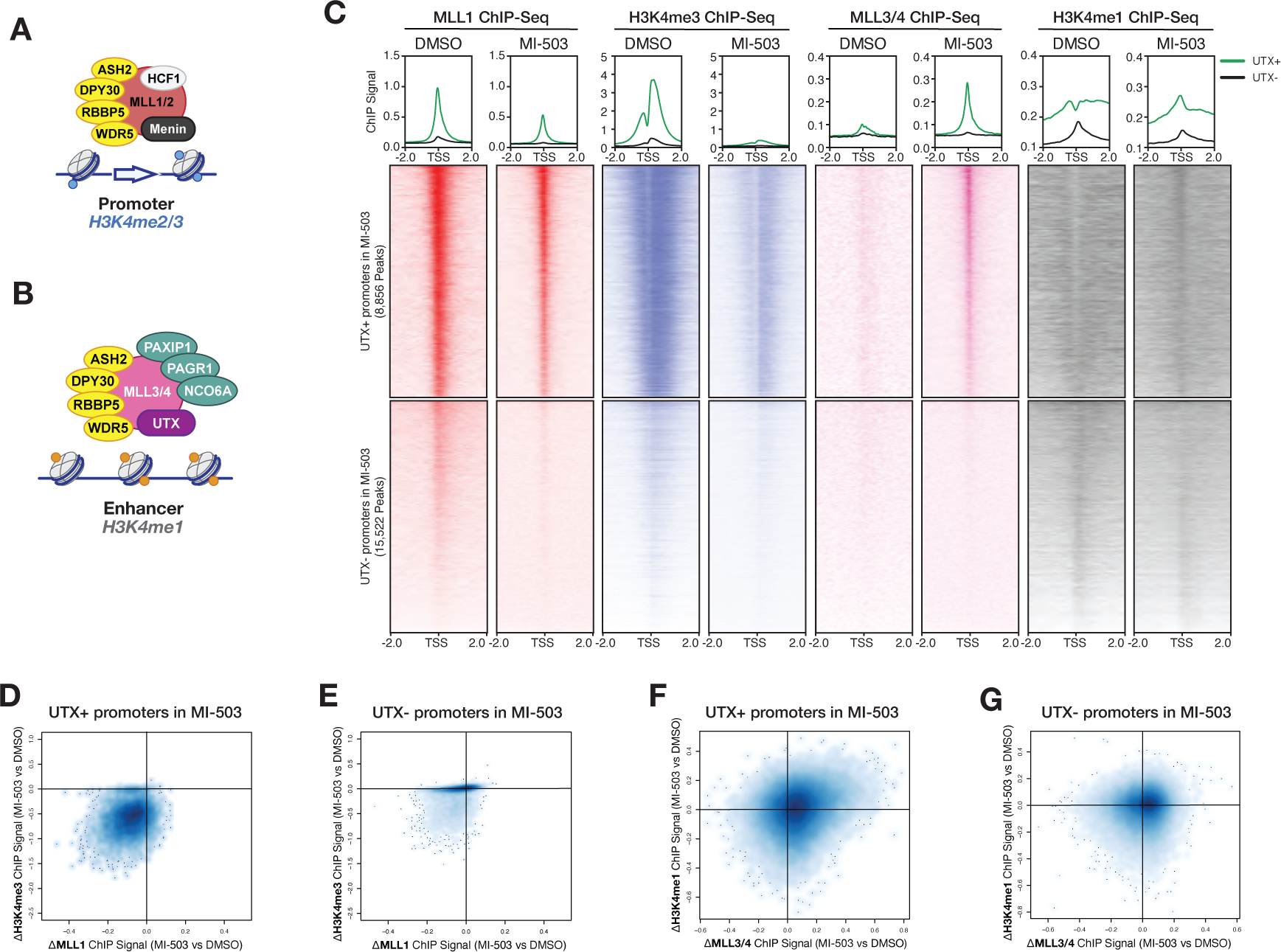
Disruption of Menin-MLL1 interaction leads to targeting of MLL3/4-UTX to sites normally bound by the MLL1-Menin complex. **(A)** Schematic representation of the mammalian MLL1 and MLL2 histone methyltransferase complexes. Highlighted in red is the enzymatic subunit of the complex (MLL1 and MLL2 are mutually exclusive). Yellow denotes shared subunits with the MLL3/4 complex. **(B)** Schematic representation of the mammalian MLL3 and MLL4 histone methyltransferase complexes. Highlighted in pink and purple are the enzymatic subunits of the complex (MLL3 and MLL4 are mutually exclusive). Yellow denotes shared subunits with the MLL1/2 complex. Green denotes non-enzymatic subunits of the MLL3/4 complex. **(C)** Heatmaps displaying MLL1 (red) and H3K4me3 (blue) ChIP-Seq signals mapping to a 4-kb window around TSSs. Data is shown for cells treated with vehicle (DMSO) or Menin-MLL inhibitor (MI-503) for 96 hours. Metagene plot represents the average ChIP-Seq signal for each protein at promoters that are UTX+ (green) or UTX- (black). Heatmaps displaying MLL3/4 (pink) and H3K4me1 (grey) ChIP-Seq signals mapping to a 4-kb window around TSSs. Data is shown for cells treated with vehicle (DMSO) or Menin-MLL inhibitor (MI-503) for 96 hours. Metagene plot represents the average ChIP-Seq signal for each protein at promoters that are UTX+ (green) or UTX- (black). **(D)** Density plot showing correlation between H3K4me3 and MLL1 ChIP-Seq signals (MI-503 vs DMSO). Signals correspond to summed signal ±2kb around TSSs that overlap with UTX peaks in MI-503 condition. **(E)** Density plot showing correlation between H3K4me3 and MLL1 ChIP-Seq signals (MI-503 vs DMSO). Signals correspond to summed signal ±2kb around TSSs that do not overlap with UTX peaks in MI-503 condition. **(F)** Density plot showing correlation between H3K4me1 and MLL3/4 ChIP-Seq signals (MI-503 vs DMSO). Signals correspond to summed signal ±2kb around TSSs that overlap with UTX peaks in MI-503 condition. **(G)** Density plot showing correlation between H3K4me1 and MLL3/4 ChIP-Seq signals (MI-503 vs DMSO). Signals correspond to summed signal ±2kb around TSSs that do not overlap with UTX peaks in MI-503 condition.

**Supplementary Figure 9.**
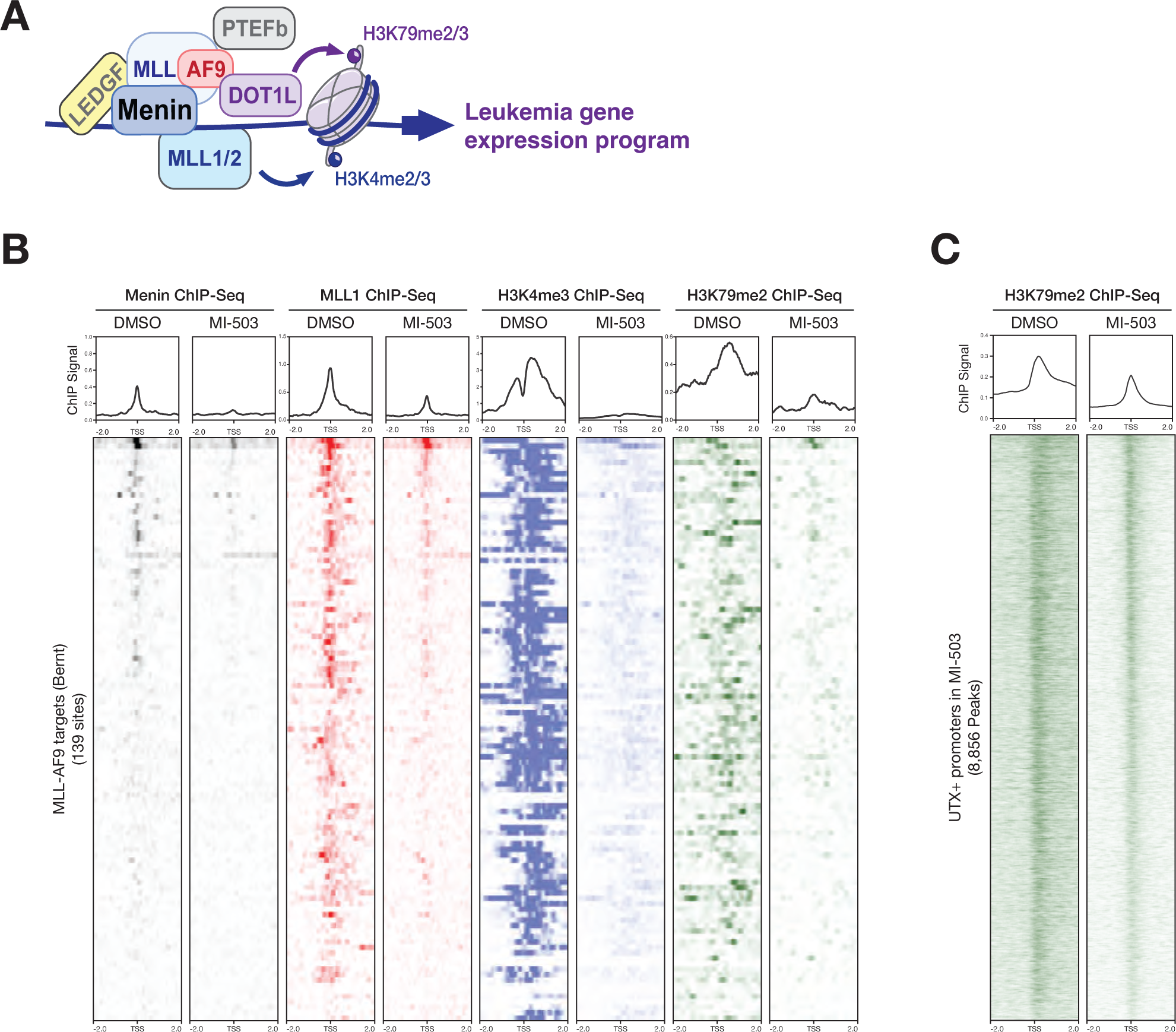
MLL1-Menin genomic targets are distinct from MLL-FP targets. **(A)** Schematic representation of ectopic chromatin complexes that are formed by MLL-fusion proteins (MLL-FPs). **(B)** Heatmaps displaying Menin (black), MLL1 (red), H3K4me3 (blue), and H3K79me2 (green) ChIP-Seq signals mapping to MLL-AF9 target loci (139 sites) from MLL-AF9 leukemia cells treated with vehicle (DMSO) or Menin-MLL inhibitor (MI-503) for 96 hours. Metagene plot represents the average ChIP-Seq signal for each protein at promoters. **(C)** Heatmaps displaying H3K79me2 ChIP-Seq signals mapping to a 4-kb window around TSSs in MLL-AF9 leukemia cells treated with vehicle (DMSO) or Menin-MLL inhibitor (MI-503) for 96 hours. Metaplot represents the average ChIP-Seq signal for H3K79me2 at promoters that are bound by UTX in MLL-AF9 cells treated with MI-503.

**Supplementary Figure 10.**
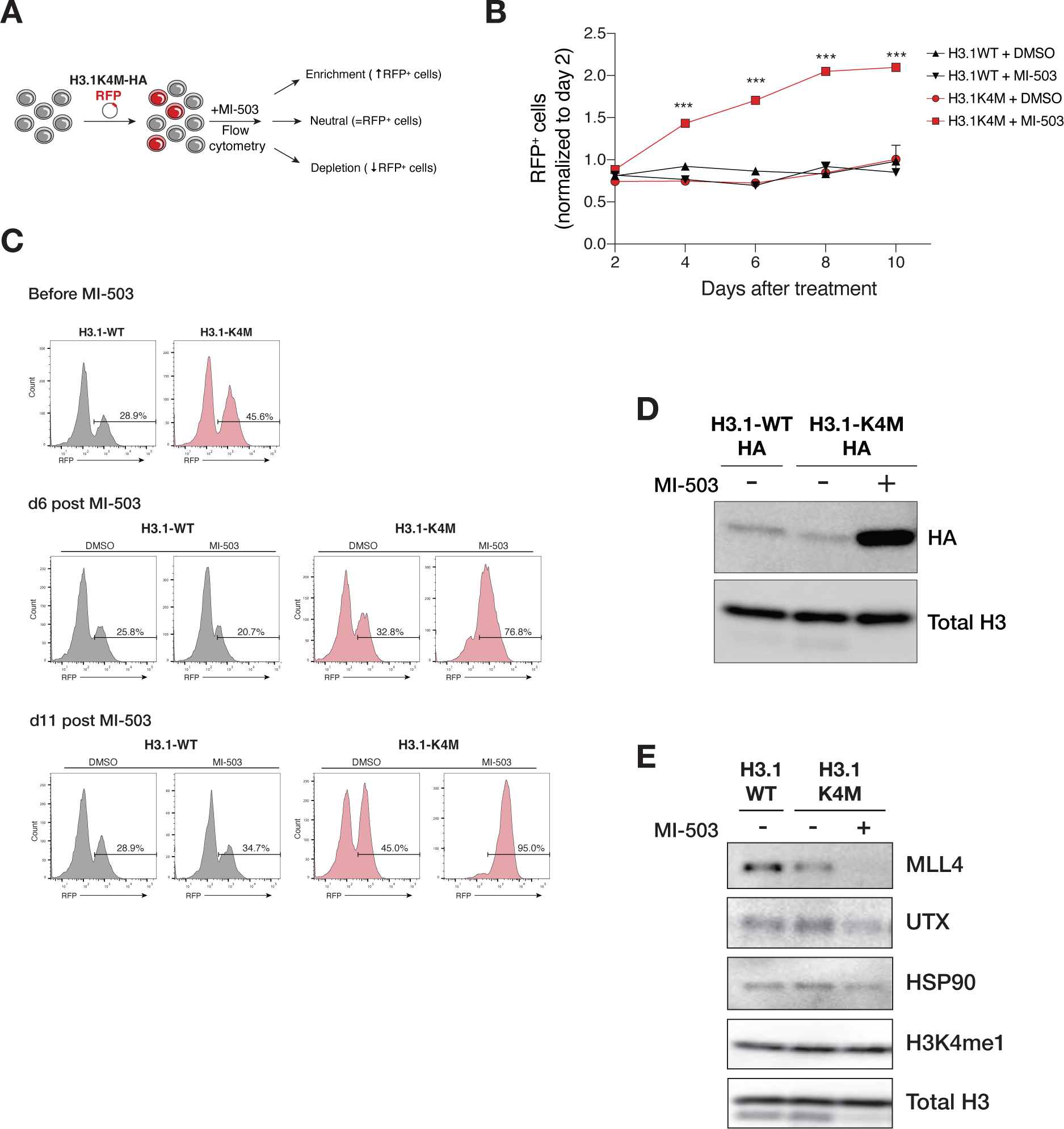
Destabilization of the MLL3/4-UTX complex is sufficient to blunt cellular responses to Menin-MLL inhibition. **(A)** Layout of growth competition assay to assess the effects of expressing a H3.1K4M oncohistone mutation in MLL-AF9 leukemia cells in response to Menin-MLL inhibition. **(B)** Relative percentage of RFP+ transgene (H3.1WT or H3.1K4M) cells over time after transduction of mouse MLL-AF9 cells treated with vehicle (DMSO) or Menin-MLL inhibitor (MI-503) (mean±SEM, n=3 replicates, *P*-value calculated by Student’s t-test, *P* < 0.001 = ***). **(C)** Expression of H3.1WT-RFP or H3.1K4M-RFP in mouse MLL-AF9 leukemia cells treated with DMSO or MI-503 was monitored by tracking RFP+ cells by flow cytometry over time. **(D)** Immunoblot analysis of HA-tag from MLL-AF9 leukemia cells expressing H3.1WT-HA or H3.1K4M-HA and treated with vehicle (DMSO) or Menin-MLL inhibitor (MI-503) for 6 days. **(E)** Immunoblot analysis of MLL4, UTX, HSP90 (loading control), H3K4me1, and total H3 (loading control) from MLL-AF9 leukemia cells expressing H3.1WT or H3.1K4M and treated with DMSO or MI-503 for 6 days.

**Supplementary Figure 11.**
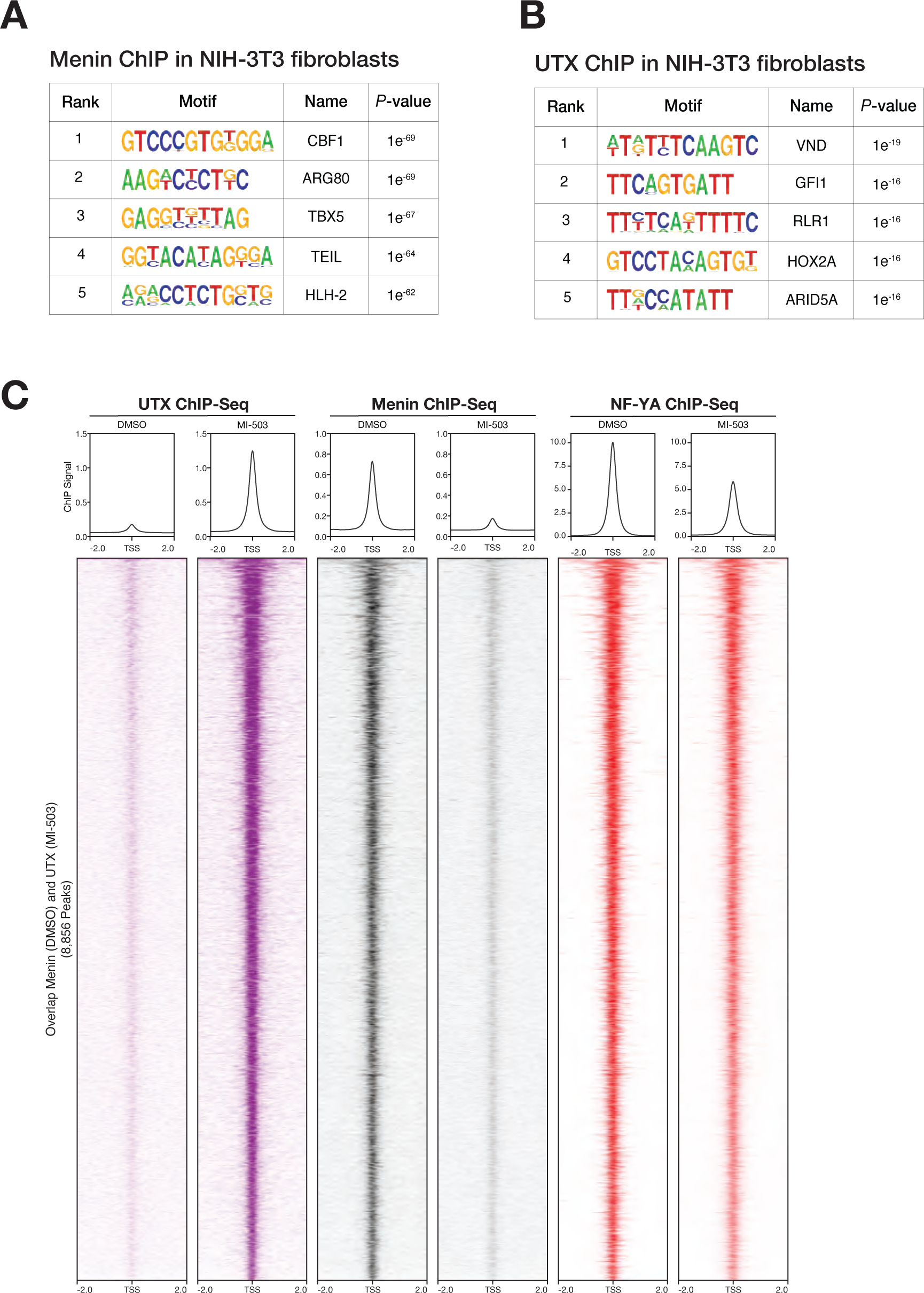
NF-Y restricts chromatin occupancy of UTX at promoter regions. **(A)** Motif analysis of Menin ChIP-Seq peaks detected in NIH-3T3 mouse fibroblasts. **(B)** Motif analysis of UTX ChIP-Seq peaks detected in NIH-3T3 mouse fibroblasts. **(C)** Heatmaps displaying UTX (purple), Menin (black), and NF-YA (red) ChIP-Seq signals mapping to a 4-kb window around TSSs. Data is shown for vehicle (DMSO) and Menin-MLL inhibitor (MI-503)-treated cells for 96 hours. Metagene plot represents the average ChIP-Seq signal for each protein at promoters.

**Supplementary Figure 12.**
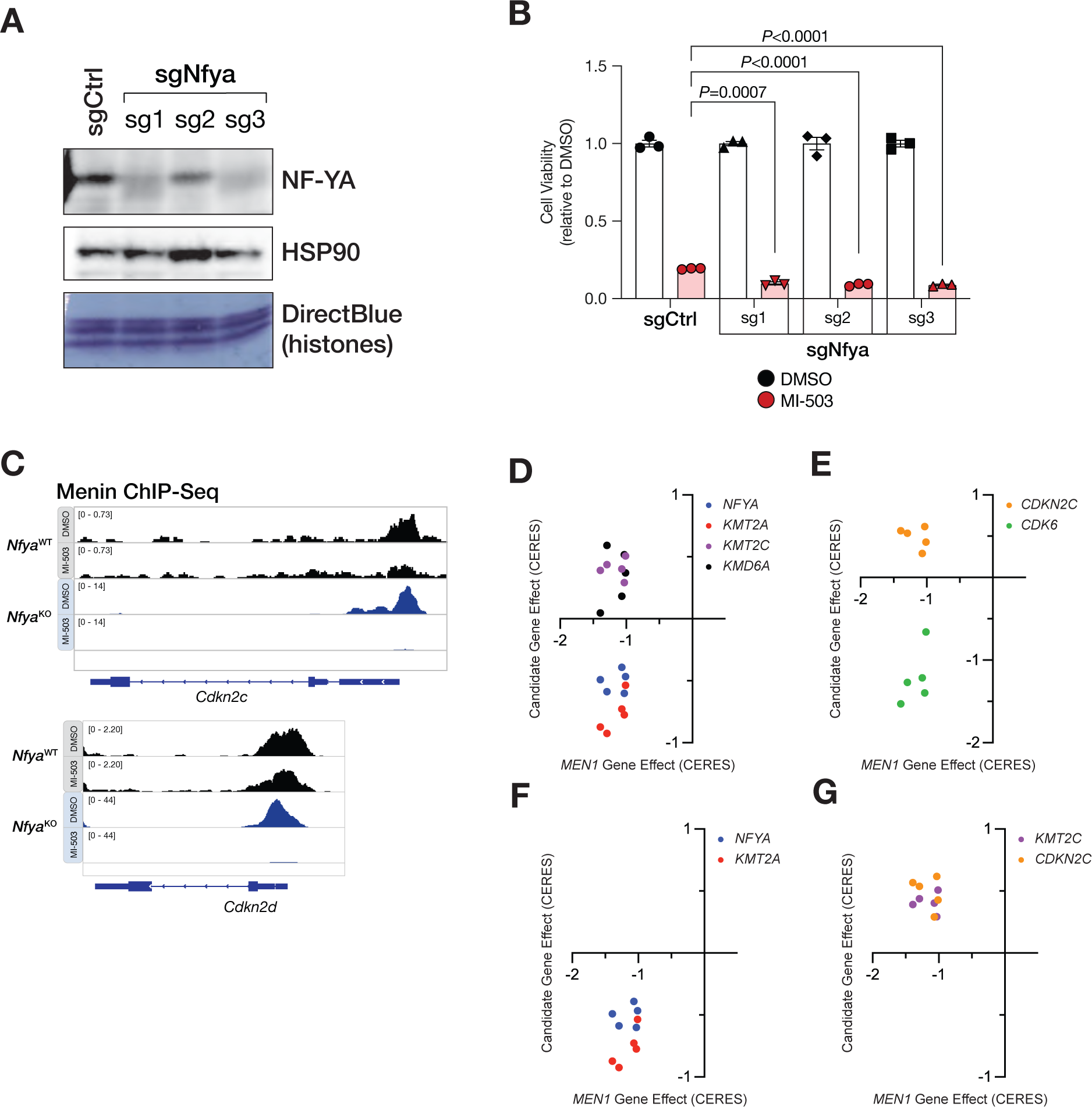
NF-Y facilitates promoter regulation by Menin. **(A)** Immunoblot analysis of NF-YA and HSP90 proteins (loading control) in mouse MLL-AF9 leukemia cells transduced with a control sgRNA (sgCtrl) or three independent Nfya-targeting sgRNAs. Direct blue staining of histones (bottom) serves as an additional loading control. **(B)** Viability assay from cells treated with vehicle (DMSO, black) or Menin-MLL inhibitor (MI-503, red) for 6 days (mean±SEM, n=3 infection replicates, *P*-value calculated by Student’s t-test). sgCtrl, control sgRNA targeting a non-genic region on chromosome 8. **(C)** Genome browser representation of Menin ChIP-Seq normalized reads (average RPKM) for representative loci bound by Menin in *Nfya*^WT^ (black) or *Nfya*^KO^ (blue) MLL-AF9 leukemia cells treated with vehicle (DMSO) and Menin-MLL inhibitor (MI-503) for 96 hours. **(D)** Co-essentiality scatter plot showing relationship between MEN1 loss-of-function and dependency to *NFYA*, *MLL1* (*KMT2A*), *MLL3* (*KMT2C*), and *UTX* (*KDM6A*). **(E)** Co-essentiality scatter plot showing relationship between MEN1 loss-of-function and dependency to *CDKN2C* and *CDK6*. **(F)** Co-essentiality scatter plot showing relationship between MEN1 loss-of-function and dependency to *NFYA* and *MLL1* (*KMT2A*) **(G)** Co-essentiality scatter plot showing relationship between *MEN1* loss-of-function and dependency to *MLL3* (*KMT2C*) and *CDKN2C*.

**Supplementary Figure 13.**
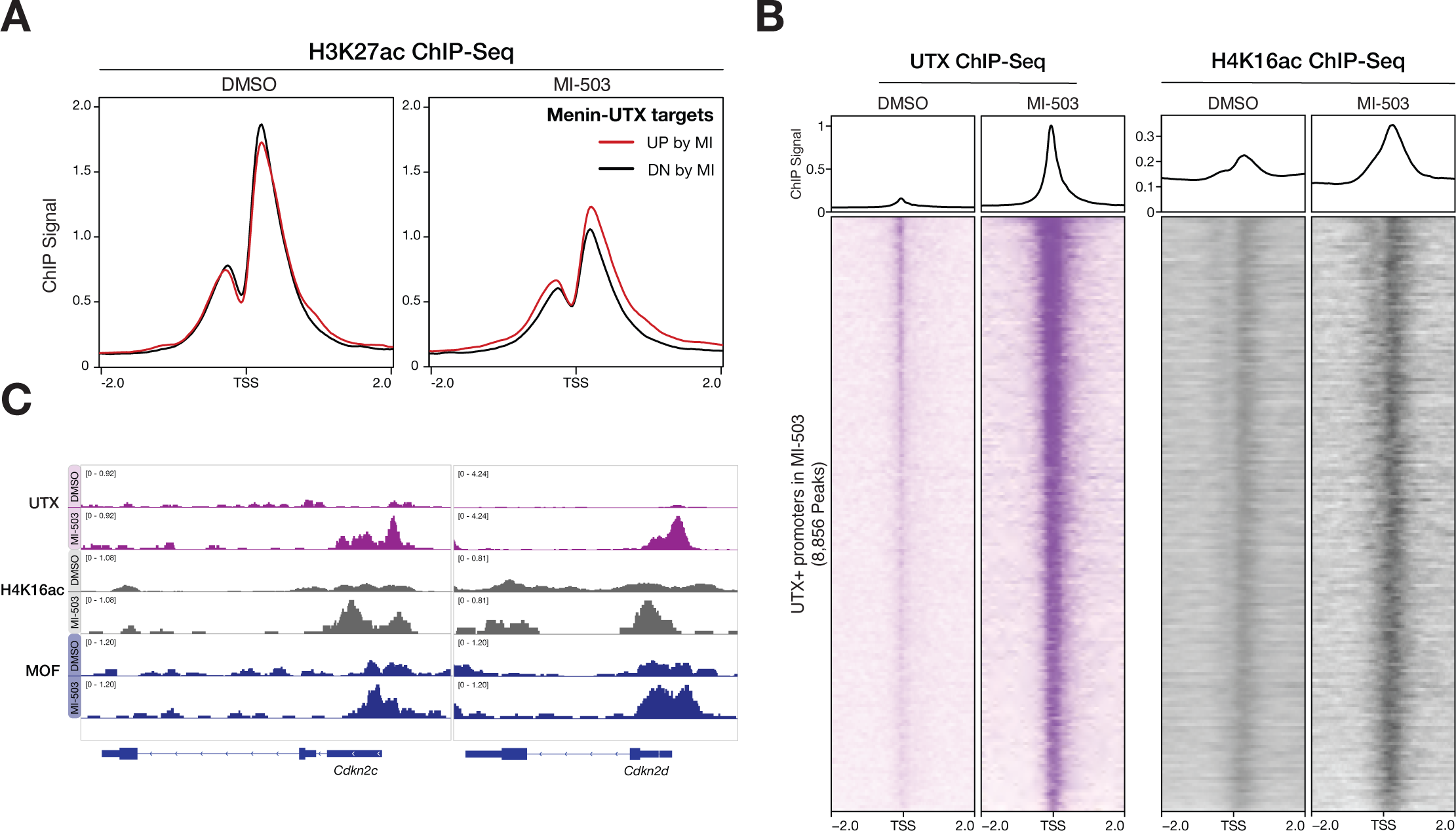
Menin-UTX molecular switch coincides with local changes in histone acetylation. **(A)** Average ChIP-Seq signal for H3K27ac at transcription start sites (TSS) ±2kb of the Menin-UTX target genes. Signals corresponding to genes that are upregulated (red) or down-regulated (black) when cells are treated with Menin-MLL inhibitor (MI-503) for 96 hours. RPM, reads per million. **(B)** Heatmaps displaying UTX (purple) or H4K16ac (black) ChIP-Seq signals mapping to a 4-kb window around TSSs in MLL-AF9 leukemia cells treated with vehicle (DMSO) or Menin-MLL inhibitor (MI-503) for 96 hours. Metagene plot represents the average ChIP-Seq signal for each protein at promoters that are bound by UTX in mouse MLL-AF9 leukemia cells treated with MI-503 for 96 hours. **(C)** Genome browser representation of ChIP-Seq normalized reads (average RPKM) for UTX (purple), H4K16ac (grey), and MOF (blue) at *Cdkn2c* and *Cdkn2d* loci from MLL-AF9 cells treated with DMSO or MI-503 for 96 hours.

**Supplementary Figure 14.**
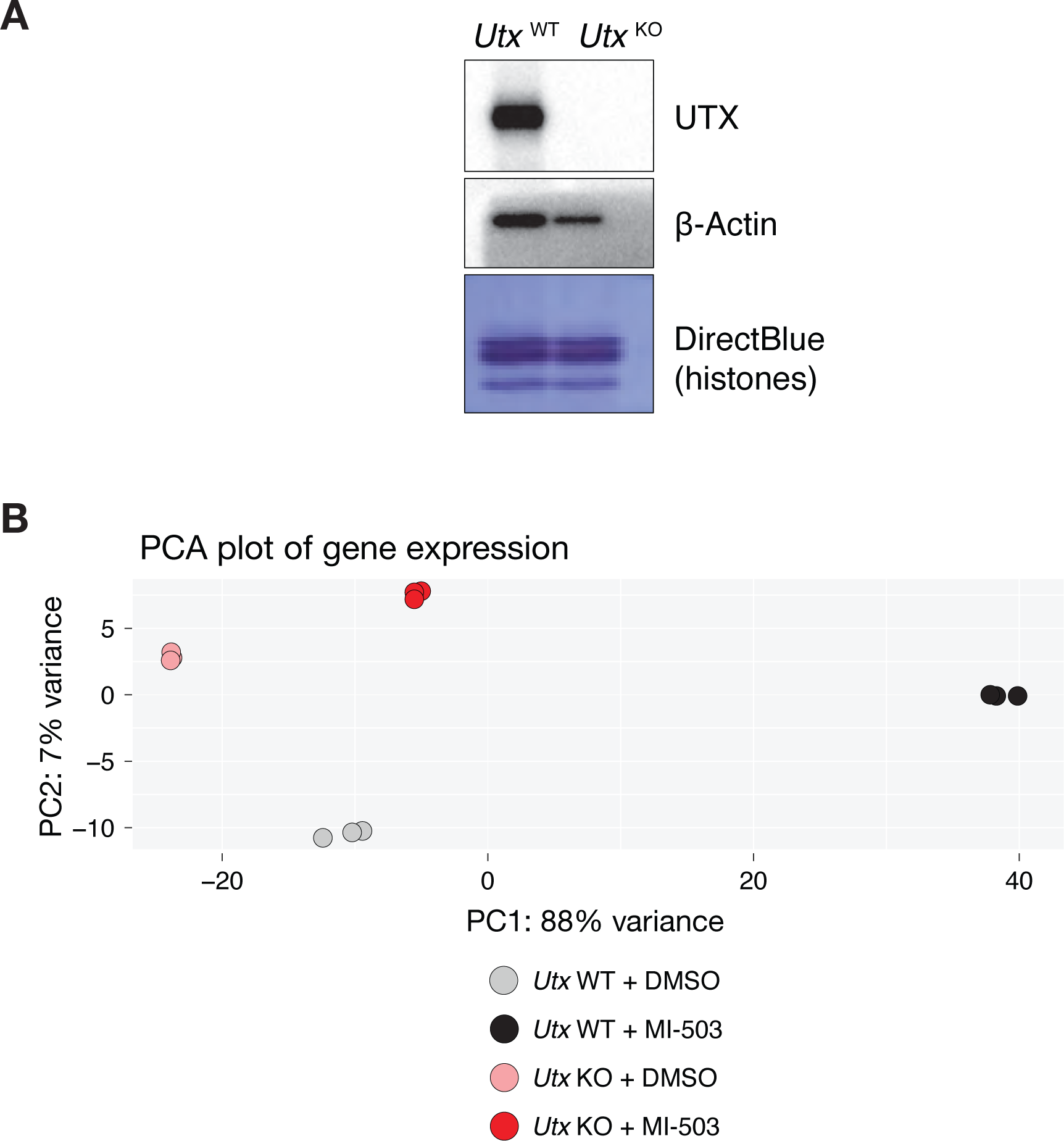
Transcriptional assessment of UtxKO MLL-AF9 leukemia cells. **(A)** Immunoblot analysis of UTX and β-actin (loading control) from mouse UtxWT and UtxKO MLL-AF9 leukemia cells. **(B)** Principal component analysis (PCA) using RNA-Seq gene expression data from *Utx*^WT^ and *Utx*^KO^ cells treated with vehicle (DMSO) or Menin-MLL inhibitor (MI-503) for 96 hours.

**Supplementary Figure 15.**
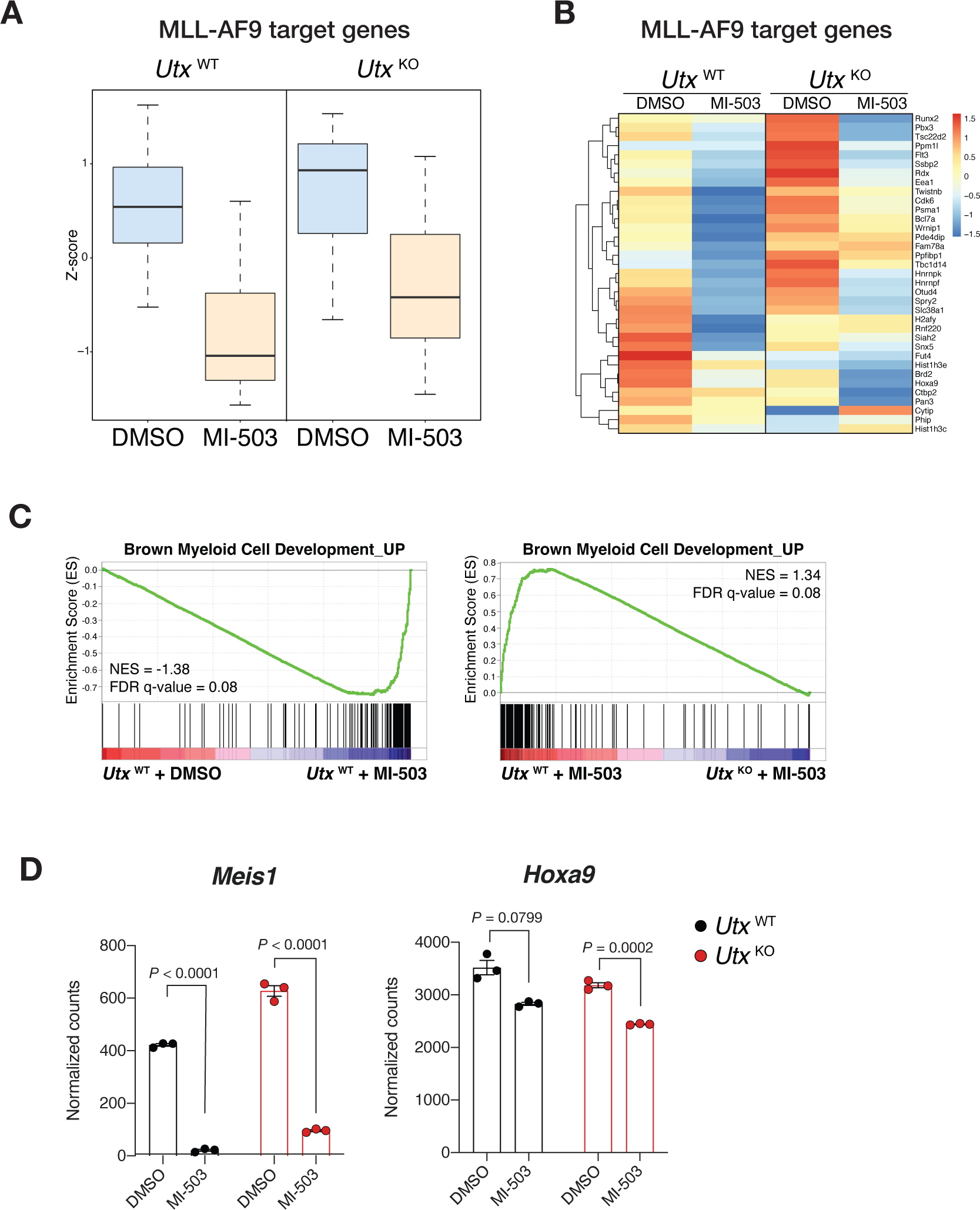
MLL-AF9 target genes are not regulated by the Menin-UTX molecular switch. **(A)** Boxplot showing expression levels of MLL-AF9 target genes, which are suppressed by the Menin-MLL inhibitor (MI-503) treatment for 96 hours. Expression levels are shown for *Utx*^WT^ (left) and *Utx*^KO^ (right) mouse leukemia cells. **(B)** Heatmap of Z-scores for expression of MLL-AF9 target genes from *Utx*^WT^ and *Utx*^KO^ leukemia cells treated with DMSO or MI-503 for 96 hours. **(C)** GSEA plots showing changes in regulation of myeloid cell differentiation in genes induced by MI-503 for 96 hours. FDR, false discovery rate; NES, normalized enrichment score. **(D)** *Meis1* (left) and *Hoxa9* (right) expression (mean normalized read counts) from *Utx*^WT^ and *Utx*^KO^ leukemia cells treated with vehicle (DMSO) or Menin-MLL (MI-503) for 96 hours (mean±SEM, n=3 replicates, *P*-value calculated by Student’s t-test).

**Supplementary Figure 16.**
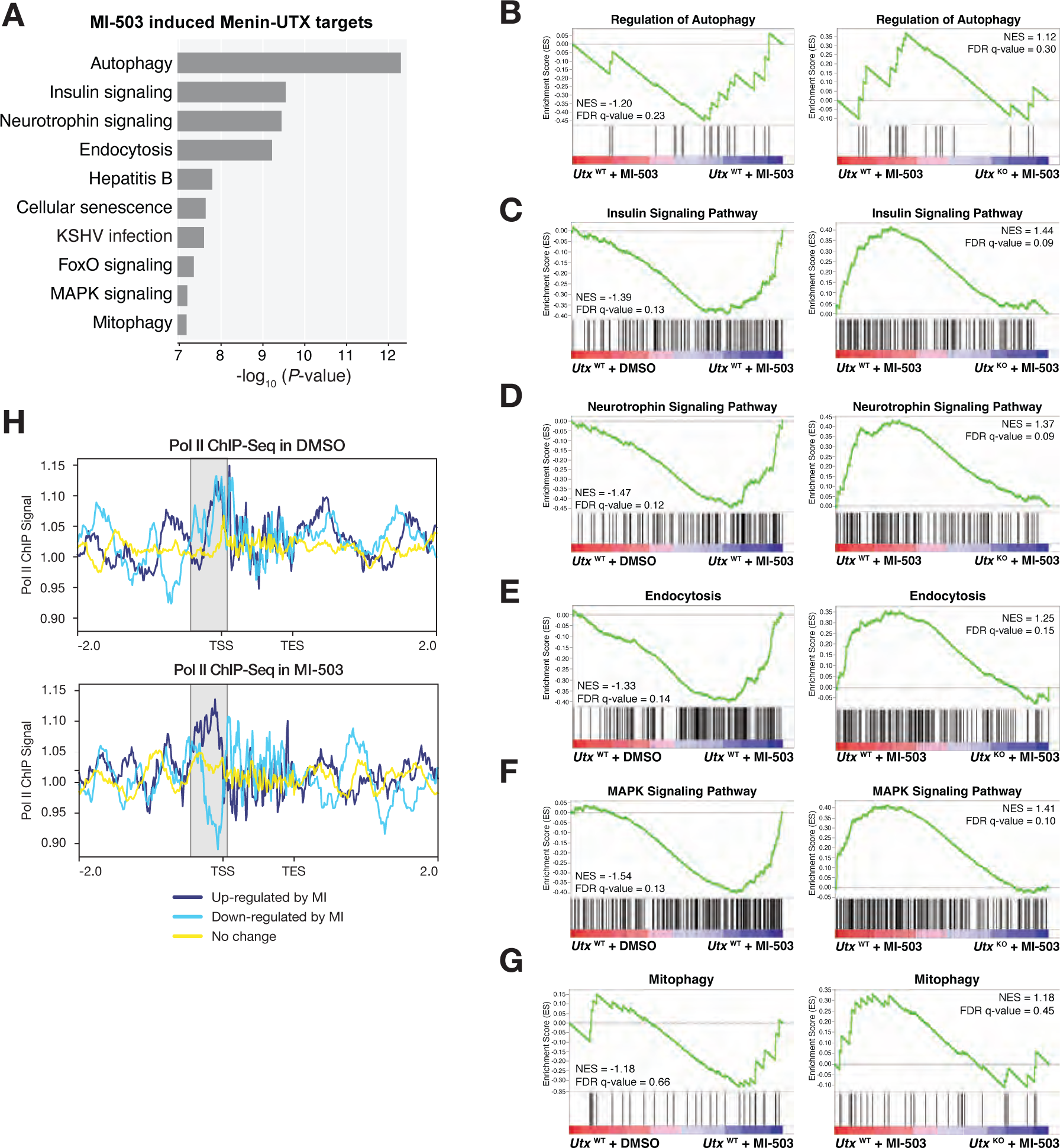
Gene sets and pathways regulated by Menin-UTX molecular switch. **(A)** Gene ontology (GO) analysis of Menin-UTX targets using expression data from mouse MLL-AF9 leukemia cells treated with vehicle (DMSO) or Menin-MLL (MI-503) for 96 hours. Top 10 GO categories sorted based on statistical significance are shown. **(B)** GSEA plots showing changes in regulation of autophagy genes induced by MI-503 treatment for 96 hours. FDR, false discovery rate; NES, normalized enrichment score. **(C)** GSEA plots showing changes in insulin signaling pathway genes induced by MI-503 treatment for 96 hours. **(D)** GSEA plots showing changes in neurotrophin signaling pathway genes induced by MI-503 treatment for 96 hours. **(E)** GSEA plots showing changes in endocytosis genes induced by MI-503 treatment for 96 hours. **(F)** GSEA plots showing changes in MAPK signaling pathway genes induced by MI-503 treatment for 96 hours. **(G)** GSEA plots showing changes in mitophagy genes induced by MI-503 treatment for 96 hours. **(H)** RNA Pol II ChIP-Seq signal determined for senescence and cell-cycle arrest-associated genes that are Menin-UTX targets.

**Supplementary Figure 17.**
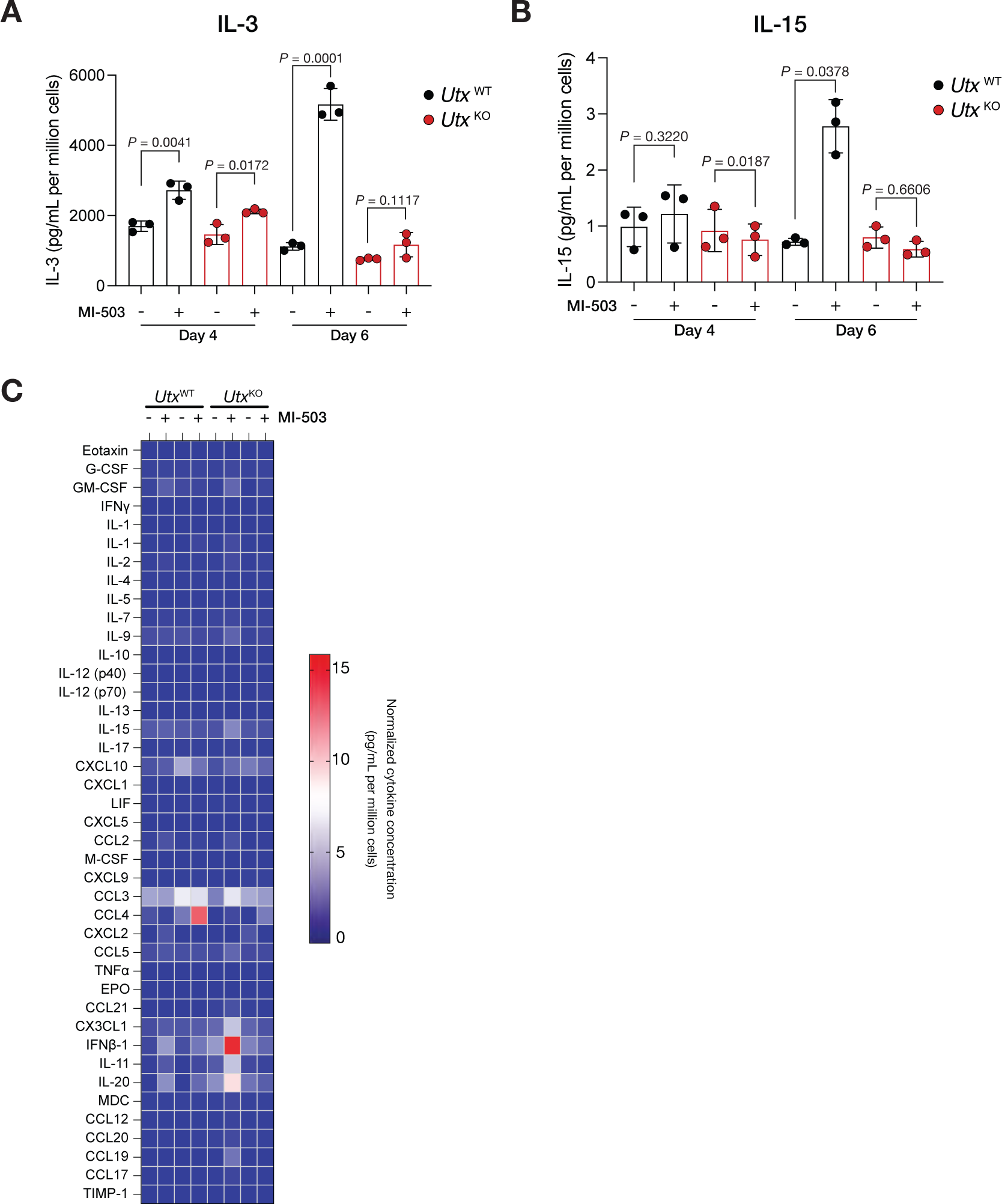
MLL3/4-UTX-dependent induction of SASP cytokines upon Menin-MLL inhibition. **(A)** Secreted levels of IL-3 in mouse UtxWT and UtxKO MLL-AF9 leukemia cells treated with Menin inhibitor (MI-503) or vehicle (DMSO) after four or six days. Data are quantified as pg/mL of secreted cytokine per million cells (mean±SEM, n=3 replicates, *P*-value calculated by Student’s t-test). **(B)** Secreted levels of IL-15 in *Utx*^WT^ and *Utx*^KO^ MLL-AF9 leukemia cells treated with Menin inhibitor (MI-503) or vehicle (DMSO) after four or six days. Data are quantified as pg/mL of secreted cytokine per million cells (mean±SEM, n=3 replicates, *P*-value calculated by Student’s t-test). **(C)** Heatmap of cytokine array results from *Utx*^WT^ and *Utx*^KO^ MLL-AF9 leukemia cells treated with Menin inhibitor (MI-503) or vehicle (DMSO) after four or six days. Data are quantified as pg/mL of secreted cytokine per million cells and represent the mean of three biological replicates. Note that the very high levels of IFNβ-1 secretion relative to the rest of the cytokines measured affect the dynamic range of data visualization.

**Supplementary Figure 18.**
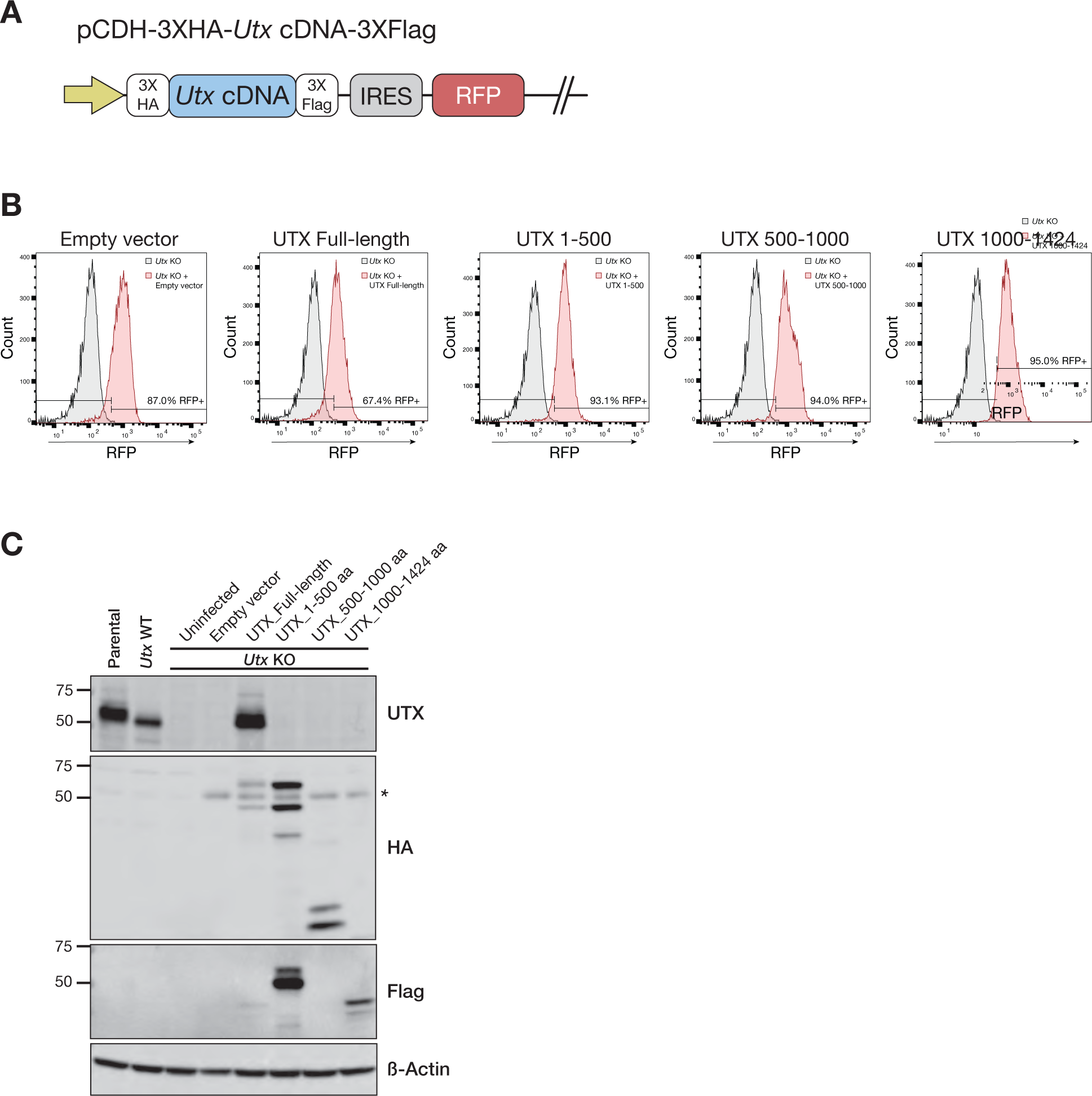
Expression of UTX truncations in MLL-AF9 *Utx*^KO^ leukemia cells. **(A)** Representation of the lentiviral vector used to stably co-express N-terminal HA-/C-terminal-Flag-tagged *Utx* cDNAs and the fluorescent protein RFP. **(B)** Flow cytometry plots showing RFP expression of the different mouse MLL-AF9 leukemia cell lines generated with the *Utx* truncations. **(C)** Immunoblotting validation of expression of the different *Utx* truncations in mouse MLL-AF9 leukemia cells.

**Supplementary Figure 19.**
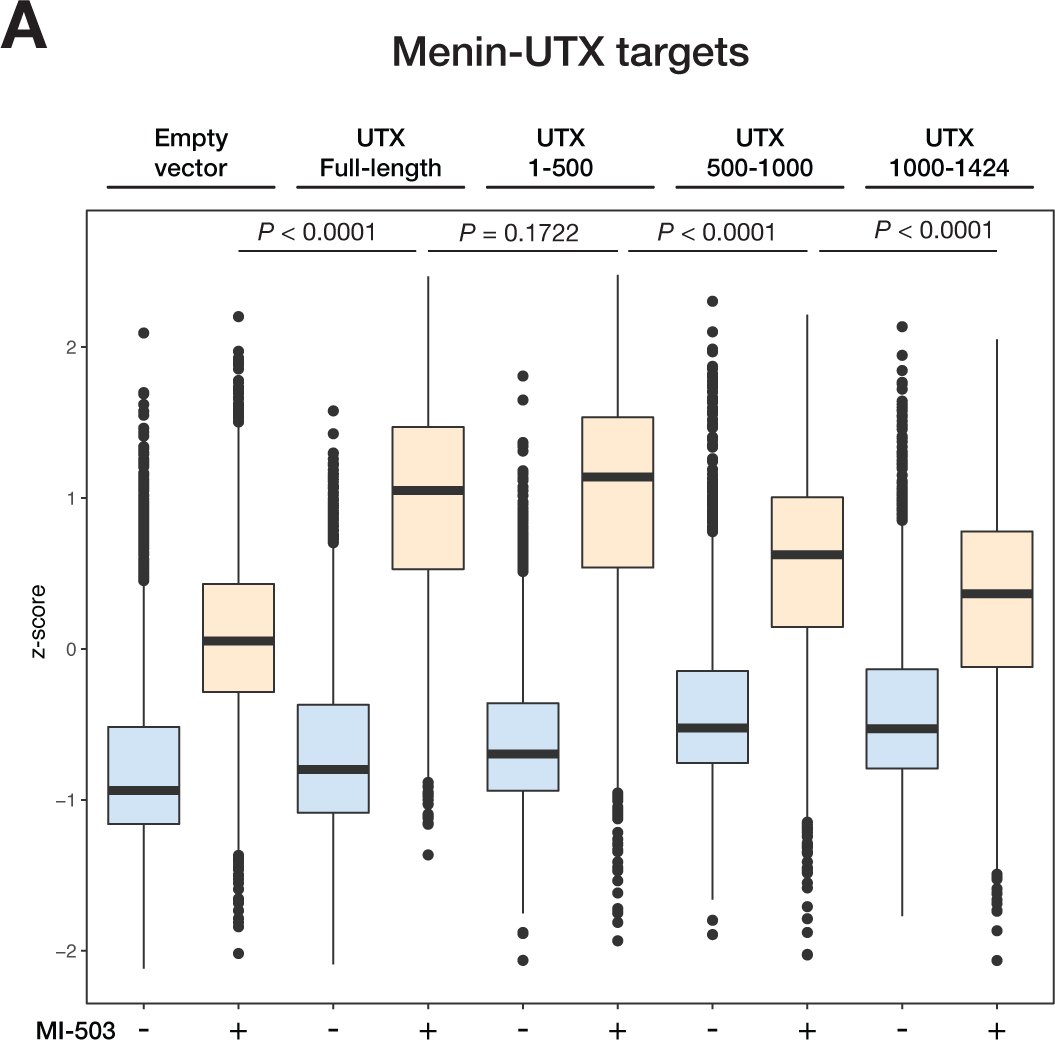
Transcriptional characterization of the different *Utx* truncations in mouse MLL-AF9 leukemia cells. **(A)** Boxplot showing expression levels of Menin-UTX target genes in mouse MLL-AF9 leukemia cells treated with vehicle (DMSO) or Menin-MLL inhibitor (MI-503) for 96 hours.

**Supplementary Figure 20.**
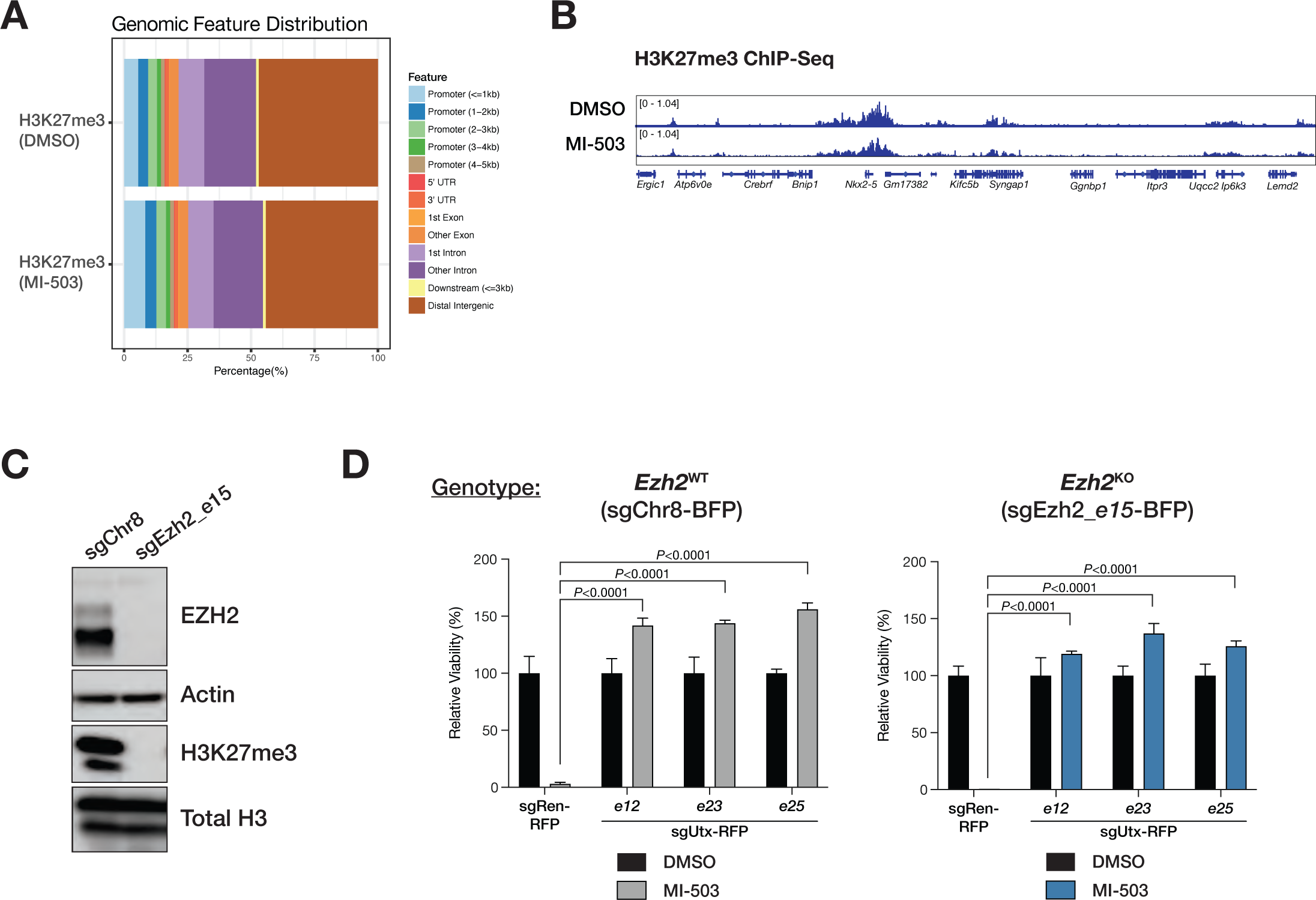
UTX-dependent phenotypes are not associated with global changes in H3K27 methylation. **(A)** Genomic distribution of H3K27me3 ChIP-Seq peaks from mouse leukemia cells treated with vehicle (DMSO) or Menin-MLL inhibitor (MI-503) for 96 hours. **(B)** Genome browser representation of ChIP-Seq normalized reads (average RPKM) for H3K27me3 from MLL-AF9 cells treated with DMSO or MI-503 for 96 hours. **(C)** Immunoblot analysis of EZH2, β-actin (loading control), H3K27me3, and total H3 (loading control) from *Ezh2*^WT^ and *Ezh2*^KO^ mouse MLL-AF9 leukemia cells. **(D)** Viability assay from mouse *Ezh2*^WT^ (left) and *Ezh2*^KO^ (right) MLL-AF9 leukemia cells treated with vehicle (DMSO, black) or Menin-MLL inhibitor (MI-503, grey or blue) for 96 hours (mean±SEM, n=3 infection replicates, *P*-value calculated by Student’s t-test). sgCtrl, control sgRNA targeting a non-genic region on chromosome 8.

**Supplementary Figure 21.**
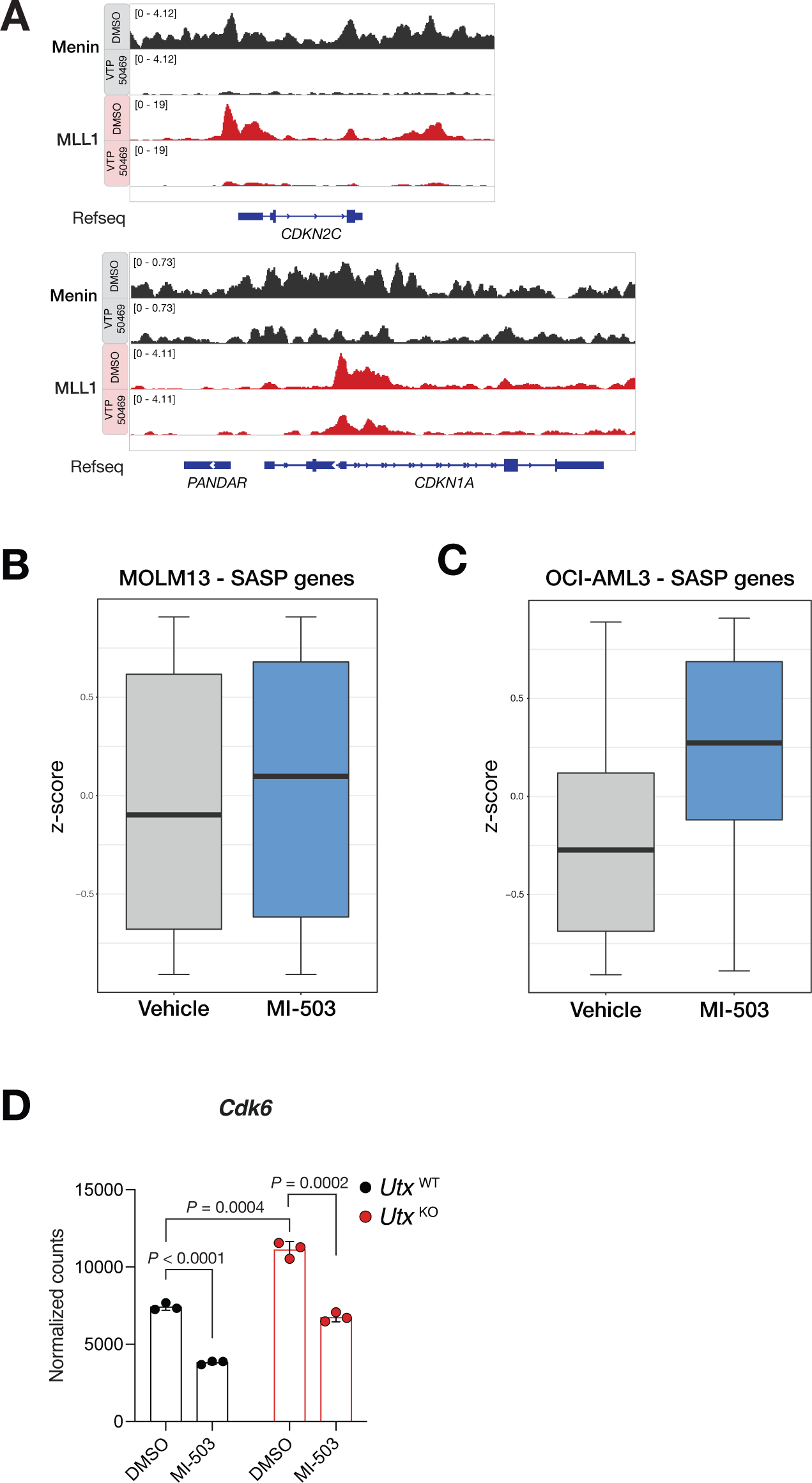
Induction of the Menin-UTX molecular switch upon Menin-MLL1 inhibition in human leukemia cells. **(A)** Genome browser representation of ChIP-Seq normalized reads (average RPKM) for Menin (black) or MLL1 (red) at *CDKN2C* and *CDKN1A* loci from MOLM13 human leukemia cells treated with vehicle (DMSO) or Menin-MLL inhibitor (VTP-50469) for 7 days. (B) Boxplot showing expression levels of SASP genes in MOLM13 human leukemia cells treated with DMSO or MI-503 for 96 hours. **(C)** Boxplot showing expression levels of SASP genes in OCI-AML3 human leukemia cells treated with DMSO or MI-503 for 96 hours. **(D)** *Cdk6* expression (mean normalized read counts) from *Utx*^WT^ (black) and *Utx*^KO^ (red) leukemia cells treated with DMSO or MI-503 for 96 hours (mean±SEM, n=3 replicates, *P*-value calculated by Student’s t-test).

**Supplementary Figure 22.**
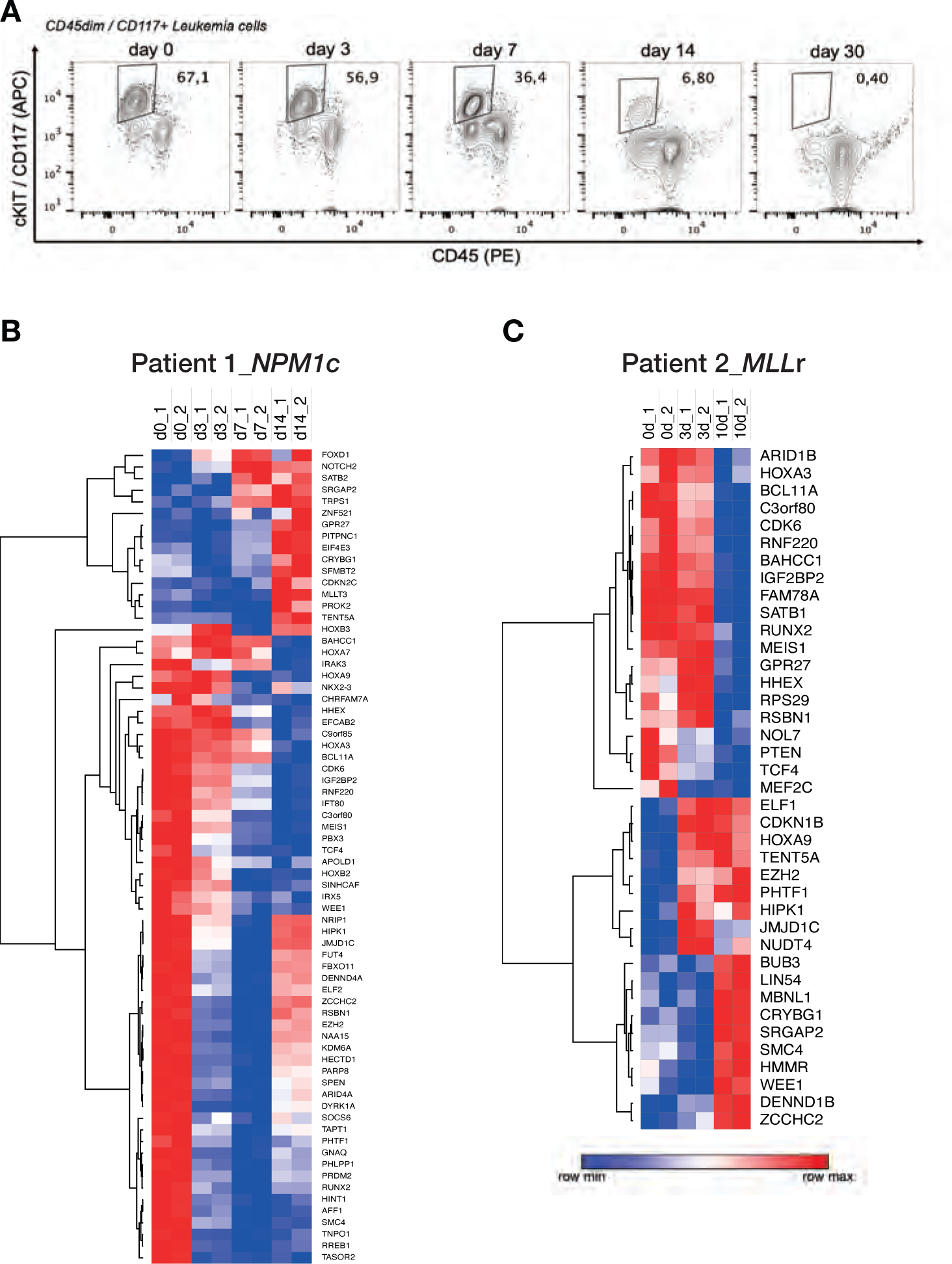
Characterization of primary human AML samples from patients participating in the Syndax Phase I Menin-MLL inhibitor trial. **(A)** Immunophenotyping of primary human AML cells isolated from Patient 1 (*NPM1c* mutant AML). **(B)** Heatmap representation of Z-scores from longitudinal gene expression analysis of AML cells derived from Patient 1 (*NPM1c* mutant AML) treated with SNDX-5613. **(C)** Heatmap representation of Z-scores from longitudinal gene expression analysis of AML cells derived from Patient 2 (*MLL*-rearranged AML) treated with SNDX-5613.

**Supplementary Figure 23.**
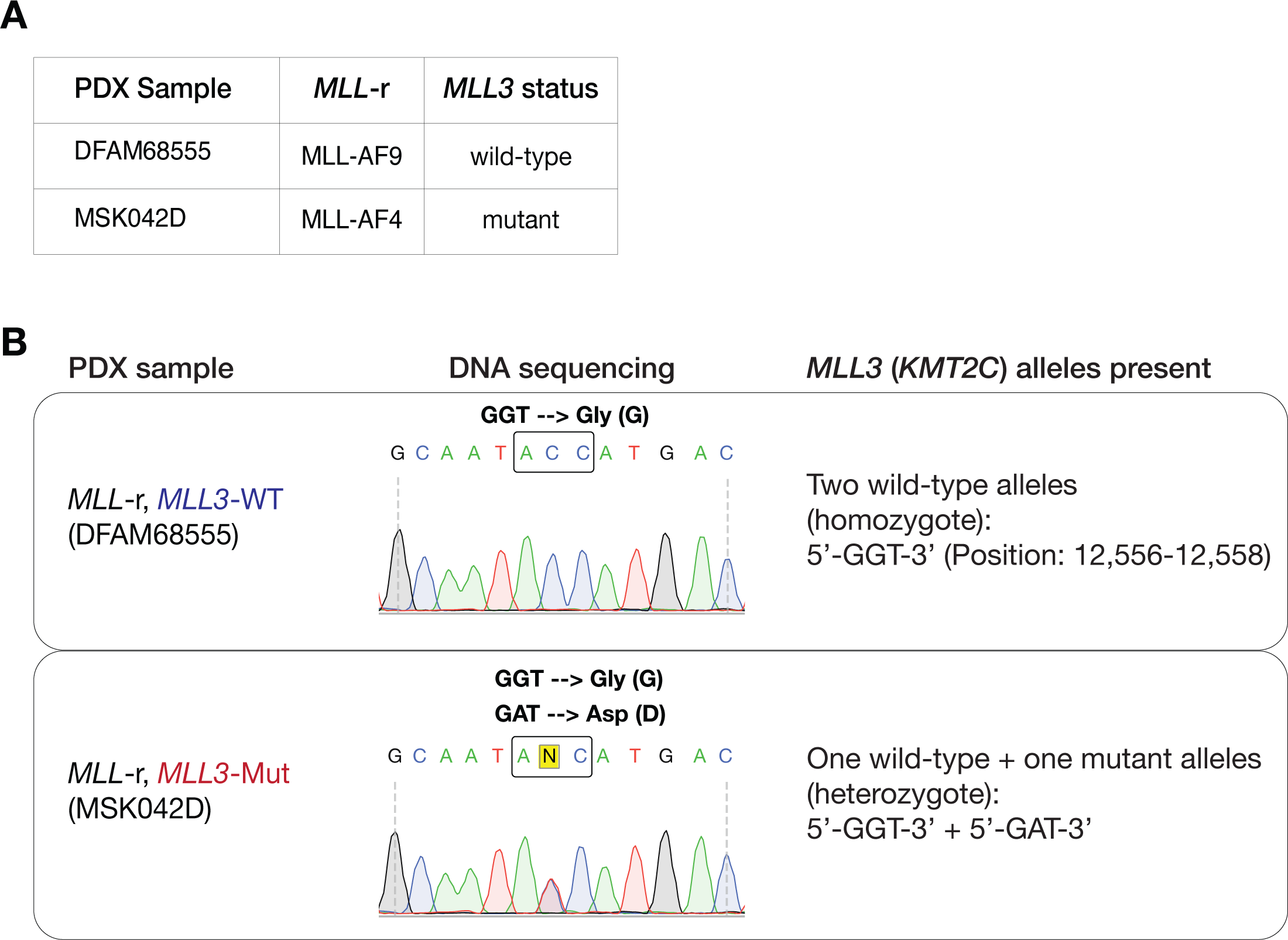
Modeling of Menin-MLL inhibitor response in patient-derived xenografts. **(A)** Characteristics of *MLL*-r AML PDX samples analyzed in this study. **(B)** Targeted sequencing validation of MLL3 status in PDX samples.

**Supplementary Figure 24.**
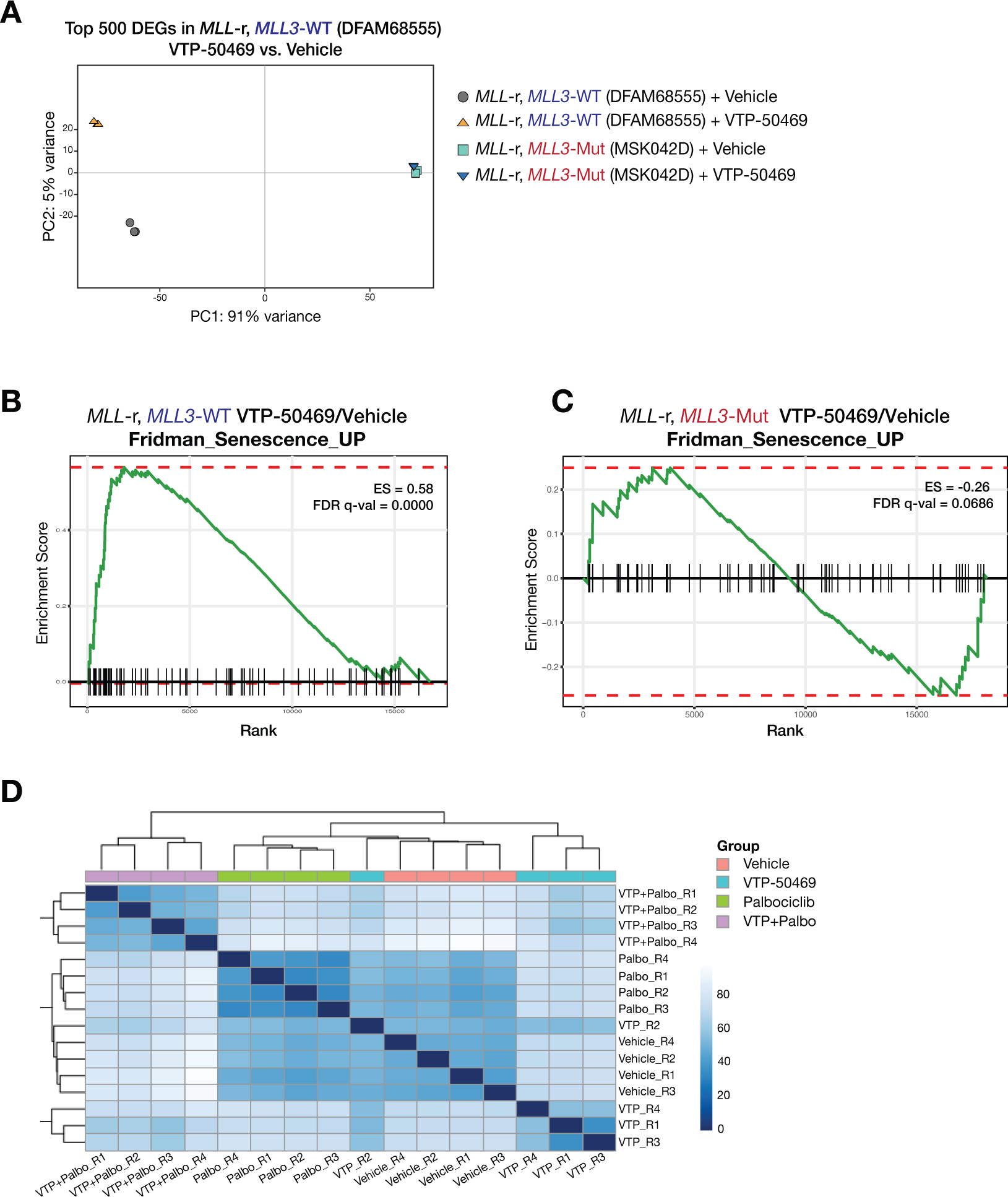
Transcriptional characterization of Menin-MLL inhibitor response in patient-derived xenografts. **(A)** PCA using gene expression data obtained from *MLL3*-WT (DFAM68555) and *MLL3*-Mutant (MSK042D) PDX samples treated with vehicle (DMSO) or Menin-MLL inhibitor (VTP-50469). Analysis was performed using the top 500 differentially-expressed genes upon treatment of *MLL3*-WT PDXs with VTP-50469. Note that *MLL3*-Mutant PDX samples are virtually overlapping independent of the condition, suggesting that the induction of the Menin-UTX molecular switch is impaired in the context of MLL3 mutation. **(B)** GSEA showing that Menin-MLL inhibition using VTP-50469 leads to induction of cellular senescence in *MLL3*-WT PDX. **(C)** GSEA showing that Menin-MLL inhibition using VTP-50469 fails to induce cellular senescence in MLL3-Mutant PDX. **(D)** Distance matrix for expression of all genes from RNA-Seq data obtained from MLL3-Mutant (MSK042D) PDX samples treated with DMSO, VTP-50469, Palbociclib, or a combination of the two inhibitors.

